# Longitudinal quantitative MRI of whole human brains during formaldehyde immersion, PBS wash-out, and tissue volume change

**DOI:** 10.64898/2026.01.31.702882

**Authors:** Francisco J. Fritz, Tobias Streubel, Laurin Mordhorst, Nina Lüthi, Luke J. Edwards, Herbert Mushumba, Klaus Püschel, Nikolaus Weiskopf, Evgeniya Kirilina, Siawoosh Mohammadi

**Affiliations:** Department of Systems Neurosciences, University Medical Center Hamburg-Eppendorf, Hamburg, Germany; Department of Legal Medicine, University Medical Center Hamburg-Eppendorf, Hamburg, Germany; Department of Neurophysics, Max Planck Institute for Human Cognitive and Brain Sciences, Leipzig, Germany; Felix Bloch Institute for Solid State Physics, Faculty of Physics and Earth System Sciences, Leipzig University, Leipzig, Germany; Wellcome Centre for Human Neuroimaging, UCL Queen Square Institute of Neurology, University College London, London, UK; Department of Neuroradiology, University Hospital Schleswig-Holstein and University of Lübeck, Lübeck, Germany; MR Physics Group, Max Planck Institute for Human Development, Berlin, Germany; Department of Cognitive Neuroscience, Faculty of Psychology and Neuroscience, Maastricht University, the Netherlands

**Keywords:** Quantitative MRI, MPM, ex vivo whole human brain, fixed tissue, longitudinal study, shrinkage

## Abstract

*Postmortem* MRI links *in vivo* MRI contrast to microstructure measured with *ex vivo* histology, yet fixation systematically alters relevant physical MRI parameters. We quantified these effects by longitudinally characterizing multi-parametric mapping (MPM) measures — longitudinal (R_1_) and effective transverse (R_2_*) relaxation rates, normalized proton density proxy (N_A_), and magnetization transfer saturation (MT_sat_) — and tissue-volume change in five whole human brains scanned unfixed *ex vivo in situ*, during formaldehyde immersion fixation, and, when available, after transfer to phosphate buffered saline solution. We included an independent *in vivo* cohort of 25 younger healthy participants to provide a qualitative *in vivo* reference. The largest changes were found for R_1_ during fixation relative to *in situ* values (up to 351% in dGM), followed by R_2_* (up to 60% in WM). N_A_ was largely unchanged during fixation. Although MT_sat_ decreased by up to 25% in gray matter from *in vivo* to *in situ,* it largely preserved tissue contrast during fixation and PBS immersion, making it an ideal candidate for longitudinal registration across tissue conditions. Phenomenological models characterized fixation-induced parameter changes and relative tissue-volume change. R_1_ and fixation-associated volume loss evolved under broadly similar timescales, consistent with a contribution from fixation-related water loss or redistribution. This densely sampled dataset and analyses provide a resource for linking *in vivo* MRI to *postmortem* fixed-tissue MRI and *ex vivo* histology under a standardized immersion-fixation protocol.

## 1. Introduction

Understanding the microstructural underpinnings of magnetic resonance imaging (MRI) contrast in the brain relies on formalin-fixed *postmortem* tissue studies. By stopping autolytic processes, formalin fixation ensures tissue long-term stability by preserving its microstructural content (Beach et al., 1987). Since the early days of MRI, combination of MRI and histology in the same brain tissue has provided key insights into the anatomical and compositional underpinnings of MRI contrast (Drayer et al., 1986). *Postmortem* MRI has gained further importance recently due to the advent of microstructural MRI or MRI histology methods (Weiskopf et al., 2021), where the link between quantitative MRI (qMRI) parameters and tissue microstructure is established using biophysical modeling and same-tissue validation (Eriksson et al., 2007; Kirilina et al., 2020; Stüber et al., 2014).

The multi-parametric mapping (MPM) protocol is a qMRI technique that enables the simultaneous estimation of longitudinal (R_1_) and effective transverse (R_2_*) relaxation rates, proton density (PD) and magnetization transfer saturation (MT_sat_) (Helms et al., 2008; Weiskopf et al., 2013). These MPM parameters are used as microstructural biomarkers in human brain (Freund et al., 2019; Leutritz et al., 2020), aging (Callaghan et al., 2014), pathology (David et al., 2019; Freund et al., 2013), and behaviorally-relevant imaging (Lehmann et al., 2023; Whitaker et al., 2016). To establish these *in vivo* MPM parameters as MRI histological biomarkers (Weiskopf et al., 2021), they need to be validated against *ex vivo* histology of human brains by using the same formalin-fixed tissue, e.g., (Kirilina et al., 2020; Stüber et al., 2014). Such studies have provided many important insights about the validity of MRI-based microstructure estimates, e.g., (Brammerloh et al., 2024; Mordhorst et al., 2025; Papazoglou et al., 2024). However, systematic differences between *in vivo* and *postmortem* fixed tissue limit the direct translation of these results into *in vivo* MRI. Thus, characterizing how MPM parameters change during *postmortem* tissue alterations (death and fixation) are therefore essential.

We treat the transition from *in vivo* to fixed *postmortem* state as several processes. Immediately after death, during the *postmortem* interval, (i.e., the interval between death and before start of fixation, PMI) tissue autolysis leads to enzymatic degradation and disruption of cellular integrity, including myelin sheath separation (Krassner et al., 2023; Shepherd et al., 2009). During this interval, tissue temperature also changes as the body cools and storage conditions vary, which can affect relaxation (Birkl et al., 2016) and diffusion scan sessions (Berger et al., 2023). Animal studies showed that R_1_ decreased and PD increased with increasing *postmortem* interval (Shepherd et al., 2009). After immersion of the excised brain in a fixative solution like formaldehyde, this solution diffuses into the whole brain. Simulations suggest that fixative penetration may take up to ∼40 days (Tendler et al., 2021), which is also in accordance with previous studies (Yong-Hing et al., 2005). The formaldehyde fixation process is temperature-dependent and involves an initial phase 24-28 hours after penetration, in which cross-linking of proteins is initiated, followed by a slower phase of about 30 days during which cross-linking becomes effectively irreversible (Thavarajah et al., 2012). This process changes qMRI parameters in whole human brains (Shatil et al., 2018; Yong-Hing et al., 2005), with several studies reporting increasing R_1_, R_2_* values during fixation (Birkl et al., 2016; Raman et al., 2017), despite differences in fixation duration, tissue size and protocol (Birkl et al., 2016; Raman et al., 2017; Yong-Hing et al., 2005). Lastly, fixed tissue is often transferred to phosphate buffered saline (PBS) solutions to reduce prolonged exposure to concentrated fixative, to stabilize R_1_ values (Shepherd et al., 2009); and to partially recover R_2_ (Shepherd et al., 2009) and R_2_* values (Dusek et al., 2019). While the fixed tissue is immersed in PBS, the bulk fixative in the tissue is partially washed out (van Duijn et al., 2011).

Previous findings are difficult to generalize to intact, whole human brains for four reasons. First, animal brains (e.g. rodent (Shepherd et al., 2009)) and sections of human brain (e.g. hemispheres (Raman et al., 2017)) differ in size and tissue structure (Beaulieu-Laroche et al., 2021). Second, prior whole-brain studies have typically sampled one or a few *postmortem* or fixation states, like *in situ* to *ex situ* comparisons (Neuhaus et al., 2025; Shatil et al., 2018), rather than densely tracking immersion fixation and subsequent PBS wash-out. Third, studies used different T_1_-mapping sequences (MP2RAGE in Shatil et al., (2018) vs. inversion recovery spin echo in Raman et al., (2017)) and fixative concentration, e.g., 4.5% formaldehyde (Neuhaus et al., 2025), 10% and 20% phosphate buffered formalin solution ((Raman et al., 2017) and (Yong-Hing et al., 2005), respectively), which can affect T_1_ estimations (see (Kjaer and Henriksen, 1988; Mezer et al., 2016)) and other qMRI parameters (Birkl et al., 2018). Last, the fixation method itself differed between studies, i.e. perfusion and immersion fixation. Perfusion fixation is widely used in animal studies and has been applied to whole human brains (Insausti et al., 2023; Maranzano et al., 2020). However, its use is constrained by *postmortem* interval, clotting and vascular patency, cannulation requirements, donor-specific vascular pathology, and brain-bank or forensic workflow constraints (McFadden et al., 2019). Therefore, immersion fixation remains widely used for whole human brains. Because it depends on diffusion, immersion fixation is much slower and may prolong the autolysis processes in deeper brain areas. The temporal and spatial progression of the fixation process in human brains remains challenging (Tendler et al., 2021) and depends on factors such as tissue size and fixative concentration. Furthermore, studies have also shown that the time that qMRI parameters like T_1_ and R_2_ require to stabilize depends not only on the tissue type, e.g. white matter or cortex (Yong-Hing et al., 2005), but also varies between samples and donors (Raman et al., 2017). The latter source of variance, hereafter denoted inter-specimen variability, was not systematically characterized in the reported works (Raman et al., 2017; Shatil et al., 2018; Yong-Hing et al., 2005), but could help to assess the generalizability of observed trends. In addition to changing qMRI parameters, aldehyde fixation can alter tissue dimensions non-uniformly. Quester and Schröder (1997) reported that formalin fixation of the human brainstem left transverse dimensions largely unchanged but reduced longitudinal distances by 1–8%, whereas subsequent dehydration and paraffin embedding caused substantially larger shrinkage (Quester and Schröder, 1997). Recent *in situ*-to-*ex situ* MRI comparisons also reported regionally heterogeneous apparent volume differences after extraction and fixation (Neuhaus et al., 2025). Together, these studies indicate that *postmortem* volume change is region-, direction-, time-, and protocol-dependent but do not relate them to associated quantitative parameter changes.

The main objective of this study is to characterize and model changes in the multiple qMRI parameters acquired by the MPM protocol through the *in vivo*-to-*in situ* transition, formaldehyde immersion fixation and PBS immersion of whole human brains. To achieve this, we acquired longitudinal MPM data in five human whole-brain specimens at *in situ* (i.e. in the skull, unfixed), during formaldehyde immersion fixation and during PBS immersion. Our study comprises three key analyses: First, we characterize the changes of MPM parameters between consecutive *postmortem* and fixation processes and compare them to *in vivo* reference values from twenty-five healthy young subjects. Secondly, we propose new fixation models and compare them with the fixation model from Yong-Hing et al. (2005). We characterize the temporal properties of the winning model per parameter and assess how well it predicts the temporal change of each parameter. Finally, we estimate the fixation-induced tissue shrinkage and investigate potential associations with MPM parameter changes.

## 2. Methods

Throughout this work, we use the following terms for specimen conditions at MRI for clarification. *In vivo* denotes MPM data from living participants. *Ex vivo in situ* denotes *postmortem* scans with the brain retained in the skull (unfixed unless stated otherwise). *Ex vivo ex situ* formaldehyde and *ex vivo ex situ* PBS denote scans of excised brains during immersion in formaldehyde or after transfer to phosphate buffered saline (PBS) with 0.1% sodium azide, respectively. For readability, we subsequently refer to these as *in vivo*, *in situ*, formaldehyde *ex situ*, and PBS *ex situ*. Longitudinal MRI scan sessions are distinguished from analysis time points used in modeling (§2.3–§2.5) and from sequence-level acquisitions (§2.2).

### 2.1. *Ex vivo* whole brains and *in vivo* subjects

The *in vivo* dataset consisted of twenty-five healthy subjects (12 female, mean age ± standard deviation: 25.4 ± 2.4 years old) that were recruited as part of a previous study at the University Medical Centre Hamburg-Eppendorf (Ellerbrock and Mohammadi, 2018). All participants in this study were screened for neurological or psychiatric illness (Figure 1A). This study received approval from the local ethics committee (Ärztekammer Hamburg #PV51141) in accordance with the Declaration of Helsinki (seventh revision, 2013).

**Figure 1:**
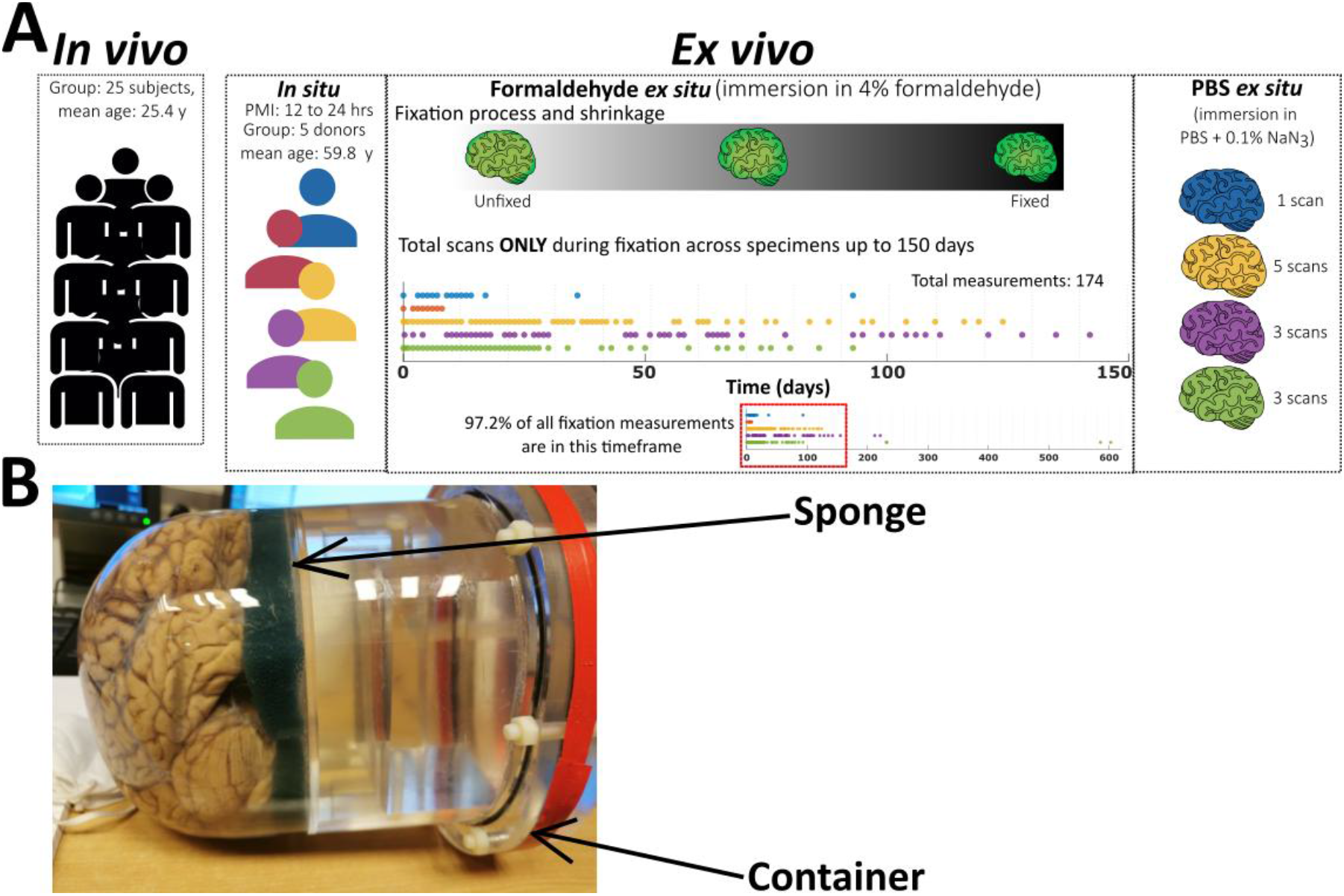
(A) Schematic timeline of MRI scan sessions from in vivo reference data to ex vivo processes. Colors represent ex vivo specimens in Table 1: Brain1 – blue, Brain2 – red, Brain3 – yellow, Brain5 – magenta and Brain6 – green. Dots indicate individual MRI scan sessions during in situ, formaldehyde and PBS ex situ processes. In situ scan session was unavailable for Brain5 and PBS ex situ scan sessions unavailable for Brain2. Formaldehyde ex situ scan sessions were acquired over up to 603 days (bottom panel); temporal modeling of fixation effects used the first 150 days only (middle subplot; 97.2% of all formaldehyde ex situ scan sessions). The PBS ex situ panel shows the number of scan sessions per specimen. For formaldehyde- to-PBS ex situ comparisons (§2.3 and Figure 3) only the last PBS ex situ scan session per specimen was used. PBS: phosphate buffered saline, NaN3: sodium azide. Y: years old (for in vivo and in situ). (B) Photo- graph of a specimen placed in the custom-made MRI-compatible 1H-free acrylic glass holder for all formalde- hyde and PBS ex situ scan sessions.

**Figure 2:**
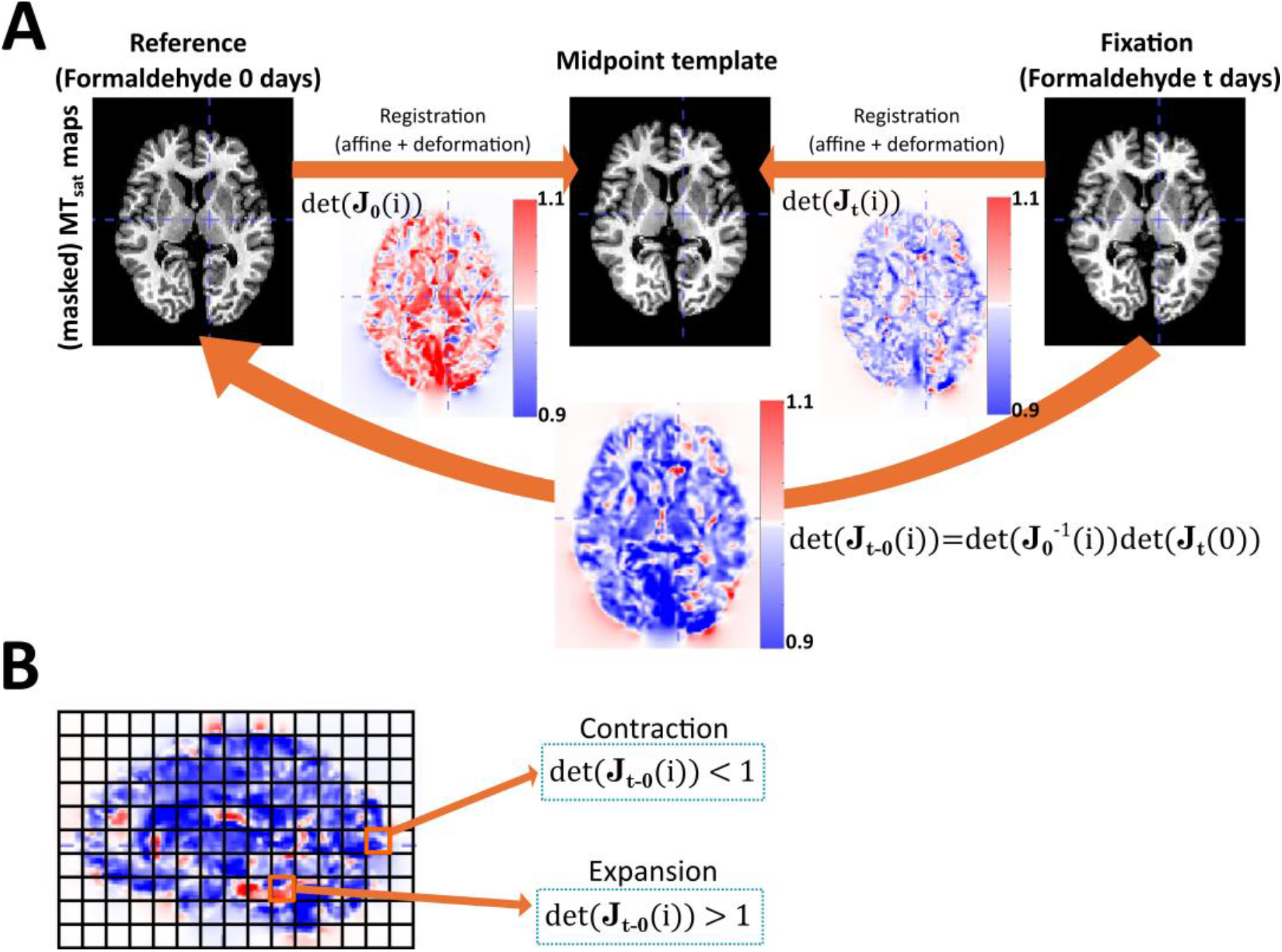
Illustration of the relative volume change estimation and interpretation. (A) We estimated whole- brain volume change from the determinant of the total Jacobian mapping each fixation time point t to the first formaldehyde ex situ scan at t = 0. This transformation combined Jacobian at time t with the inverse Jacobian of the first fixation time point; both were defined relative to the SPM midpoint template. (B) Determinants greater than 1 indicate expansion, whereas values less than 1 indicate contraction. The tissue-class average defines relative volume change (§2.5).

**Table 1:**
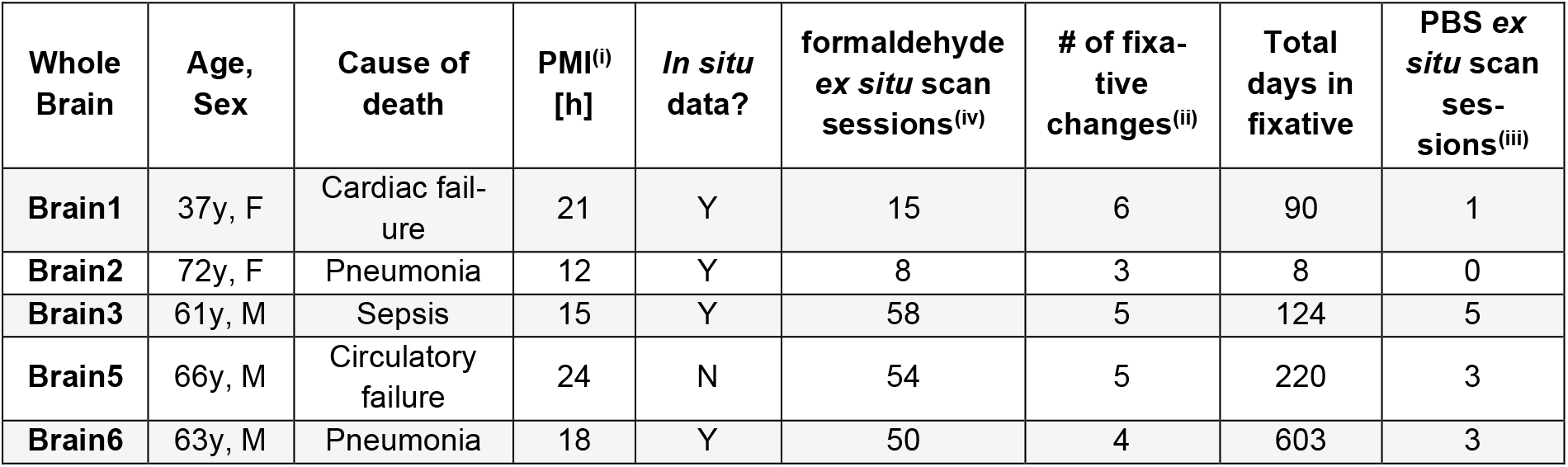
Information on the five ex vivo specimens included in this work. Total days in fixative denote the last day in formaldehyde before transferring to PBS. (i) PMI: postmortem interval, which is the time interval from death to immersion in formaldehyde, (ii) number of formaldehyde solution changes during the initial three weeks of fixation, (iii) number of PBS ex situ MRI scan sessions per specimen (Brain2: none), (iv) number of formaldehyde ex situ scan sessions. This count differs from analysis time points used in later analyses, which may subset scans, i.e., first 150 days of formaldehyde ex situ and last PBS ex situ scan sessions. PBS: phosphate buffered saline.

| Whole Brain | Age, Sex | Cause of death | PMI<sup>(i)</sup> [h] | In situ data? | formaldehyde ex situ scan sessions<sup>(iv)</sup> | # of fixative changes<sup>(ii)</sup> | Total days in fixative | PBS ex situ scan sessions<sup>(iii)</sup> |
| --- | --- | --- | --- | --- | --- | --- | --- | --- |
| Brain1 | 37y, F | Cardiac failure | 21 | Y | 15 | 6 | 90 | 1 |
| Brain2 | 72y, F | Pneumonia | 12 | Y | 8 | 3 | 8 | 0 |
| Brain3 | 61y, M | Sepsis | 15 | Y | 58 | 5 | 124 | 5 |
| Brain5 | 66y, M | Circulatory failure | 24 | N | 54 | 5 | 220 | 3 |
| Brain6 | 63y, M | Pneumonia | 18 | Y | 50 | 4 | 603 | 3 |

For the *ex vivo* dataset, deceased donors were received at the Department of Legal Medicine and stored without refrigeration at approximately 16°C. Donors were included according to pre-specified criteria: no known neurological disease based on available records; an intact brain without macroscopic trauma expected to alter morphology; and a postmortem interval (PMI) not more than 24 h, defined as the time interval from death to brain extraction and immersion in formaldehyde (Table 1). We included five brains that met these criteria (2 females, mean age ± standard deviation: 59.8 ± 13.4 years old; ethics approval WF-74/16); further specimen details, including fixation duration, are summarized in Table 1. During *algor mortis*, all donors except Brain5 underwent whole brain *in situ* MRI (Table 1). Two whole-brain movies of each specimen are provided in Supplementary Materials §S6 to illustrate that no visible white matter hyperintensities (WMHs) were detected. All five donors were subsequently extracted at autopsy using an electrical saw, with care taken to avoid cutting into the specimen. The leptomeninges were not removed and none of the major vessels were ligated. The transfer and excision processes occurred in most cases within 24 h after death (Table 1). After extraction, specimens were immersed in 4% formaldehyde (acid-free 37% formaldehyde for histology, Roth item no. P733.2, diluted with PBS^1^; pH-value 7.4). The formaldehyde solution was renewed during the initial three weeks to support uniform immersion fixation. Brain2 was fixed for only eight days because it was subsequently used for later histological studies unrelated to this work.

After the formaldehyde immersion period, all specimens except Brain2 were transferred to PBS containing 0.1% sodium azide and remained in this solution for at least 7 days before performing the first PBS *ex situ* MRI scan session (Table 1). Specimens contributed one to five PBS *ex situ* scan sessions in total (Supplementary Materials §S2 and Table 1).

Brains were placed in a custom-made MRI-compatible, ^1^H-free acrylic glass holder to improve repositioning across longitudinal scans (Figure 1B). Visible bubbles were reduced by gently massaging them out of the specimens. Unlike Neuhaus et al. (2025), who used brain-specific support plates, and Boonstra et al. (2021), who used a vacuum-sealed container, our holder was designed primarily for longitudinal repositioning. The holder was larger than the specimens and was not designed to compress the tissue. Contact with the support surface may have altered local brain shape, but substantial effects on global fixation-associated volume changes are unlikely under this setup (see §2.5).

### 2.2. MR acquisition and preprocessing

Whole-brain MR images were acquired for healthy controls (*in vivo* dataset) and for *ex vivo* specimens. For the latter, the MR images were acquired throughout *in situ*, formaldehyde and PBS *ex situ* scan sessions, when available (see Table 1). All MR images were acquired with the MPM protocol (Weiskopf et al., 2013). *In vivo* data were acquired on a 3 T TRIO scanner, whereas *in situ* and *ex situ* data were acquired on the same 3 T PRISMA^fit^ (Siemens Healthcare, Erlangen, Germany). Both systems used a Siemens 32-channel radiofrequency receive head-coil. All *in situ* and *ex situ* sessions were scanned in an air-conditioned room held near 22.5°C by site staff; this set-point was not logged per session, and bore temperature is used by the system for SAR limits rather than specimen temperature regulation (MR facility, personal communication). This protocol consisted of performing a transmit, B_1_+ field measurement for flip angle correction (Lutti et al., 2012) followed by three 3D multi-echo spoiled fast-low-angle-shot (FLASH) sequences. These three FLASH sequences generated MR signals weighted by T_1_ (T_1_w), PD (PDw), and MT (MTw). All the MPM parameter maps were estimated using the hMRI toolbox version 0.6.1-dev (Github commit hash: 6cbdb42) (Tabelow et al., 2019) integrated into SPM 12 (Friston, 2007) within Matlab R2021b (MATLAB. (2021). 9.11.0.1837725 (R2021b) Update 2. Natick, Massachusetts: The MathWorks Inc.). A detailed description of MR acquisition and preprocessing for *ex vivo* specimen and *in vivo* subjects is given as follows:

#### In situ and ex situ MPMs

Both *in situ* and *ex situ* MPM data were acquired with an isotropic resolution of (0.8 mm)³. The nominal excitation flip angle (FA) was 6° for MTw and PDw, and 21° for T_1_w; with eight echo times (TE) from 2.34 to 18.44 ms for PDw and T_1_w and six echoes to 13.84 ms for MTw in steps of 2.30 ms. The readout bandwidth was 488 Hz/pixel and the repetition time (TR) was 25.0 ms.

#### Processing and map creation of in situ and ex situ MPMs

The acquired MPM data were processed using the hMRI toolbox. Prior to map creation, the maps were denoised using the local complex Principal Component Analysis (LCPCA) approach (Bazin et al., 2019; Edwards et al., 2024). The map creation module yielded the MPM parameter maps MT_sat_, R_1_, R_2_* and uncalibrated proton density, A, corrected for B_1_+ biases (Weiskopf et al., 2013), as well as the R_2_*-error (ε) map based on the ESTATICS model (Weiskopf et al., 2014).

Because the ex vivo specimens were scanned at 3 T in solution-filled containers, small residual air bubbles were occasionally present near the tissue surface, within the ventricles, or within other fluid spaces. Although susceptibility effects are expected to be less pronounced at 3 T than at ultra-high field, FLASH acquisitions—particularly R2*— remain sensitive to local tissue–air interfaces. ESTATICS ε maps were therefore used to identify affected voxels. Voxels with elevated ε values consistent with bubble-related artifacts were excluded from subsequent analyses. The complete thresholding procedure, based on multilevel Otsu thresholding, is described in Supplementary Materials §S1, and the proportion of excluded brain-mask voxels is summarized per specimen in Supplementary Figure S2. Furthermore, the effect of bubble masking is illustrated in Figures S3 and S4.

Registration involved two complementary approaches. For the MPM-parameter analyses (see §2.3 and §2.4), we created a longitudinal template from all formaldehyde and PBS *ex situ* scans using the optimized ANTs script antsMultivariateTemplateConstruction (https://github.com/CoBrALab/optimized_antsMultivariateTemplateConstruction/?tab=readme-ov-file, Github commit hash b340447, (Germann et al., 2025)). Then, we registered the *in situ* scan to this template. For the volume-change analysis (see §2.5), we registered only the formaldehyde and PBS *ex situ* scans using the longitudinal registration in SPM (Ashburner and Ridgway, 2013). We selected the SPM method because its spatial constraints are already optimized for morphometric analysis. Simultaneously, the ε maps were also transformed using the previously estimated transforms. A detailed description of each registration method is given in Supplementary Materials §S1.

For each specimen, all scan sessions were registered either to the *in situ* or the first formaldehyde *ex situ* scan, which served as the reference scan session. We segmented the reference MT_sat_ and R_1_ with SPM into white matter (WM), cortical (cGM) and deep gray matter (dGM), and cerebro-spinal fluid (CSF) regions. This CSF map was also estimated for Brain5, even though the reference scan was already the first formaldehyde *ex situ* scan session. To avoid confusion, we hereafter call it the “fluid map” for all specimens. This estimated fluid map was used for normalization of the A parameter, hereafter the normalized proton density N_A_, by dividing the A values in the whole brain by the median of A in the fluid map per specimen and time point, i.e. N_A_(t) = A_tissue−class_(t)/ 〈A〉_fluid−mask_(t). This normalization accounts for the variation in the A parameter due to sample positioning, especially during early formaldehyde solution changes (Table 1); coils and gradients stability (Mezer et al., 2016), and differing experimental conditions across *postmortem* processes. Therefore, we only apply this normalization for both formaldehyde and PBS *ex situ* scan sessions. Detailed information about preprocessing, including registration and segmentation, can be found in the Supplementary Materials §S1.

The tissue registration and segmentation were performed using ANTs version 2.6.3 and SPM 12 as described above.

#### Processing and map creation of in vivo MPMs

The *in vivo* MPM data were acquired with equal sequence parameters as for *ex vivo*, and previously used in another study (Ellerbrock and Mohammadi, 2018).

The estimation of the *in vivo* MPM parameter maps followed the procedures described in (Emmenegger et al., 2021) and (Mohammadi et al., 2022). This procedure was identical for the *ex vivo* MR images, except for the calibration of the proton density, PD (here N_A_). Afterwards, the MPM parameter maps were spatially normalized to MNI space and averaged across subjects following the methodology detailed in (Emmenegger et al., 2021). The average and MNI-registered MPM parameter maps were also segmented into cGM, dGM and WM.

### 2.3. Analysis 1: Comparison of the MPM parameters during *in situ*, formaldehyde *ex situ*, and PBS *ex situ* conditions

To assess the transition in the MPM parameters, i.e. R_1_, R_2_*, MT_sat_, N_A_, across specimen conditions, we estimated the relative change of the mean medians per tissue class (cGM, dGM and WM). Relative mean median changes were estimated between consecutive tissue conditions (*in vivo, in situ*, formaldehyde and PBS *ex situ*; Figure 1A). Within the formaldehyde *ex situ* phase, we additionally investigated days 30 and 90. Day 30 corresponds to the days after which most of the brain was penetrated by the fixative solution (30-46 days, (Tendler et al., 2021)), whereas day 90 corresponds the latest formaldehyde *ex situ* scan session across all brain specimens.

To estimate the relative change of the mean medians between consecutive tissue conditions (Figure 1A), we used a hierarchical statistical approach. First, for each brain specimen, tissue class (dGM, cGM, WM), and scan session, we computed the median MPM parameter value across all voxels within that tissue class (median(MPM)). Second, we aggregated these tissue-class-level medians across specimens and took the relative difference as follows: 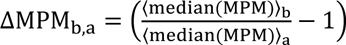, where ⟨median(MPM)⟩_a_ and ⟨median(MPM)⟩_b_ denote the average across specimens at stages a and b (e.g. “a” being *in vivo* and “b” being *in situ*). For formaldehyde *ex situ* scan sessions, where individual brains contributed with multiple sessions at different fixation durations (last formaldehyde *ex situ* session from 93 to 603 days), we computed the average of relative differences of medians across specimens, defined as: 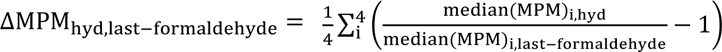 where the subscript i indicates the specimen. This approach ensures equal weighting of each specimen, accounting for inter-specimen variability. In this latter calculation, we only used four specimens, since Brain2 had no PBS *ex situ* scan sessions (Table 1).

Because the *in vivo* cohort differs from the postmortem specimens in age, scanner, and temperature, the *in vivo*–*in situ* comparison is reported qualitatively, without statistical testing. For the within-specimen transitions (*in situ* to the first formaldehyde *ex situ* scan, and between fixation time points), where the same specimens were compared across conditions, we report the relative median change with its across-specimen spread and, as a descriptive index only, the exact p-value from a paired t-test. Given the small sample (n = 4–5), we make no claims of statistical significance and interpret all comparisons descriptively.

### 2.4. Analysis 2: Modeling of formaldehyde *ex situ* fixation effects on the MPM parameters

To assess whether the temporal variation of the MPM parameters is specimen-independent and can be modeled, we described the temporal evolution of each *ex vivo* (ev) MPM parameter ⟨R_ev_⟩(t), with R_ev_ = {R_1_, R^∗^, MT_sat_, N_A_}. For this analysis, we included all the formaldehyde *ex situ* scan sessions up to 150 days in fixative and excluding all later scan sessions (Table 1 and Figure 1A). For each MPM parameter and tissue class, we compared four candidate models using the corrected Akaike Information Criterion (AICc) (Burnham et al., 2011; Ghosh et al., 2007).

The first model is the reciprocal logistic model from Yong-Hing (Yong-Hing et al., 2005):

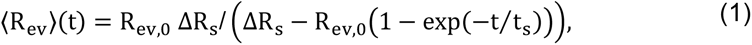

where R_ev,0_ corresponds to the MPM parameter in fixative at day zero, i.e. R_ev,0_ = ⟨R_ev_⟩(t = 0), ΔR_s_ is the change in the MPM parameter due to the fixative and t_s_ is the saturation time. In this context, “s” in ΔR_s_ and t_s_ denotes “saturation”. R_ev,0_ and ΔR_s_ parameters are described in 1/seconds when R_ev_ = {R_1_, R^∗^} or in the respective unit for R_ev_ = {MT_sat_, N_A_}; t_s_ is given in days. The original model was formulated for T_1_ and T_2_; i.e., 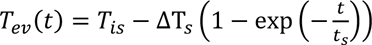, here we use its reciprocal form for R_1_ and R_2_*.

For the next three models, we used saturation-type functions of the form ⟨R_ev_⟩(t) = R_ev,0_ + s(t), where s(t) captures fixation-related changes across all specimens. A linear iron–myelin reading of s(t), distinct from the fitting itself, is given in Supplementary Materials §S7. The first of these saturation models is the mono-exponential model, which postulates that the changes in relaxation rates between R_ev,0_ and ⟨R_ev_⟩ can be expressed through an exponential saturation process defined by

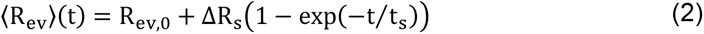

The second saturation model is the bi-exponential model, which is based on the same principle underlying the mono-exponential model but proposes that the fixation process in the tissue occurs over two timescales, as proposed in (Thavarajah et al., 2012)

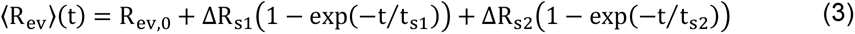

wherein ΔR_S1_ and t_S1_ describe the fast fixation process shortly after fixative penetration and ΔR_s2_ and t_s2_ the longer time fixation processes.

The last saturation model is the null model, which is used as a baseline relative to the previous models and represents the scenario where the MPM parameter has no temporal dependency. This is simply defined as

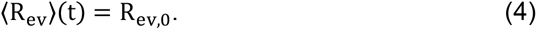

The fitted parameters are R_ev,0_, ΔR_s_, ΔR_s1_, ΔR_s2_, t_s_, t_s1_ and t_s2_. We defined these three models as saturation models based on two hypotheses, complementary to the motivation mentioned previously: first, the fixative solution diffuses into the tissue until it reaches a maximum concentration of the chemical and, second, the formation of cross links which changes the tissue properties and therefore MPM parameters also follows a saturation process. Here, “saturation time” should not be interpreted as the time that the MPM parameter requires to reach a stable plateau at that specific fixative concentration.

To compare the fitted parameters of the mono- and bi-exponential saturation models between tissue classes and MPM parameters, we harmonized these parameters by following the next definitions in Table 2.

**Table 2:**
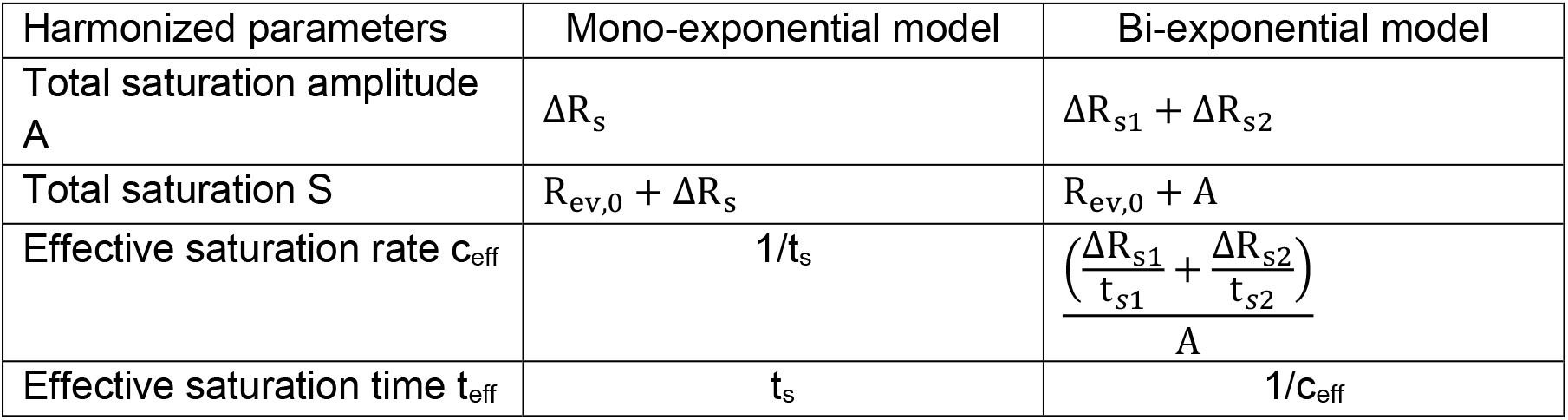
Definition of the harmonized parameters as a function of the fitted parameters from the mono- and bi-exponential models. Standard deviation of each harmonized fitted parameter is estimated via error propagation.

Standard deviation of these harmonized parameters was also estimated via firstorder error propagation, by assuming that the fitted parameters are uncorrelated.

To select the preferred model for each MPM parameter, we first calculated the AICc:

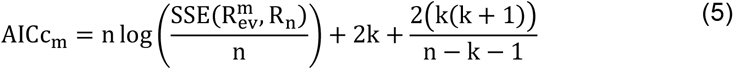

where “n” is the number of time samples, “k*”* is the number of parameters in model “m” and 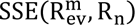 is the sum of squares residual of the fitted model. The preferred model is the one with the lowest AICc, i.e., AICc_min_.

Then we summarized the relative support across all competing models per MPM parameter and tissue class by the Evidence Ratio (ER), defined as ER(AICc_min_, AICc_m_) = exp(0.5(AICc_m_ − AICc_min_)), where the subscript “m” indicates the models (Equations 1 to 4). The preferred model has always ER = 1. Larger ER values indicate progressively weaker support. Models with ER < 2.71 were considered equally plausible, whereas models with ER > 148.41 were discarded (see (Burnham et al., 2011; Burnham and Anderson, 2004, 2002) and Supplementary Materials §S3 for more details).

To assess the agreement between the predicted and observed temporal trends in each MPM parameter, Kendall’s τ coefficient was used (Kendall, 1938). Kendall’s τ coefficient evaluates the rank correlation between the preferred model and observed values, offering a non-parametric measure of predictive accuracy. A high τ value indicates a strong agreement between the predicted and observed temporal trends. In addition, RMSE was calculated as a complementary error metric to assess absolute agreement on the predicted temporal trends and the estimated MPM parameters. To further assess the influence of inter-specimen variability, τ and RMSE were determined in a leave-one-out cross-validation (see (Molinaro et al., 2005)). We investigated the median of τ and RMSE across specimens to quantify the error and used the interquartile range (IQR) as a measure of their variation, i.e. *median (IQR: Q1 Q3)*, with Q1 being the 25% and Q3 being the 75% quartiles. When necessary, we additionally reported the magnitude of this range, i.e. the difference between these quartiles: Q3-Q1.

### 2.5. Analysis 3: relative volume change during *ex vivo-ex situ* fixation

To model relative tissue-volume change during formaldehyde *ex situ* condition, we characterized its temporal change, rV(t), with respect to the first formaldehyde *ex situ* scan for all specimens per tissue class. For that, we calculated the determinant of the Jacobian of the nonlinear transformation from the SPM registration (§2.2) (Ashburner and Ridgway, 2013) that mapped the formaldehyde *ex situ* scan at each time point t back to the reference formaldehyde *ex situ* scan t0 per voxel i: det(*J_t_*_−*t*0_ (*i*)) (details about this composition can be found in Supplementary Materials §S1, in *Jacobian composition*). Five movies, i.e., one per brain specimen, were created to show the resulting determinant of the Jacobian at each time point. For each time point, rV(t) was calculated by averaging this determinant per tissue class, i.e., 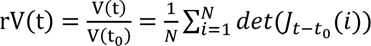, where N is the total number of voxels in such tissue class. Each tissue class was defined based on the reference formaldehyde *ex situ* scan.

Relative volume change was referenced to the first formaldehyde *ex situ* scan rather than to the *in situ* scan, because volume is derived from the deformation field. During the *in situ*-to-*ex situ* transition, deformation is dominated by extraction and handling-related morphological change; this precludes interpreting an *in situ*-to-formaldehyde volume difference as fixation-induced shrinkage. We therefore did not estimate an *in situ*-to-day- 0 volume change. PBS *ex situ* volume change was not modeled as a longitudinal trajectory. Instead, we made an exploratory comparison between each specimen’s last formaldehyde *ex situ* scan and its latest PBS *ex situ* scan session, as reported in Supplementary Materials §S5.2 and Figure S19.

To describe the measured volume change, we proposed a heuristic model which assumes an additive exponential change in volume (heuristic model) like the model in Equation 2:

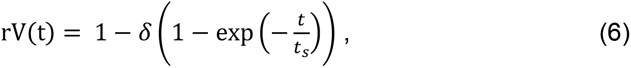

where δ is the dimensionless decrement and t_s_ the “saturation time” at which 63.2% of the relative change took place in days (cf. Equation 2). In this model, δ and t_s_ were fitted to rV(t).

Like in the previous analysis (§2.4), we performed model selection between Equation 6 and the null model (cf. Equation 4) using the AICc and assessed the agreement between the predicted and observed temporal trends in each MPM parameter with the Kendall’s τ coefficient and RMSE through the leave-one-out cross-validation. We also investigated the median of τ and RMSE across specimens to quantify the error and used the interquartile range (IQR) as a measure of their variation, i.e. reported as *median (IQR: Q1 Q3)*, with Q1 being the 25% and Q3 being the 75% quartiles. When necessary, we additionally reported the magnitude of this range, i.e. the difference between these quartiles: Q3-Q1.

In a complementary analysis, we correlated both the temporal changes of the MPM parameters and tissue shrinkage (via rV(t)) across specimens per tissue class. We determined both the correlation coefficient (r) and its p-value.

## 3. Results

### 3.1. Comparison of the MPM parameters during *in situ*, formaldehyde *ex situ*, and PBS *ex situ* conditions

Figure 3 illustrates the means and standard deviation across specimen MPM parameter medians (rows) at different *postmortem* and fixation-related conditions (x-axis) per tissue class (columns). The pairwise percentages compare those mean medians between consecutive *postmortem* conditions (Methods §2.3). Descriptive p-values below 0.05 are marked in Figure 3 (*, **, ***); exact values are given in the text.

**Figure 3:**
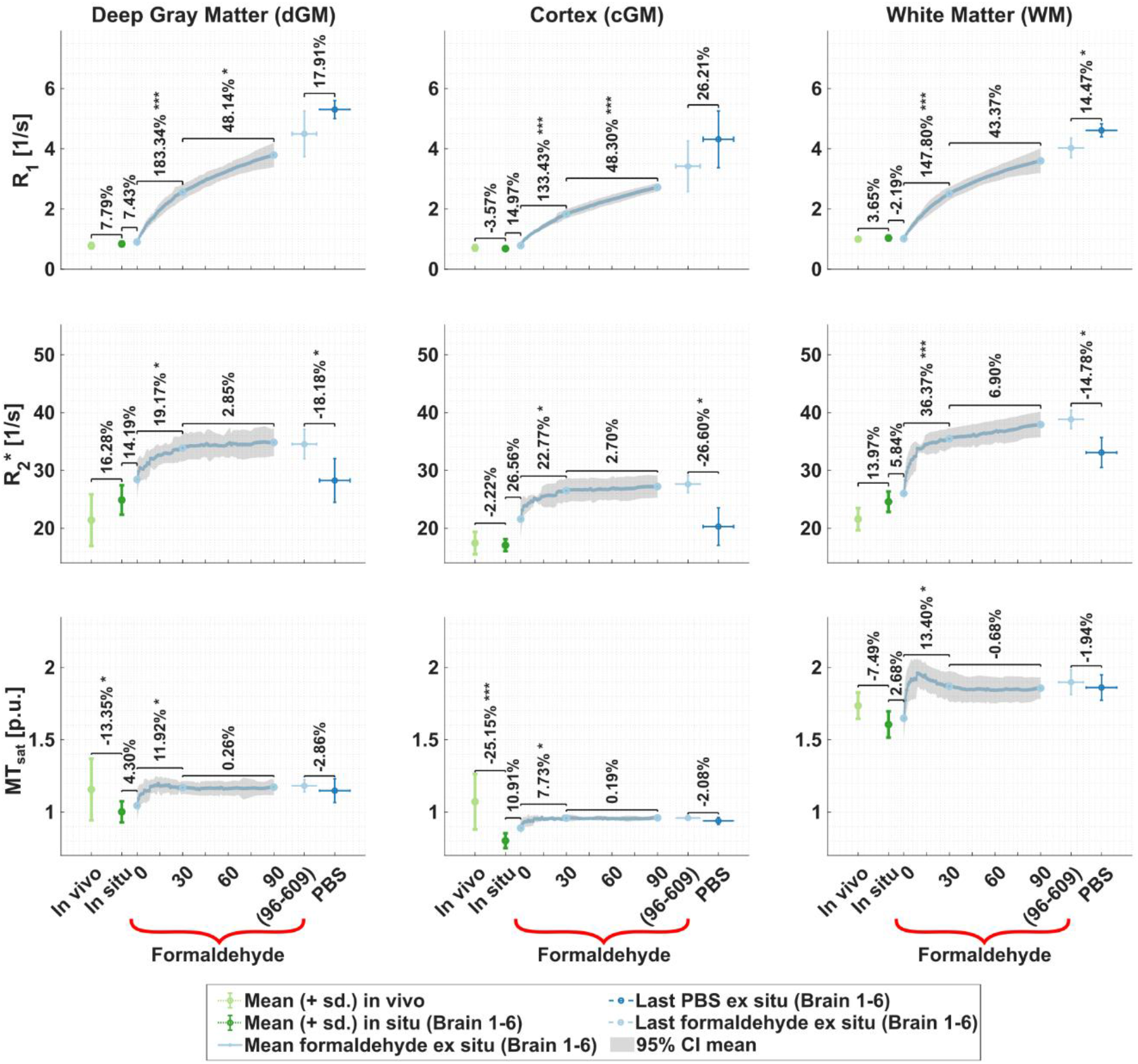
Evolution of the MPM parameters (R_1_, R_2_* and MT_sat_) across postmortem and fixation processes for all specimens. Summary markers show mean and standard deviation across specimen medians at different times and postmortem and fixation processes per tissue class (columns from left to right: deep gray matter, cortical gray matter, and white matter). Pairwise percentages compare those mean medians between consecutive postmortem processes in percentage: in situ and in vivo, formaldehyde ex situ scan session at day 0 and in situ, formaldehyde ex situ scan sessions at day 30 and day 0, at day 90 and day 30; and between last formaldehyde and last PBS ex situ scan sessions. Star symbols denote the magnitude of the descriptive p-value (**not statistical significance**): * reflects p-value < 0.05; ** reflects p-value < 0.01 and *** reflects p- value < 0.001. Exact p-values are reported in Results §3.1. The last PBS ex situ scan was acquired when the specimen was, at least, day 7 in PBS + 0.1% sodium azide. The shaded curve during the fixation time represents the 95% confidence interval (CI) across specimens and the last formaldehyde scan session across specimens and time (between 93 to 603 days) is displayed by the x-y error bar and used to compare with the last PBS ex situ scan session.

Each MPM parameter exhibited a characteristic pattern throughout the different *postmortem* processes independently of the tissue class. For example, R_1_ and R_2_* varied strongly during fixation up to 30 days, while MT_sat_ had the most pronounced change between the *in vivo* reference and *in situ*. Additionally, MT_sat_ displayed a peak within the first 10 days of fixation which was not observed in any other MPM parameter. R_1_ and R_2_* showed noticeable changes during the transition from formaldehyde to PBS immersion, while MT_sat_ exhibited almost no change.

During the transition from *in vivo* to *in situ*, WM-R_2_* and dGM-R2* increased by up to 16.28%, whereas dGM-MT_sat_ and cGM-MT_sat_ decreased by up to 25.15%. The remaining MPM parameters exhibited negligible changes of less than 8%.

In the transition from *in situ* to formaldehyde *ex situ* on day 0 almost all the MPM parameters increased, ranging from 2.68% for WM-MTsat to 26.56% for cGM-R2*. The exception was WM-R1, which decreased by 2.19%. An exploratory plot of these first formaldehyde values against PMI is given in Supplementary Materials §S4.6.

Moving from *formaldehyde ex situ* at day 0 to day 30, R_1_, R_2_* and MT_sat_ parameters increased. The increase was substantial for R_1_ (from 133.43% to 183.34%, with p-values from 1.8e^-6^ to 5.527e^-4^), moderate for R_2_* (from 19.17% to 36.37%, with p-values from 1.965e^-5^ to 2.52e^-2^), and smallest for MT_sat_ (from 7.73% to 13.40%, with p-values from 2.63e^-2^ to 4.55e^-2^). While R_2_* and MT_sat_ parameters showed a fast initial increase followed by a slower increase (R_2_*) or plateau (MT_sat_), R_1_ showed a continuous steep increase.

From day 30 to day 90, R_1_ further increased by an additional 43.37% in cGM to 48.30% in dGM (respective p-values of 8.67e^-5^ and 2.11e^-2^). R_2_* increased by at most 6.90%, and MT_sat_ and N_A_ changed by less than 0.8%.

Lastly, in the comparison between last formaldehyde *ex situ* scan session (light blue error bar) and its last PBS *ex situ* scan session (dark blue error bar), R_1_ increased by up to 26.21%, including 14.47% in WM (p-value = 2.95e^-2^). R_2_* decreased by up to 26.60% (p-values = 1.31e^-2^ to 3.76e^-2^), whereas MT_sat_ varied by no more than 2.86%.

Per specimen trends were more variable and occasionally opposite to the across-specimen mean; a detailed analysis is reported in Supplementary Materials §S2.

### 3.2. Modeling of formaldehyde *ex situ* fixation effects on the MPM parameters

We first report the preferred temporal model per MPM parameter based on the lowest AICc value (Equation 5) shown in Figure S7 (Supplementary Materials §S3.1). We then show specimen-wise trajectories and the fitted preferred models during the formaldehyde *ex situ* phase (Figure 4). Predictive performance of the preferred model was assessed with “leave-one-out” cross-validation using Kendall’s τ coefficient shown in Figure S9 (Supplementary Materials §S3.1).

**Figure 4:**
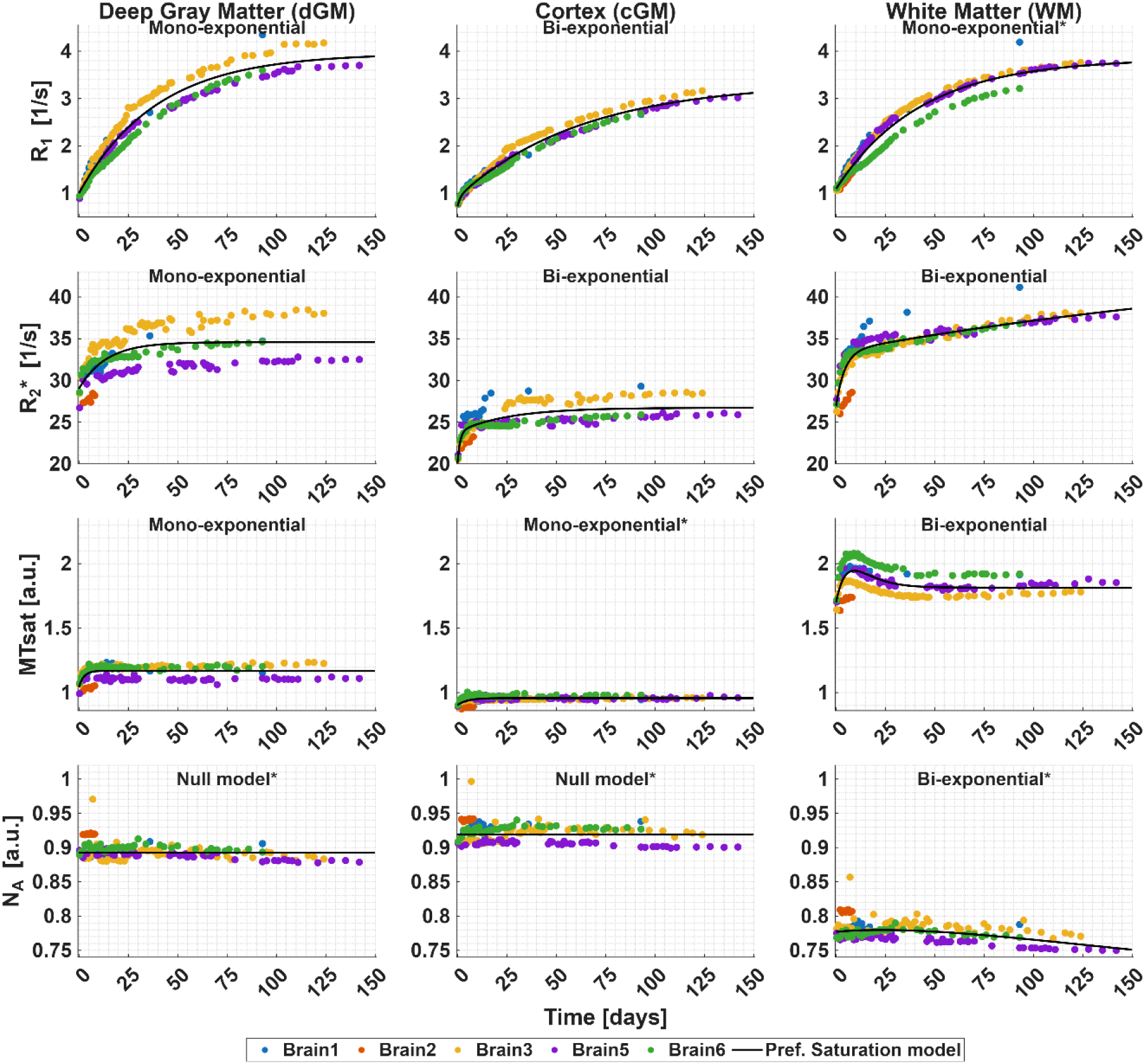
Temporal evolution of the MPM parameters during fixation. For each tissue class (columns, left to right: deep gray matter, dGM; cortical gray matter, cGM; and white matter, WM), the median value of the MPM parameters is plotted (rows, top to bottom: R_1_, R_2_*, MT_sat_ and N_A_) for each brain specimen (colored markers) and per time point. The curve of the preferred saturation model (⟨R_ev_⟩(t)) is plotted for each tissue class and MPM parameter (black solid line) and its name shown on top of each subplot. For better visualization of temporal trends, a version with independent y-axis scaling for each subplot is shown in Figure S8 in Supplementary Materials §S3.1.

For R_1_, R_2_*, and MT_sat_, the monoand bi-exponential saturation models (Equations 2 and 3) were generally preferred over the reciprocal logistic and null models (Figure 4; Figure S7 and Table S2 in Supplementary Materials §S3.1). The bi-exponential decay best described the increase in cGM- and WM-R_2_*, cGM-R_1_, and WM-MT_sat_ (Figure 4). The mono-exponential model best described WM- and dGM-R_1_, dGM-R2*, and cGM- and dGM-MT_sat_. The changes in N_A_ were best described by the null model, except in WM, where the bi-exponential saturation model was preferred.

Quantitatively, the saturation times varied moderately across MPM parameters but were similar across tissue classes. The fast and slow change seen in R_2_* and the MT_sat_ overshoot were observed between 5 to 15 days of fixation in both WM and dGM. Notably, the peak of the overshoot in WM-MT_sat_ occurred at the same time when WM-R_2_* changed its trend from fast to slower increase, i.e. ∼10 days (Figure 4 and in more detail in Figure S8 in Supplementary Materials §S3.1). The effective saturation time, i.e. t_eff_ in Table 2 from Methods §2.4, is reported below. For R_1_, t_eff_ was 39.2321 ± 2.4845 days for dGM, 15.1349 ± 0.6564 days for cGM, and 42.9235 ± 2.4101 days for WM. For R_2_*, these times were 13.6629 ± 3.0990 days for dGM, 2.2080 ± 0.5467 days for cGM, and 12.7190 ± 0.2040 days for WM. For MTsat, these times were 2.4789 ± 0.9273 days for dGM, 4.9290 ± 1.2452 days for cGM, and 1.4731 ± 595.94 days for WM. Note that WM-MTsat had an effective saturation time’s standard deviation larger than the estimate, indicating an unstable fit. A detailed list of the fitted parameters per MPM parameter and tissue contrast can be found in Table S3 from Supplementary Materials §S3.1, and for all the harmonized parameters in Table S5 from the same section.

In the leave-one-out cross-validation analysis, the highest median τ (interquartile range, IQR: (Q1 Q3)) for R_1_ was 1.000 (0.9881 1.000) across all tissue classes. The τ and IQR for R_2_* were smaller between 0.7365 (0.6300 0.8050) for cGM and 0.9471 (0.9024 0.9670) for WM, and it was even lower for MT_sat_ from 0.4161 (0.1159 0.5000) for dGM to 0.6044 (0.3836 0.8271) for WM. Kendall’s τ coefficient was not calculated for N_A_, because, by definition, the null-model assumes no temporal trend. This also applied for the bi-exponential model for WM-NA; since this model is highly overfitted (Supplementary Materials §S3.1, Table S3).

### 3.3. Relative volume change during ex vivo-*ex situ* fixation

Figure 5 shows the temporal change of relative volume, rV(t), in dGM (left), cGM (middle) and WM (right), normalized to the first formaldehyde *ex situ* scan session volume. We quantified how well the preferred model characterized the temporal data using “leave- one-out” cross-validation (Supplementary Materials §S3.2, Figure S11). Figure 6 shows the correlation between rV(t) and the temporal changes of MPM parameters, whereas an exploratory comparison between 1/rV(t) and MPM’s is shown in Supplementary Materials §S5.1 (Figure S18).

**Figure 5:**
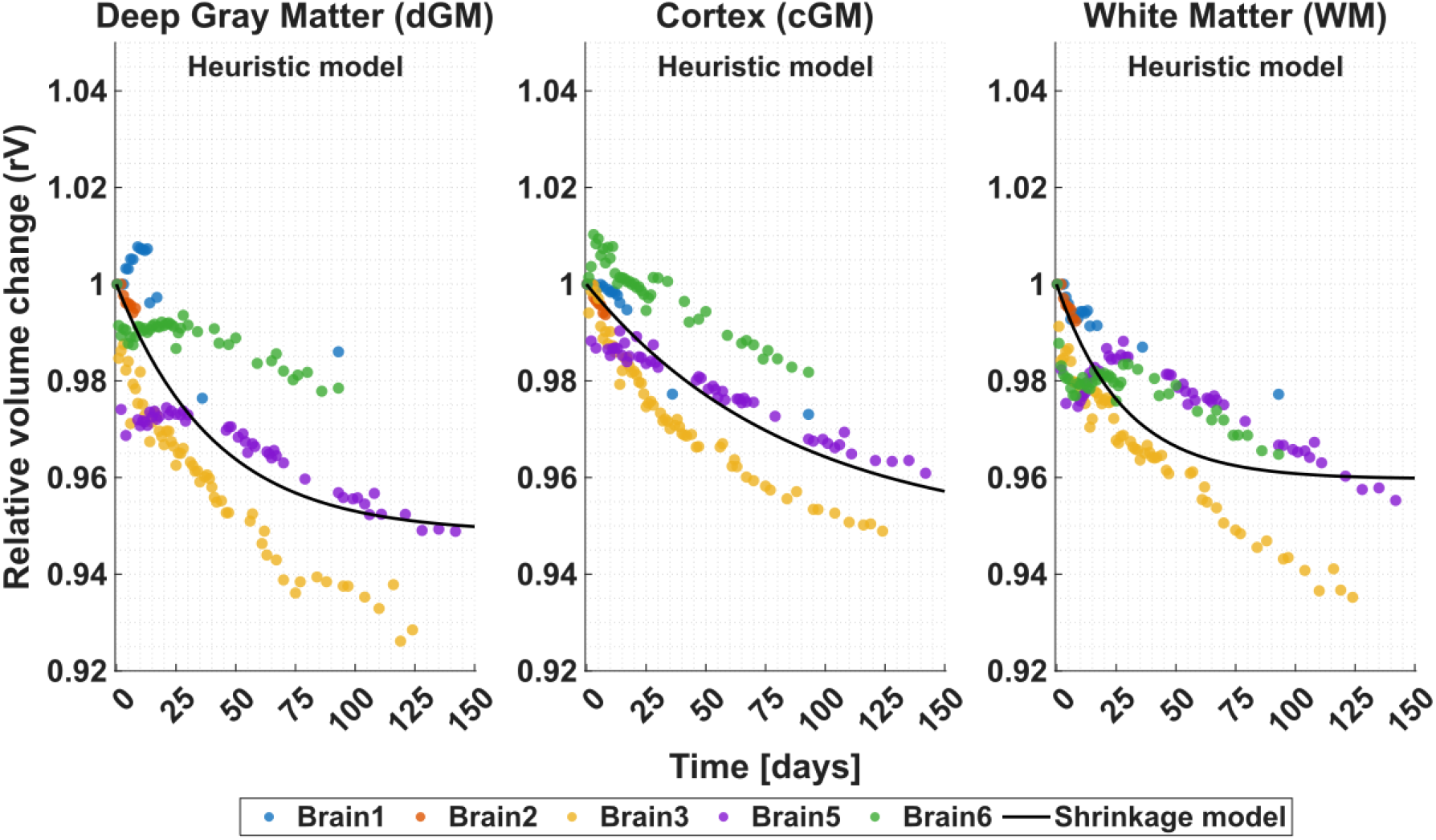
Temporal relative volume change of brain specimens (color) and per tissue class (left to right): deep and cortical gray matter, and white matter. This relative volume change was estimated by the average determinant of the Jacobian (the volume changes relative to the first fixation measurement) per time point. Additionally, the black line shows the fitted relative volume change model (rV(t)).

**Figure 6:**
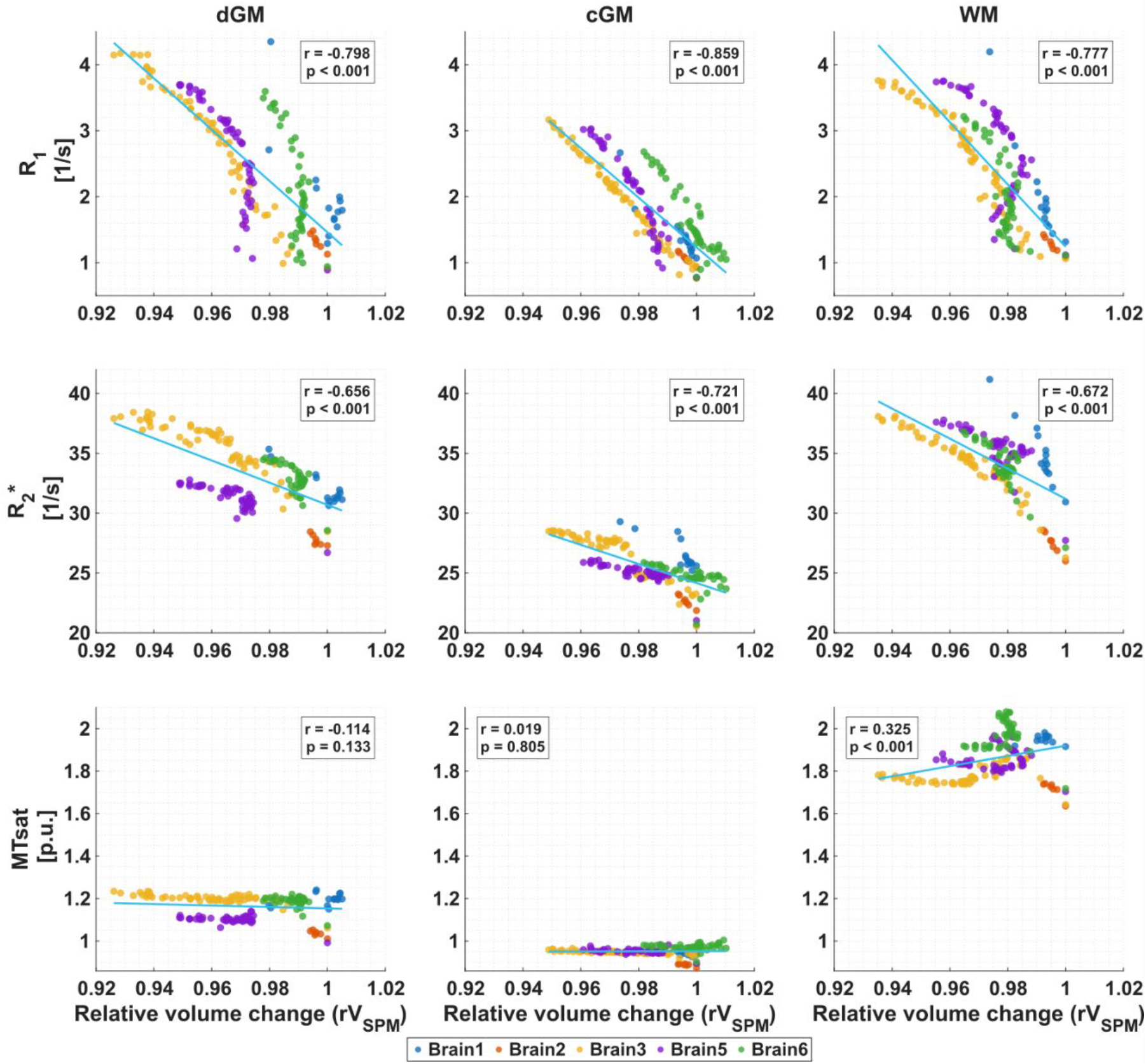
The correlation between the relative volume change model (rV(t)) and the MPM parameters R_1_, R_2_*, and MT_sat_ for each tissue class and across all specimens (color-coded dots). The correlation line overlaid in each subplot (blue line) and the magnitude of the p-value is shown in inset (if lower than 0.001, just showed as an interval). Inset: the correlation coefficient (r) and the magnitude of the p-value. Top row: rV(t) vs. R_1_; middle row: rV(t) vs. R_2_*; bottom row: rV(t) vs. MT_sat_.

For all tissue classes, the heuristic model (cf. Equation 6) described the smooth decay of rV(t) and revealed that the estimated decrement coefficient δ reflected shrinkage at infinite time between 4.031% and 5.178% (0.04031 ± 0.00193 for WM, 0.05149 ± 0.00398 for dGM and 0.05178 ± 0.007996 for cGM, Supplementary Materials §S3.2, Table S7). The saturation times *t_s_* per tissue class were: 41.118 ± 6.7688 days for dGM, 85.4 ± 20.647 days for cGM and 27.976 ± 3.3745 days for WM. Brain5 and Brain6 showed atypical rV(t) trajectories. In Brain6, relative volume increased in cGM, whereas WM showed a strong decrease followed by a partial rebound; both patterns persisted until about day 25 and then declined. Brain5 showed the same WM pattern and a weaker version of the Brain6 cGM pattern in dGM.

Across tissue classes, the median τ (interquartile range: (Q1 Q3)) between individual trends and the fitted model was generally highest in cGM with value 0.9121 (0.8444 0.9627), followed by WM with value 0.78022 (0.54013 0.9390), and lowest and more variable in dGM with value 0.75 (0.32323 0.9111). Interestingly, the magnitude of the interquartile range was larger for dGM (Q3-Q1 = 0.5879) and WM (Q3-Q1 = 0.3989) than cGM (Q3-Q1 = 0.1183).

The linear correlation between *rV*(*t*) and MPM parameters across fixation time points (Figure 6) revealed that R_1_ had the strongest correlation with shrinkage (r from −0.859 to −0.777). R_2_* was also negatively correlated, albeit slightly less strongly (r from −0.712 to −0.656). MT_sat_ showed the weakest correlation (|r| < 0.325). These values indicate that both R_1_ and R_2_* followed similar shrinkage-related trends across specimens. The MT_sat_ correlations had small descriptive p-values in dGM and WM. However, visual inspection indicated that these associations were driven more by the offset between specimens rather than by within-specimen temporal trends (see color-coding). No correlation was found between *rV*(*t*) and N_A_ (data not shown).

## 4. Discussion

This study used densely sampled quantitative, whole-human-brain longitudinal MPM data, with an independent *in vivo* cohort as qualitative context, to characterize key changes across the *in vivo-to-in situ* transition, immersion fixation, and PBS immersion. Using these data, we quantified change-specific trends of R_1_, R_2_*, MT_sat_, and a PD proxy, tested which parsimonious temporal models generalized across specimens (using leave-one-out cross-validation), and linked long-fixation time R_1_ to tissue shrinkage, providing a practical bridge between *in vivo* MRI, fixed-tissue MRI, and histology.

### 4.1 Comparison of the MPM parameters during *in situ*, formaldehyde *ex situ*, and PBS *ex situ* conditions

We assessed MPM parameter changes across *in vivo*, *in situ,* formaldehyde immersion fixation, and PBS wash-out; by measuring five *postmortem* whole human brains and comparing them with a cohort of younger healthy *in vivo* subjects.

From *in vivo* to *in situ*, overall changes were modest except for an increase in white matter (WM) R_2_* and a decrease in both deep and cortical gray matter (dGM and cGM) MT_sat_. Confounding factors such as temperature (Birkl et al., 2014) and age difference between the *in vivo* and ex vivo groups (Callaghan et al., 2014) make interpretation of those findings challenging (see Limitations in Discussion §4.4).

From *in situ* through fixation, including the transition from *in situ* to first formaldehyde *ex situ* scan session, formaldehyde had the greatest impact on R_1_ and R_2_*. R_1_ increased from 247.55% (WM) to 351.08% (dGM) and R_2_* from 39.71% (dGM) to 59.54% (WM) consistent with previous reports (Birkl et al., 2016; Shatil et al., 2018; Shepherd et al., 2009; Yong-Hing et al., 2005). Although the saturation models provided empirical effective time constants, these should not be interpreted as evidence that stable plateau was observed. R_1_ and R_2_* continued increasing beyond 90 days, suggesting that a plateau was not reached within the robustly sampled period and might occur only after several months, consistent with prolonged R_1_ changes reported elsewhere (Raman et al., 2017). WM is dominated by densely packed myelinated axons, whereas cGM contains more neuronal cell bodies, dendrites and neuropil, and dGM has a greater contribution from iron. These compositional differences may contribute to tissue-specific fixation responses. In particular, the transient WM-MT_sat_ overshoot around days 10-11 during fixation may reflect myelin changes like the formation of vacuoles between myelin laminae and associated changes in water-macromolecule exchange (Seifert et al., 2019). N_A_ remained relatively stable, contrary to prior reports (Yong-Hing et al., 2005), suggesting that N_A_ is less sensitive to the present fixation protocol.

Immersion in PBS was associated with partial reversal of the fixation-effect on R_2_* increase, but also with a further increase in R_1_ and unchanged MT_sat_. Scan sessions of R_2_* in the surrounding solution (Supplementary Materials §S2, Figure S6) showed that PBS immersion reduced R_2_* by ∼ 6 s⁻¹, consistent with wash-out of residual fixative and with the observed decrease in tissue R_2_*. This agrees with previous reports of R_2_* reduction after PBS wash-out/rehydration (Dusek et al., 2019; Rivlin et al., 2014) and supports the interpretation that the R_2_* decrease is consistent with removal of residual formaldehyde reaction products. In contrast, the continued increase in R_1_ warrants further investigation.

### 4.2. Fixative modeling of the MPM parameters

Across parameters, the most robust temporal signatures of fixation were observed for R_1_ and R_2_*. Saturation-type models yielded stable estimates of saturation times for R_1_ and, to a lesser extent, for R_2_*. These saturation times were broadly comparable across tissue classes, with an unstable long-time component for WM R_2_*, indicating that these parameters capture consistent components of fixation progression at the whole-brain scale. In contrast, N_A_ exhibited no systematic temporal trend, and MT_sat_ time-constant estimates were less stable—particularly for the long saturation component—suggesting that MTsat is less suitable as a temporal marker of fixation stage under the present protocol. Conversely, its relative stability after the transient early overshoot and preservation of anatomical tissue contrast were advantageous for longitudinal image registration. MTsat was therefore used as the principal registration contrast, enabling visually robust spatial alignment across *in situ*, formaldehyde *ex situ*, and PBS *ex situ* states, despite the pronounced changes in R1 and R2* (Supplementary Materials §S1 and for visual inspection, see Supplementary Materials §S6). Thus, MTsat appears particularly useful as a comparatively stable anatomical contrast for longitudinal *postmortem* MRI.

To quantify robustness across donors, we assessed cross-specimen predictability using leave-one-out cross-validation. R_1_ trajectories were highly predictable across brains, R_2_* showed intermediate consistency, and MT_sat_ showed lower cross-specimen predictability. This pattern indicates that inter-subject variability is a dominant limiting factor for modeling MT_sat_ dynamics, whereas R_1_ provides the most generalizable temporal marker of fixation progression across brains in this dataset. Donor-specific factors such as *postmortem* delay, age, and microstructural differences may contribute to the cross-specimen variability and should be considered in future work when extending fixation models.

Model comparison revealed that the mono- and bi-exponential saturation models generally provided a better description of fixation-related dynamics than the reciprocal logistic model (Yong-Hing et al., 2005). These models worked particularly well for R_1_, R_2_*, and captured MT_sat_ in-sample, but showed lower cross-specimen predictability. For N_A_, the null model was preferred, supporting our observation that N_A_ estimates remained relatively stable during fixation. Among the nine fits, i.e. R_1_, R_2_*, and MT_sat_ for the three tissue-classes, five were best described by a mono-exponential form (WM- and dGM-R1, dGM-R2*, and cGM- and dGM-MTsat) and four by a bi-exponential form (cGM-R1, cGM- and WM-R2*, and WM-MTsat). Notably, the transient WM-MT_sat_ peak around days ∼10- 11 coincided with a transition in WM-R_2_* from a rapid to a slower increase, suggesting that different processes may dominate early versus late fixation. The saturation models remain phenomenological: they specify the form of s(t) in Methods §2.4 (Equations 2–3) but do not identify its source. Supplementary Materials §S7 (Equations S8–S11) shows that this additive form is compatible with a linear iron–myelin relaxivity model if concentrations of myelin and iron are taken as constant and the relaxivity changes are absorbed into a time-dependent component. Note that this interpretation is not unique and was not used for model selection.

Overall, R_1_ provides the most reproducible and generalizable temporal marker for harmonizing longitudinal *postmortem* qMRI across brains in our dataset.

### 4.3. Volumetric change of brain specimens due to fixation

Our findings confirm the qualitatively observed shrinkage of brain specimens reported by Birkl et al. (2016). For dGM, cGM, and WM, the fitted relative volume, rV(t), trend followed a smooth decay over fixation (∼4% over ∼150 days). The observed shrinkage could be explained by dehydration of the amino groups during the protein cross-linking reaction (Hoffman et al., 2015; Metz et al., 2004; Thavarajah et al., 2012). The magnitude of shrinkage agrees with the reported reduction of the water content after fixation (1-6%) in Birkl et al. (2016).

We found strong negative correlations between the relative volume change rV(t) and the change in R_1_, and weaker but comparable negative correlations with R_2_*. In contrast, MT_sat_ and N_A_ exhibited no systematic correlations with rV(t), consistent with their relative stability in our temporal analyses (Figure 6). In WM, the fitted relationship was R1=43.9/rV−42.7 s−1, whereas the Fatouros model predicts an R1–inverse-water-content coefficient of approximately 2.7 s−1 under the assumptions used for human WM. Thus, although the linear relationship is qualitatively consistent with the Fatouros model, the effective slope with respect to 1/rV is approximately one order of magnitude larger. This discrepancy is expected because 1/rV is not equivalent to 1/W, and because the Fatouros coefficient depends on the hydration fraction and relaxation properties of motionally restricted water. Both are likely altered during formaldehyde fixation through protein crosslinking, changes in hydration layers, and altered water–macromolecule exchange. The substantially larger R1 response relative to the small macroscopic volume change therefore suggests that fixation-associated water loss or redistribution alone cannot explain the observed R1 increase but likely acts together with fixation-induced changes in tissue relaxivity. The 1/rV analysis (Supplementary Materials §S5.1) addresses only this water-content contribution; it is distinct from the linear iron–myelin construction in Supplementary Materials §S7 (Equations S8–S11).

The comparison of individual trends with the heuristic model showed that cGM followed the fitted shrinkage pattern most closely, with high Kendall’s τ and low IQR, whereas dGM and WM were more variable. This suggests that dGM and WM respond more heterogeneously to immersion fixation, potentially reflecting differences in local microstructure, myelin content, or distance to fixative solution (i.e., deeper brain regions are later fixed). Moreover, we observed early WM expansion in some brains during fixation. The origins of these expansions, however, are still unclear to us.

We additionally performed an exploratory PBS-stage analysis (Supplementary Materials §S5.2), which showed continued volume reduction rather than reversal of fixation-associated shrinkage. However, PBS wash-out was not standardized across specimens, and whole-brain wash-out is slow and spatially heterogeneous. These findings should therefore be interpreted as changes under our experimental protocol rather than as an equilibrated PBS wash-out state.

Note that here we only investigate volume change during fixation, whereas the question about the initial volume change after brain extraction was not addressed. For further read on the latter topic, we refer to (Schulz et al., 2011).

### 4.4. Limitations and future directions

We analyzed five whole-brain *ex vivo* specimens and not all of them were scanned at each tissue condition. Although to our knowledge, this is the most densely sampled longitudinal *ex vivo* qMRI dataset to date, the sample remains small for robust statistical inference. Therefore, statistical results should be interpreted with appropriate caution. Nevertheless, our sample size was sufficient to identify preferred models for fixation-related temporal changes in MPM parameters and to characterize their predictive performance across specimens. Moreover, the main results from Figures 3 and 4 remained qualitatively consistent, when only three specimens with full sampling across all tissue conditions were investigated (Supplementary Materials §S4.5).

The *in vivo* data were used only as a qualitative reference for the *in situ* data, because several confounders like difference in tissue temperature, cohort’s age, scanner systems and *postmortem* tissue condition prohibit quantitative comparison. Published temperature coefficients suggest that cooling from body to room temperature may increase R1 and R2* (Supplementary Materials §S4.4), aging may decrease R1, particularly in WM, and increase R2* mainly in dGM and cGM (Supplementary Materials §S4.4). These estimates indicate that age and temperature are likely to contribute to some *in vivo*to-*in situ* offsets, but they do not explain the full tissue-class-specific pattern. Scanner-related differences between *in vivo*-to-*in situ* scans, however, were unlikely to introduce additional bias (Supplementary Materials §S4.2 and Figure S13).

Temperature was not directly monitored during scanning and was not included in the temporal saturation models. For the longitudinal ex vivo scan sessions, specimens were stored and scanned at approximately room temperature (∼22.5 °C). The estimated temperature effects due to variation in the room temperature or due to RF heating are substantially smaller than the observed fixation-related R1 and R2* changes (see Supplementary Materials §S4.3 and §S4.4). Moreover, because published MTsat temperature coefficients are lacking, and our unpublished pilot is not used as evidence here, temperature remains an unresolved confounder for MTsat; a temperature-only explanation of the observed fixation-related MTsat trajectory would require a systematic temperature drift across the entire fixation period. Temperature and the other confounders may thus contribute to residual variability, but they are unlikely to explain the main longitudinal trajectories, which should nevertheless be interpreted as empirical rather than pure fixation kinetics.

We used the same MPM protocol for *ex vivo* MRI that was optimized for *in vivo* MRI of healthy subjects (Helms et al., 2008; Weiskopf et al., 2013). To address potential biases associated with this choice, we conducted simulations to assess the influence of varying MPM parameters on the MPM signal and fitting (Supplementary Materials §S4.1). We found that the introduced bias was small (0-2%). The coefficient of variation, however, increased by up to 5 percentage points from 15% to 20% during the fixation process. In both cases, the effect was largest for MT_sat_. Similarly, the normalization of the “A” parameter (based on the fluid mask) could be an additional source of variance because the MPM protocol was not optimized for fluids.

We explored whether the first formaldehyde MPM values tracked PMI for four of the five specimens (Supplementary Materials §S4.6, Figure S17; Brain1 excluded). Most tissue-class–MPM parameter combinations showed no clear trend except for WM-R2* and, more weakly, WM-MTsat, in which they increased with PMI. Given the low number of samples, n = 4, these trends are descriptive only and do not support adjusting the fixation modeling with PMI. Effects of fixative concentration and PMI on the temporal evolution of the MPM parameters therefore remain insufficiently characterized; both may contribute to residual cross-specimen variability, especially in MTsat, and should be studied systematically. Future fixation models could incorporate those confounders explicitly.

An important prerequisite for the characterization of the quantitative MRI parameter changes from *in situ* to *ex vivo* is accurate longitudinal (deformable) registration of the brain specimens and robust segmentation of cGM, dGM and WM. Imprecision in either step can introduce additional noise into the measured longitudinal changes of MPM parameters. We assessed the quality of longitudinal registration and manually removed apparent outliers (for visual confirmation, see movies in Supplementary Materials §S6). To reduce residual effects of misregistration and other sources for temporal fluctuation (e.g., noise, instrumental imperfections, or variation in positioning), we assessed the median of each qMRI parameter per tissue class and used a single-time-point segmentation for all data points.

Accurate volume estimation is challenging and should be interpreted cautiously. For example, gentle contact with the holder may have altered local brain shape. However, because brain tissue is water-rich and approximately incompressible under small loads, such contact is expected to redistribute tissue rather than change its total volume. For an accurately registered volume-preserving deformation, regional Jacobian determinants may differ from one, but their whole-tissue average should remain approximately one. The observed global fixation-associated volume reduction is therefore unlikely to reflect mechanical compression by the holder, although local shape changes may contribute to registration uncertainty. Moreover, residual air bubbles may produce local susceptibility artifacts in FLASH images. Although these effects should be less pronounced at 3 T and in the MTsat map that was used for registration than at ultra-high field, subtle effects may remain and could contribute to registration uncertainty (for visual inspection see Supplementary Materials §S6).

Future work could investigate whether the proposed models are generalizable to other qMRI methods, smaller brain samples, other fixatives and by incorporating fixation-associated volume shrinkage.

## Conclusions

We used densely sampled longitudinal whole-human-brain MPM data, supplemented by an independent *in vivo* cohort, to quantify how R_1_, R_2_*, MT_sat_ and a PD proxy evolve across the *in vivo*–to–*in situ* transition, immersion fixation, and PBS wash-out. R_1_ and R_2_* showed the largest fixation-related changes. MT_sat_ showed an early fixation over- shoot but it became comparatively stable during fixation and PBS wash-out, preserving tissue contrast. This contrast stability made MT_sat_ suitable for spatial alignment between *in vivo* and *ex vivo* MRI, facilitating correspondence between MRI-derived microstructural patterns and histological readouts. Fixation-induced increases in R1 and relative tissue-volume loss evolved over similar timescales (up to 150 days). A mechanistically inspired inverse-volume analysis supported the hypothesis that decreased tissue volume associated with fixation-related water loss contributes to increasing R1, without establishing that shrinkage directly causes the R1 change. Leave-one-out analyses showed that R_1_ trajectories were highly predictable across brains, indicating that R_1_ was the most reproducible empirical marker of fixation progression under the present experimental conditions. Together, these findings provide practical guidance for *postmortem* MRI studies and strengthen the link between *in vivo* MRI, fixed-tissue imaging, and histology.

## Data and Code Availability

MRI data, both *in vivo* and *ex vivo*, are available from the corresponding authors under reasonable request.

The code used for pre-processing and analysis is available on Github at https://github.com/franciscofritz/LongitudinalFixation_MPM. At the moment of submission, the code is being refactored to improve preprocessing efficiency without changing the results reported in this manuscript.

After first revision, a new folder called “Revision” was uploaded to the same Github repository, containing all the code created for evaluating some of the confounders found by the reviewers.

This code also made use of other publicly available packages, such as Statistical Parametric Mapping (SPM 12, (Friston, 2007)) with the hMRI toolbox integrated (Tabelow et al., 2019), FMRIB Software Library (FSL, (Jenkinson et al., 2012)), FreeSurfer (Fischl, 2012) and Advanced Normalization Tools (ANTs, (Avants et al., 2008)).

## Author Contributions

**FJF:** MR data curation, investigation, formal analysis, visualization and writing of the original draft. **TS:** conceptualization and MR data acquisition. **LM**: review and editing. **NL:** review and editing. **LJE:** review and editing. **HM:** provided brain specimens. **KP:** provided brain specimens. **NW:** review and editing. **EK:** visualization, incorporation of extra analysis (tissue shrinkage), review and editing. **SM:** conceptualization, MR data acquisition and curation, resources, funding acquisition, review, main lead in paper writing and editing.

## Funding

The research leading to these results has received funding from the European Research Council under the European Union’s Seventh Framework Programme (FP7/2007-2013) / ERC grant agreement number 616905. This work was supported by the German Research Foundation (DFG Priority Program 2041 “Computational Connectomics”, [MO 2397/5-1, MO 2397/5-2, MO 2249/3-1, MO 2249/3-2], by the Emmy Noether Stipend: MO 2397/4-1; MO 2397/4-2) and the project no. 347592254 (WE 5046/4-2). Other stipends were ERA-NET NEURON (hMRI- ofSCI), the Bundesministerium für Bil- dung und Forschung (BMBF; 01EW1711A and B) and the Forschungszentrums Medizintechnik Hamburg (fmthh; grant 01fmthh2017)

Since this project was funded by the European Union, the views and opinions expressed are, however, those of the author(s) only and do not necessarily reflect those of the European Union or the European Research Council Executive Agency. Neither the European Union nor the granting authority can be held responsible for them. This work is supported by ERC grant (Acronym: MRStain, Grant agreement ID: 101089218, DOI: 10.3030/101089218).

## Declaration of Competing Interests

The Max Planck Institute for Human Cognitive and Brain Sciences and Wellcome Centre for Human Neuroimaging have institutional research agreements with Siemens Healthcare. NW holds a patent on acquisition of MRI data during spoiler gradients (US 10,401,453 B2).

## Supplementary Material

Two supplementary materials were added to this manuscript and are available online. A PDF document and 15 .avi format videos. These videos show the resulting longitudinal registration of the five brain specimens across all MPM parameters.

## Acknowledgments

We gratefully acknowledge Sophie Otte for sharing her MTsat temperature measurements, which contributed to better understanding of our results. We also acknowledge Carsten Jäger for providing the formaldehyde solution for all our brain specimens.

## Abbreviations

PBS: Phosphate buffered saline solution
In situ: Postmortem ex vivo in situ (i.e. dead brain retained in skull)
Formaldehyde *ex situ*: *Ex vivo ex situ* human brain immersed in formal-dehyde
PBS: *ex situ1Ex vivo ex situ* human brain immersed in PBS + 0.1% sodium azide

## Footnotes

1 The PBS solution used for dilution was composed of 1.43 g of Na_2_HPO_4_ (Roth 4984), 0.43 g of NaH_2_PO_4_ (Roth 2370) and 7.2 g of NaCl (Roth 3957), diluted in 1 L of water. The pH of this solution is 7.4. The same PBS solution was supplemented with 0.1% NaN_3_ for the PBS immersion phase.

## Notes

https://github.com/franciscofritz/LongitudinalFixation_MPM.git

### Summary of Updates

General: clearer, more precisely positioned within the current postmortem MRI literature, and more cautious in its interpretation. Revision: text, terminology, figures, and methodological descriptions. New analyses: we performed eight new quantitative analyses and sensitivity checks (in Supplementary Material). Found mistakes: Moreover, we found two mistakes in our original manuscript that we changed immediately. First, we corrected a nomenclature error regarding the fixative. In the original submission we referred to the fixative as paraformaldehyde (PFA), whereas the working solution was in fact prepared from a commercial acid-free 37% formaldehyde solution diluted in PBS to a final concentration of 4% formaldehyde. Paraformaldehyde is the polymeric solid from which formaldehyde solutions can be prepared, the active fixing agent is formaldehyde in either case. Because our solution was obtained by diluting aqueous formaldehyde rather than by depolymerizing paraformaldehyde powder, we now use "formaldehyde" throughout and have removed the acronym PFA. This change is terminological and does not affect experimental procedures, analyses, or results. Second, in our longitudinal fixation fitting (Methods section 2.4 and Results section 3.2), where we included the in situ time point for all the available specimens, while in Methods section 2.4 we mentioned that we only used the formaldehyde ex situ time points. We therefore refit using only the formaldehyde ex situ time points. Overall trends were unchanged; fitted parameters changed only slightly, and two preferred models changed: dGM-R2* and cGM-MTsat changed from bi- to mono-exponential saturation models.

## References

Ashburner, J., Ridgway, G.R., 2013. Symmetric Diffeomorphic Modeling of Longitudinal Structural MRI. Frontiers in Neuroscience Volume 6–2012. 10.3389/fnins.2012.00197

Avants, B.B., Epstein, C.L., Grossman, M., Gee, J.C., 2008. Symmetric diffeomorphic image registration with cross-correlation: Evaluating automated labeling of elderly and neurodegenerative brain. Medical Image Analysis, Special Issue on The Third International Workshop on Biomedical Image Registration – WBIR 2006 12, 26–41. 10.1016/j.media.2007.06.004

Bazin, P.-L., Alkemade, A., van der Zwaag, W., Caan, M., Mulder, M., Forstmann, B.U., 2019. Denoising High-Field Multi-Dimensional MRI With Local Complex PCA. Frontiers in Neuroscience Volume 13–2019. 10.3389/fnins.2019.01066

Beach, T.G., Tago, H., Nagai, T., Kimura, H., McGeer, P.L., McGeer, E.G., 1987. Perfusion-fixation of the human brain for immunohistochemistry: comparison with immersion-fixation. Journal of Neuroscience Methods 19, 183–192. 10.1016/S0165-0270(87)80001-8

Beaulieu-Laroche, L., Brown, N.J., Hansen, M., Toloza, E.H.S., Sharma, J., Williams, Z.M., Frosch, M.P., Cosgrove, G.R., Cash, S.S., Harnett, M.T., 2021. Allometric rules for mammalian cortical layer 5 neuron biophysics. Nature 600, 274–278. 10.1038/s41586-021-04072-3

Berger, C., Bauer, M., Scheurer, E., Lenz, C., 2023. Temperature correction of post mortem quantitative magnetic resonance imaging using real-time forehead temperature acquisitions. Forensic Science International 348, 111738. 10.1016/j.forsciint.2023.111738

Birkl, C., Langkammer, C., Golob-Schwarzl, N., Leoni, M., Haybaeck, J., Goessler, W., Fazekas, F., Ropele, S., 2016. Effects of formalin fixation and temperature on MR relaxation times in the human brain. NMR in Biomedicine 29, 458–465. 10.1002/nbm.3477

Birkl, C., Langkammer, C., Haybaeck, J., Ernst, C., Stollberger, R., Fazekas, F., Ropele, S., 2014. Temperature-induced changes of magnetic resonance relaxation times in the human brain: A postmortem study. Magnetic Resonance in Medicine 71, 1575–1580. 10.1002/mrm.24799

Birkl, C., Soellradl, M., Toeglhofer, A.M., Krassnig, S., Leoni, M., Pirpamer, L., Vorauer, T., Krenn, H., Haybaeck, J., Fazekas, F., Ropele, S., Langkammer, C., 2018. Effects of concentration and vendor specific composition of formalin on postmortem MRI of the human brain. Magnetic Resonance in Medicine 79, 1111–1115. 10.1002/mrm.26699

Brammerloh, M., Sibgatulin, R., Herrmann, K.-H., Morawski, M., Reinert, T., Jäger, C., Müller, R., Falkenberg, G., Brückner, D., Pine, K.J., Deistung, A., Kiselev, V.G., Reichenbach, J.R., Weiskopf, N., Kirilina, E., 2024. In Situ Magnetometry of Iron in Human Dopaminergic Neurons Using Superresolution MRI and Ion-Beam Microscopy. Phys. Rev. X 14, 021041. 10.1103/PhysRevX.14.021041

Burnham, K.P., Anderson, D.R., 2004. Multimodel Inference: Understanding AIC and BIC in Model Selection. Sociological Methods & Research 33, 261–304. 10.1177/0049124104268644

Burnham, K.P., Anderson, D.R., 2002. Model selection and multimodel inference: a practical information-theoretic approach. Springer Verlag.

Burnham, K.P., Anderson, D.R., Huyvaert, K.P., 2011. AIC model selection and multimodel inference in behavioral ecology: some background, observations, and comparisons. Behavioral Ecology and Sociobiology 65, 23–35. 10.1007/s00265-010-1029-6

Callaghan, M.F., Freund, P., Draganski, B., Anderson, E., Cappelletti, M., Chowdhury, R., Diedrichsen, J., FitzGerald, T.H.B., Smittenaar, P., Helms, G., Lutti, A., Weiskopf, N., 2014. Widespread age-related differences in the human brain microstructure revealed by quantitative magnetic resonance imaging. Neurobiology of Aging 35, 1862–1872. 10.1016/j.neurobiolaging.2014.02.008

David, G., Mohammadi, S., Martin, A.R., Cohen-Adad, J., Weiskopf, N., Thompson, A., Freund, P., 2019. Traumatic and nontraumatic spinal cord injury: pathological insights from neuroimaging. Nat Rev Neurol 15, 718–731. 10.1038/s41582-019-0270-5

Drayer, B., Burger, P., Darwin, R., Riederer, S., Herfkens, R., Johnson, G., 1986. Magnetic Resonance imaging of brain iron. American Journal of Neuroradiology 7, 373–380.

Dusek, P., Madai, V.I., Huelnhagen, T., Bahn, E., Matej, R., Sobesky, J., Niendorf, T., Acosta-Cabronero, J., Wuerfel, J., 2019. The choice of embedding media affects image quality, tissue R2*, and susceptibility behaviors in post-mortem brain MR microscopy at 7.0T. Magnetic Resonance in Medicine 81, 2688–2701. 10.1002/mrm.27595

Edwards, L., Bazin, P.L., Tabelow, K., Ugurcan, B.E., Mohammadi, S., Weiskopf, N., 2024. Denoising improves contrast while retaining sharpness of high resolution multiparameter R1, R2* and proton density maps. Magnetic Resonance Materials in Physics, Biology and Medicine 37, 1–781. 10.1007/s10334-024-01191-6

Ellerbrock, I., Mohammadi, S., 2018. Four in vivo g-ratio-weighted imaging methods: Comparability and repeatability at the group level. Hum Brain Mapp 39, 24–41. 10.1002/hbm.23858

Emmenegger, T.M., David, G., Ashtarayeh, M., Fritz, F.J., Ellerbrock, I., Helms, G., Balteau, E., Freund, P., Mohammadi, S., 2021. The Influence of Radio-Frequency Transmit Field Inhomogeneities on the Accuracy of G-ratio Weighted Imaging. Frontiers in Neuroscience 15, 770. 10.3389/fnins.2021.674719

Eriksson, S.H., Free, S.L., Thom, M., Martinian, L., Symms, M.R., Salmenpera, T.M., McEvoy, A.W., Harkness, W., Duncan, J.S., Sisodiya, S.M., 2007. Correlation of quantitative MRI and neuropathology in epilepsy surgical resection specimens— T2 correlates with neuronal tissue in gray matter. NeuroImage 37, 48–55. 10.1016/j.neuroimage.2007.04.051

Fischl, B., 2012. FreeSurfer. NeuroImage, 20 YEARS OF fMRI 62, 774–781. 10.1016/j.neuroimage.2012.01.021

Freund, P., Seif, M., Weiskopf, N., Friston, K., Fehlings, M.G., Thompson, A.J., Curt, A., 2019. MRI in traumatic spinal cord injury: from clinical assessment to neuroimaging biomarkers. The Lancet Neurology 18, 1123–1135. 10.1016/S1474-4422(19)30138-3

Freund, P., Weiskopf, N., Ashburner, J., Wolf, K., Sutter, R., Altmann, D.R., Friston, K., Thompson, A., Curt, A., 2013. MRI investigation of the sensorimotor cortex and the corticospinal tract after acute spinal cord injury: a prospective longitudinal study. The Lancet Neurology 12, 873–881. 10.1016/S1474-4422(13)70146-7

Friston, K., 2007. CHAPTER 2 Statistical parametric mapping, in: Friston, K., Ash-Burner, J., Kiebel, S., Nichols, T., Penny, W. (Eds.), Statistical Parametric Mapping. Academic Press, London, pp. 10–31. 10.1016/B978-012372560-8/50002-4

Germann, J., Gouveia, F.V., Chakravarty, M.M., Devenyi, G.A., 2025. Longitudinal deformation-based morphometry pipeline to study neuroanatomical differences in structural MRI based on SyN unbiased templates. Aperture Neuro 5. 10.52294/001c.133510

Ghosh, J.K., Delampady, M., Samanta, T., 2007. An Introduction to Bayesian Analysis: Theory and Methods, Springer Texts in Statistics. Springer New York.

Helms, G., Dathe, H., Dechent, P., 2008. Quantitative FLASH MRI at 3T using a rational approximation of the Ernst equation. Magnetic Resonance in Medicine 59, 667– 672. 10.1002/mrm.21542

Hoffman, E.A., Frey, B.L., Smith, L.M., Auble, D.T., 2015. Formaldehyde Crosslinking: A Tool for the Study of Chromatin Complexes*. Journal of Biological Chemistry 290, 26404–26411. 10.1074/jbc.R115.651679

Insausti, R., Insausti, A.M., Muñoz López, M., Medina Lorenzo, I., Arroyo-Jiménez, M.D.M., Marcos Rabal, M.P., de la Rosa-Prieto, C., Delgado-González, J.C., Montón Etxeberria, J., Cebada-Sánchez, S., Raspeño-García, J.F., Iñiguez de Onzoño, M.M., Molina Romero, F.J., Benavides-Piccione, R., Tapia-González, S., Wisse, L.E.M., Ravikumar, S., Wolk, D.A., DeFelipe, J., Yushkevich, P., Artacho-Pérula, E., 2023. Ex vivo, in situ perfusion protocol for human brain fixation compatible with microscopy, MRI techniques, and anatomical studies. Front Neuroanat 17, 1149674. 10.3389/fnana.2023.1149674

Jenkinson, M., Beckmann, C.F., Behrens, T.E.J., Woolrich, M.W., Smith, S.M., 2012. FSL. NeuroImage 62, 782–790.

Kendall, M.G., 1938. A new measure of rank correlation. Biometrika 30, 81–93. 10.1093/biomet/30.1-2.81

Kirilina, E., Helbling, S., Morawski, M., Pine, K., Reimann, K., Jankuhn, S., Dinse, J., Deistung, A., Reichenbach, J.R., Trampel, R., Geyer, S., Müller, L., Jakubowski, N., Arendt, T., Bazin, P.-L., Weiskopf, N., 2020. Superficial white matter imaging: Contrast mechanisms and whole-brain in vivo mapping. Science Advances 6, eaaz9281. 10.1126/sciadv.aaz9281

Kjaer, L., Henriksen, O., 1988. Comparison of different pulse sequences for in vivo determination of T1 relaxation times in the human brain. Acta Radiol 29, 231–236.

Krassner, M.M., Kauffman, J., Sowa, A., Cialowicz, K., Walsh, S., Farrell, K., Crary, J.F., McKenzie, A.T., 2023. Postmortem changes in brain cell structure: a review. Free Neuropathol 4, 4–10. 10.17879/freeneuropathology-2023-4790

Lehmann, N., Aye, N., Kaufmann, J., Heinze, H.-J., Düzel, E., Ziegler, G., Taubert, M., 2023. Changes in Cortical Microstructure of the Human Brain Resulting from Long-Term Motor Learning. J. Neurosci. 43, 8637–8648. 10.1523/JNEUROSCI.0537-23.2023

Leutritz, T., Seif, M., Helms, G., Samson, R.S., Curt, A., Freund, P., Weiskopf, N., 2020. Multiparameter mapping of relaxation (R1, R2*), proton density and magnetization transfer saturation at 3 T: A multicenter dual-vendor reproducibility and repeatability study. Human Brain Mapping 41, 4232–4247. 10.1002/hbm.25122

Lutti, A., Stadler, J., Josephs, O., Windischberger, C., Speck, O., Bernarding, J., Hutton, C., Weiskopf, N., 2012. Robust and fast whole brain mapping of the RF transmit field B1 at 7T. PLoS One 7, e32379–e32379. 10.1371/journal.pone.0032379

Maranzano, J., Dadar, M., Bertrand-Grenier, A., Frigon, E.-M., Pellerin, J., Plante, S., Duchesne, S., Tardif, C.L., Boire, D., Bronchti, G., 2020. A novel *ex vivo, in situ* method to study the human brain through MRI and histology. Journal of Neuroscience Methods 345, 108903. 10.1016/j.jneumeth.2020.108903

McFadden, W.C., Walsh, H., Richter, F., Soudant, C., Bryce, C.H., Hof, P.R., Fowkes, M., Crary, J.F., McKenzie, A.T., 2019. Perfusion fixation in brain banking: a systematic review. acta neuropathol commun 7, 146. 10.1186/s40478-019-0799-y

Metz, B., Kersten, G.F.A., Hoogerhout, P., Brugghe, H.F., Timmermans, H.A.M., Jong, A. de, Meiring, H., Hove, J. ten, Hennink, W.E., Crommelin, D.J.A., Jiskoot, W., 2004. Identification of Formaldehyde-induced Modifications in Proteins: REACTIONS WITH MODEL PEPTIDES*. Journal of Biological Chemistry 279, 6235– 6243. 10.1074/jbc.M310752200

Mezer, A., Rokem, A., Berman, S., Hastie, T., Wandell, B.A., 2016. Evaluating quantitative proton-density-mapping methods. Hum Brain Mapp 37, 3623–3635. 10.1002/hbm.23264

Mohammadi, S., Streubel, T., Klock, L., Edwards, L.J., Lutti, A., Pine, K.J., Weber, S., Scheibe, P., Ziegler, G., Gallinat, J., Kühn, S., Callaghan, M.F., Weiskopf, N., Tabelow, K., 2022. Error quantification in multi-parameter mapping facilitates robust estimation and enhanced group level sensitivity. Neuroimage 262, 119529. 10.1016/j.neuroimage.2022.119529

Molinaro, A.M., Simon, R., Pfeiffer, R.M., 2005. Prediction error estimation: a comparison of resampling methods. Bioinformatics 21, 3301–3307. 10.1093/bioinformatics/bti499

Mordhorst, L., Edwards, L.J., Morozova, M., Ashtarayeh, M., Streubel, T., Fricke, B., Fritz, F.J., Rusch, H., Jäger, C., Starke, L., Gladytz, T., Tasbihi, E., Periquito, J.S., Pohlmann, A., Mushumba, H., Püschel, K., Niendorf, T., Weiskopf, N., Morawski, M., Mohammadi, S., 2025. MRI-scale histology validates spatial sensitivity of in-vivo MRI-based axon radius estimation. Imaging Neuroscience 3, IMAG.a.1030. 10.1162/IMAG.a.1030

Neuhaus, D., Rost, T., Haas, T., Wendebourg, M.J., Schulze, K., Schlaeger, R., Scheurer, E., Lenz, C., 2025. Comparative analysis of in situ and ex situ postmortem brain MRI: Evaluating volumetry, DTI, and relaxometry. Magnetic Resonance in Medicine 93, 213–227. 10.1002/mrm.30264

Papazoglou, S., Ashtarayeh, M., Oeschger, J.M., Callaghan, M.F., Does, M.D., Mohammadi, S., 2024. Insights and improvements in correspondence between axonal volume fraction measured with diffusion-weighted MRI and electron microscopy. NMR in Biomedicine 37, e5070. 10.1002/nbm.5070

Quester, R., Schröder, R., 1997. The shrinkage of the human brain stem during formalin fixation and embedding in paraffin. Journal of Neuroscience Methods 75, 81–89. 10.1016/S0165-0270(97)00050-2

Raman, M.R., Shu, Y., Lesnick, T.G., Jack, C.R., Kantarci, K., 2017. Regional T1 relaxation time constants in Ex vivo human brain: Longitudinal effects of formalin exposure. Magnetic Resonance in Medicine 77, 774–778. 10.1002/mrm.26140

Rivlin, M., Eliav, U., Navon, G., 2014. NMR studies of proton exchange kinetics in aqueous formaldehyde solutions. Journal of Magnetic Resonance 242, 107–112. 10.1016/j.jmr.2014.02.021

Schulz, G., Crooijmans, H.J.A., Germann, M., Scheffler, K., Müller-Gerbl, M., Müller, B., 2011. Three-dimensional strain fields in human brain resulting from formalin fixation. Journal of Neuroscience Methods 202, 17–27. 10.1016/j.jneumeth.2011.08.031

Seifert, A.C., Umphlett, M., Hefti, M., Fowkes, M., Xu, J., 2019. Formalin tissue fixation biases myelin-sensitive MRI. Magnetic Resonance in Medicine 82, 1504–1517. 10.1002/mrm.27821

Shatil, A.S., Uddin, M.N., Matsuda, K.M., Figley, C.R., 2018. Quantitative Ex Vivo MRI Changes due to Progressive Formalin Fixation in Whole Human Brain Specimens: Longitudinal Characterization of Diffusion, Relaxometry, and Myelin Water Fraction Measurements at 3T. Frontiers in Medicine 5.

Shepherd, T.M., Flint, J.J., Thelwall, P.E., Stanisz, G.J., Mareci, T.H., Yachnis, A.T., Blackband, S.J., 2009. Postmortem interval alters the water relaxation and diffusion properties of rat nervous tissue–implications for MRI studies of human autopsy samples. Neuroimage 44, 820–826.

Stüber, C., Morawski, M., Schäfer, A., Labadie, C., Wähnert, M., Leuze, C., Streicher, M., Barapatre, N., Reimann, K., Geyer, S., Spemann, D., Turner, R., 2014. Myelin and iron concentration in the human brain: A quantitative study of MRI contrast. NeuroImage 93, 95–106. 10.1016/j.neuroimage.2014.02.026

Tabelow, K., Balteau, E., Ashburner, J., Callaghan, M.F., Draganski, B., Helms, G., Kherif, F., Leutritz, T., Lutti, A., Phillips, C., Reimer, E., Ruthotto, L., Seif, M., Weiskopf, N., Ziegler, G., Mohammadi, S., 2019. hMRI – A toolbox for quantitative MRI in neuroscience and clinical research. NeuroImage 194, 191–210. 10.1016/j.neuroimage.2019.01.029

Tendler, B.C., Qi, F., Foxley, S., Pallebage-Gamarallage, M., Menke, R.A.L., Ansorge, O., Hurley, S.A., Miller, K.L., 2021. A method to remove the influence of fixative concentration on postmortem T2 maps using a kinetic tensor model. Human Brain Mapping 42, 5956–5972. 10.1002/hbm.25661

Thavarajah, R., Mudimbaimannar, V.K., Elizabeth, J., Rao, U.K., Ranganathan, K., 2012. Chemical and physical basics of routine formaldehyde fixation. J Oral Maxillofac Pathol 16, 400–405. 10.4103/0973-029X.102496

van Duijn, S., Nabuurs, R.J.A., van Rooden, S., Maat-Schieman, M.L.C., van Duinen, S.G., van Buchem, M.A., van der Weerd, L., Natté, R., 2011. MRI artifacts in human brain tissue after prolonged formalin storage. Magn Reson Med 65, 1750– 1758. 10.1002/mrm.22758

Weiskopf, N., Callaghan, M.F., Josephs, O., Lutti, A., Mohammadi, S., 2014. Estimating the apparent transverse relaxation time (R2*) from images with different contrasts (ESTATICS) reduces motion artifacts. Frontiers in Neuroscience 8.

Weiskopf, N., Edwards, L.J., Helms, G., Mohammadi, S., Kirilina, E., 2021. Quantitative magnetic resonance imaging of brain anatomy and in vivo histology. Nat Rev Phys 3, 570–588. 10.1038/s42254-021-00326-1

Weiskopf, N., Suckling, J., Williams, G., Correia, M.M., Inkster, B., Tait, R., Ooi, C., Bullmore, E.T., Lutti, A., 2013. Quantitative multi-parameter mapping of R1, PD(*), MT, and R2(*) at 3T: a multi-center validation. Front Neurosci 7, 95.

Whitaker, K.J., Vértes, P.E., Romero-Garcia, R., Váša, F., Moutoussis, M., Prabhu, G., Weiskopf, N., Callaghan, M.F., Wagstyl, K., Rittman, T., Tait, R., Ooi, C., Suckling, J., Inkster, B., Fonagy, P., Dolan, R.J., Jones, P.B., Goodyer, I.M., NSPN Consortium, Bullmore, E.T., 2016. Adolescence is associated with genomically patterned consolidation of the hubs of the human brain connectome. Proc Natl Acad Sci U S A 113, 9105–9110. 10.1073/pnas.1601745113

Yong-Hing, C.J., Obenaus, A., Stryker, R., Tong, K., Sarty, G.E., 2005. Magnetic resonance imaging and mathematical modeling of progressive formalin fixation of the human brain. Magnetic Resonance in Medicine 54, 324–332. 10.1002/mrm.20578

