## Supplemental_Text for "Longitudinal quantitative MRI of whole human brains during formaldehyde immersion, PBS wash-out, and tissue volume change"

### Supplementary material

#### S1. MR registration and preprocessing, including the bubble masking for the *ex vivo* human brain specimens

The *ex vivo* MRI datasets, i.e. *in situ*, formaldehyde *ex situ* and PBS *ex situ*, underwent denoising (previous MPM analysis), manual realignment, threshold-based masking, bias field correction, registration, and segmentation. Each step is below.

Denoising: To minimize the effect of noise and bias in the calculation of the MPM parameter maps due to reduced SNR, all MR images, i.e. 16 PDw-, 16 $T_1$w- and 12 MTw-magnitude images, were denoised using the local complex PCA (LCPCA) approach (Bazin et al., 2019; Edwards et al., 2024). The LCPCA default setup was used: neighborhood size of 4 and standard deviation cut-off of 1.05. We used the implementation available in the hMRI toolbox (Tabelow et al., 2019).

Manual realignment: Each brain was rigidly aligned to the *in situ* time point when available, otherwise the first formaldehyde *ex situ* scan. We performed this alignment in 3D Slicer software (Fedorov et al., 2012) to correct large field-of-view misalignments and improve the subsequent automated registration.

Thresholding and bias field correction: The $MT_{sat}$ maps were thresholded to the range 0-5 p.u., which means that all values outside this range were replaced with 0 p.u. This step reduces extreme outliers and improves the stability of subsequent registration. Later, the thresholded $MT_{sat}$ maps were first segmented using *mri_synthseg* from FreeSurfer version 7.4 (Billot et al., 2023). We created a brain mask by combining all estimated tissue segments and applied it to the $MT_{sat}$ and A maps. We then estimated a bias field from the masked A map at each time point and applied the correction to the unmasked A map.

Registration: two different registration processes were performed for the analyses of MPM parameters and estimation of shrinkage.

The first approach consisted of creating a non-linear spatially registered template across all the formaldehyde and PBS *ex situ* brain-masked $MT_{sat}$ maps using the optimized version of the ANTs command *antsMultivariateTemplateConstruction2.sh* (https://github.com/CoBrALab/optimized_antsMultivariateTemplateConstruction) and then registering the *in situ* time point to the template. Later, a composite transform was applied to the n-th time point to register it with respect to the *in situ* or 1$^{st}$ fixative time point, i.e. GenericAffine_nTimePoint -> Warp_nTimePoint -> InverseWarp_InSituTimePoint -> Inverse [GenericAffine_InSituTimePoint]. The *in situ* time point was not included in the template construction, instead, the formaldehyde + PBS *ex situ* template was built from *ex vivo* scans, and the template was registered to the *in situ* time point to define a common reference. This decision was taken due to the anatomical deformations which occurred during excision and manipulation. The final maps were used for analysis 1 and 2.

If the optimized pipeline fails, as occurred during reproducibility test, the following ANTs commands can be used instead:

```
antsMultivariateTemplateConstruction2.sh -d 3 -c 2 -j 36 -i 2 -n 0 -o anLR_ -r 1 -t BSplineSyN -z <InSitu image> input_list.txt
```

where input_list.txt is the file text containing all the images to be deformed; and then

```
antsRegistrationSyN.sh -d 3 -f ../intermediateTemplates/BSplineSyN_iteration1_anLR_template0.nii.gz -m <InSitu_image> -r 4 -g 0.4 -t b -s 13 -j 1 -o <InSitu_deformed_image>
```

The second command registers the *in situ* image to the template. It may also be rerun for an input image if visual quality control identifies an incorrect deformation.

The second approach was based on SPM. We used the auto-realign module in the hMRI toolbox to bring the specimens into MNI space for better segmentation. Then, the $MT_{sat}$ and A contrasts were used to estimate tissue probability maps and construct a brain mask for each time point. Afterwards, the longitudinal registration module in SPM was used to estimate the non-linear transformations per time point (Ashburner and Ridgway, 2013). To this end, the masked $MT_{sat}$ maps were resampled to a voxel size of 1.5 mm isotropic and used as source images. For each masked $MT_{sat}$ map a deformation field was obtained.

Jacobian composition (only for §2.5): To estimate the Jacobian determinant used for the relative volume change analysis, we composed the deformation field from the fixation time point *t* to the mean template with the inverse deformation field from the reference time point $t_0$ (first formaldehyde *ex situ* scan session) to the mean template. This composition yields the total deformation from any fixation time point *t* to reference, from which the Jacobian determinant is calculated.

Let $g$ map a voxel from a fixation time point to template space and let $f$ map the voxel from template space to the reference space. The Jacobian of the composite transformation is therefore:

$$J_{fog}(i) = J_f\big(g(i)\big)J_g(i) \tag{S1}$$

Here, $J_f(g(i))$ is the Jacobian of the deformation field that goes from the template to the first formaldehyde *ex situ* scan session space, whereas $J_g(i)$ is the Jacobian of the deformation field that transforms any i-voxel from any fixation time point to template. This equivalence is demonstrated through the multivariate chain-rule. This proof supports the diagram shown in Figure 2A.

Segmentation: Either the *in situ* or first formaldehyde time $MT_{sat}$ images were used to create the tissue classes used for all the analyses: cerebrospinal fluid ("fluid mask"), cortical gray matter (cGM), deep gray matter (dGM), and white matter (WM), using "spm_segment". The resulting segmentation masks were manually inspected and corrected (e.g., to add missed voxels and remove outliers). For characterizing temporal changes in MPM parameters and the tissue shrinkage, tissue probability maps were thresholded between 0.5 and 0.9 in steps of 0.1. The results reported in Results of the main manuscript used a threshold value of 0.8.

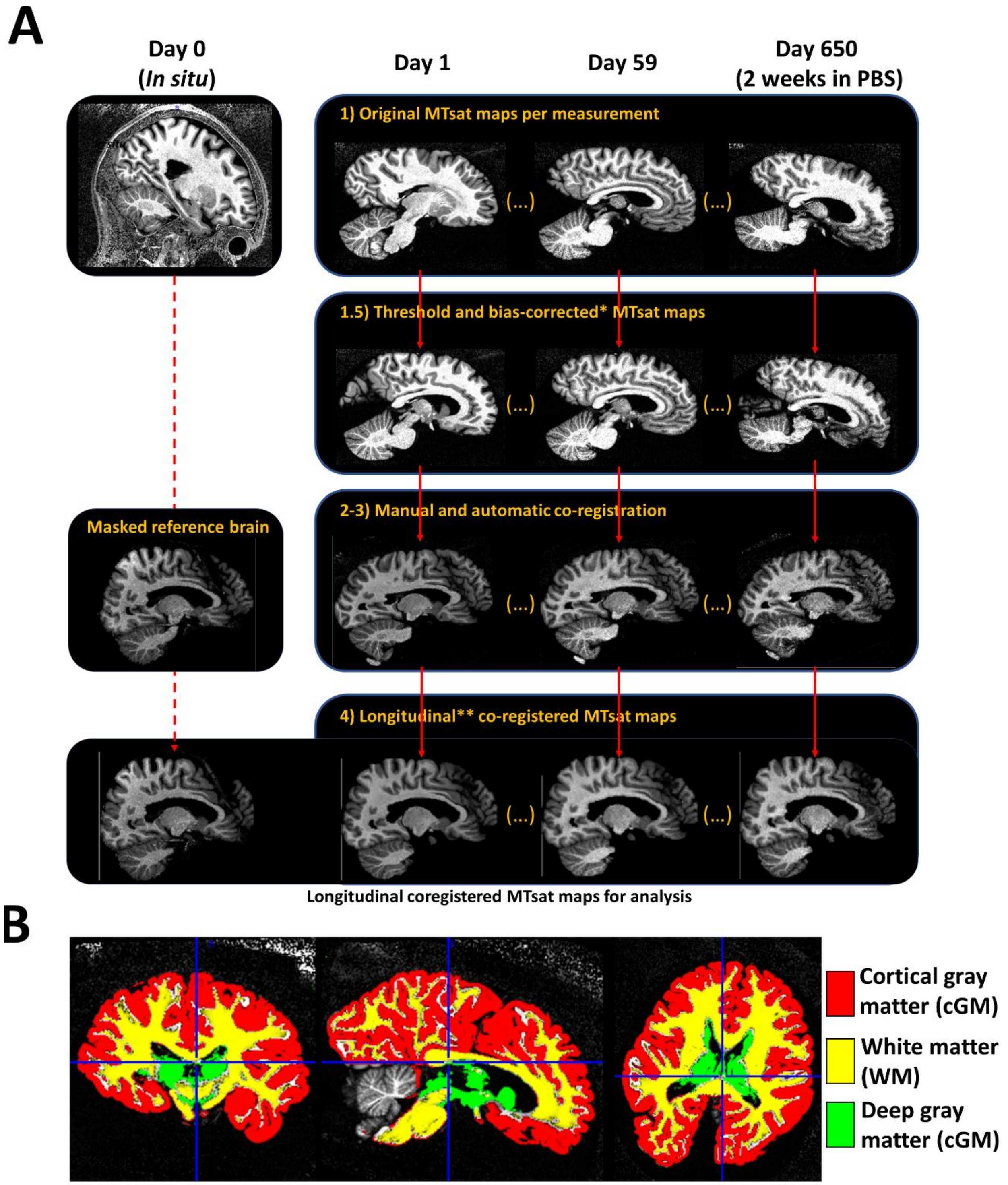


*Figure S1: (A) Illustration of the longitudinal registration pipeline for the MPM parameters of one brain specimen (Brain 6). Across the pipeline, only the $MT_{sat}$ images were used, and each estimated transformation applied to the remaining MPM parameters. The pipeline is shown in four stages. (1) The original in situ image and representative ex vivo $MT_{sat}$ volumes are shown before registration. (1.5) The MPM maps are thresholded; bias-field correction is applied only to A. (2–3) The ex vivo $MT_{sat}$ maps undergo rigid and automated registration to the brain-masked in situ image. (4) The final panel shows the longitudinally registered in situ and ex vivo $MT_{sat}$ maps. The present study used ANTs for the automated registration step. *Bias-field correction was only applied to the A maps. **In this step, either SPM or ANTs software can be used. In this work we used ANTs. (B) Illustration of the regions of interest (ROIs) used for analysis. Brain tissue*

*classes covered the cortical (red) and deep (green) gray matter, and white matter (yellow). The illustrated masks represent the resulting 90% threshold applied to the corresponding tissue probability masks (TPMs).*

<u>Bubble masking:</u> In our study, we enclosed *postmortem* human brain tissue in containers filled with solutions without using a vacuum device, leading to the formation of unwanted air bubbles. Because air bubbles exhibit a different magnetic susceptibility compared to the brain tissue, they strongly affect the estimation of $R_2^*$ (Alexander et al., 1996) and can introduce bias. To mitigate this, we created for each measurement a mask that excludes voxels containing air bubbles using the estimated $\epsilon$ maps. $\epsilon$ maps were thresholded into three categories using the Otsu's method (Otsu, 1979): background, $R_2^*$ error fitting in normal tissue, and $R_2^*$ error altered by air bubbles (with multithresh function in Matlab). In this estimation, two thresholds are estimated (using a search-based method) on the aggregate histogram for the entire image. The highest threshold value divides the high-intensity $R_2^*$ error values from the bubbles and the $R_2^*$ error values from tissue, while the lowest threshold divides the $R_2^*$ error values from tissue and background. The former was used to create a (complementary) brain mask, hereafter referred to as bubble mask (brain without air bubbles).

For our study, the contribution of air bubbles in comparison to the estimated tissue classes, i.e., dGM, WM and cGM; and the fluid mask in percentage of voxels were less than 10% as shown in Figure S2 across all formaldehyde (dark blue dots) and PBS (light blue dots) *ex situ* scan sessions per specimen (range 0-9.15%). If we consider only the tissue classes, the median volume contribution due to air bubbles was 1.55% (-0.481 p.p.), 1.70% (-1.177 p.p.), 0.86% (-0.767 p.p.), 0.89% (-0.587 p.p.) and 0.40% (-0.927 p.p.) for Brain1 to Brain6, respectively[1].

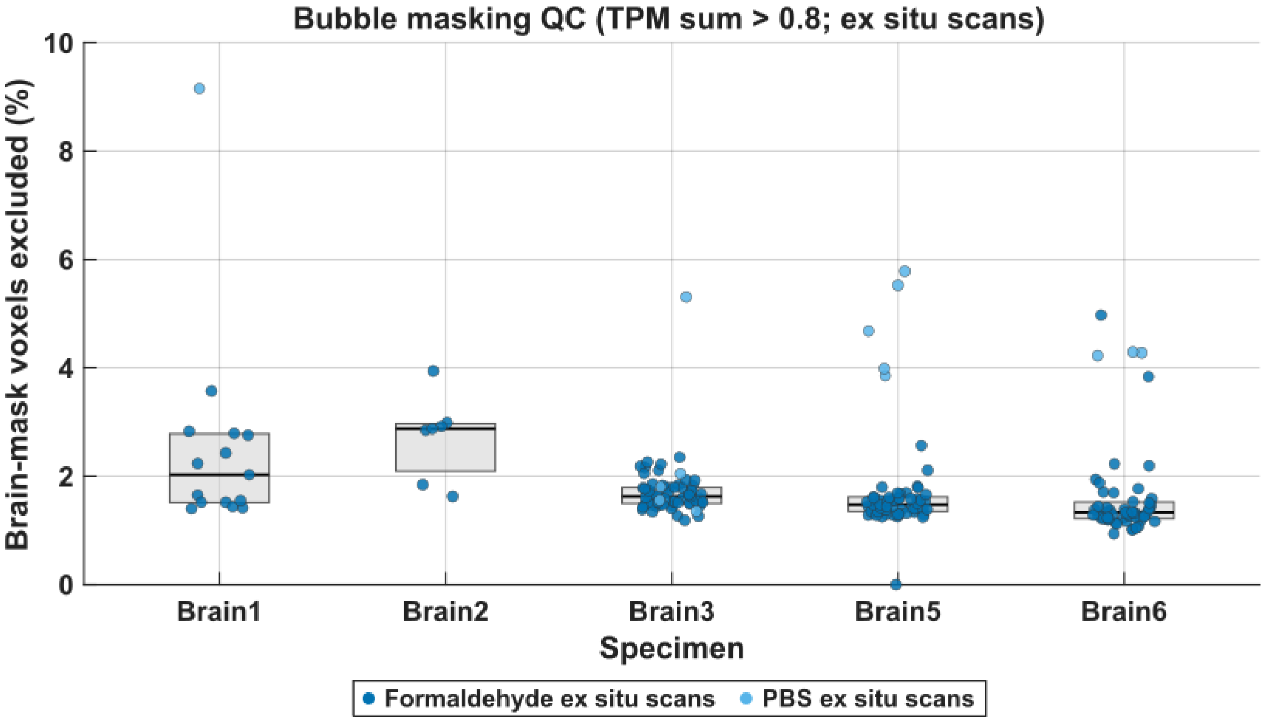


*Figure S2: Percentage of brain-mask voxels excluded by bubble masking (TPM sum > 0.8) for each formaldehyde and PBS ex situ session. Boxes show the median and interquartile range per specimen; points are individual sessions (dark*

[1] P.p. refers to percentage points. In this case, this difference is given between the volume % using 4 tissue classes and the % using three tissue classes.

*blue: formaldehyde; light blue: PBS). Across n = 202 sessions, the median exclusion was 1.53% (range 0–9.15%). Specimen medians were 2.03% (Brain1), 2.88% (Brain2), 1.63% (Brain3), 1.48% (Brain5), and 1.33% (Brain6).*

To illustrate the bubble-masking procedure, we used two figures (Figure S3 and S4). Figure S3 shows MPM parameter maps with and without the identified bubble-affected voxels for one scan session per specimen with a relatively high percentage of voxels classified as bubble affected. It also compares the median MPM parameter values within each tissue class before and after removal of these voxels. Figure S4 presents the same analysis for one scan session per specimen with a relatively low percentage of bubble-affected voxels. The maximum difference in median MPM parameters between analyses with and without bubble masking was larger when more bubble-affected voxels were identified (up to 1.93%; Figure S3) than when fewer bubble-affected voxels were identified (up to 0.35%; Figure S4). These results indicate that bubble masking reduces bias in the estimated MPM parameters. However, the magnitude of this effect was small compared with the parameter changes observed across tissue stages and during longitudinal fixation. Thus, bubble-related bias is unlikely to account for the main longitudinal MPM trajectories reported in this study.

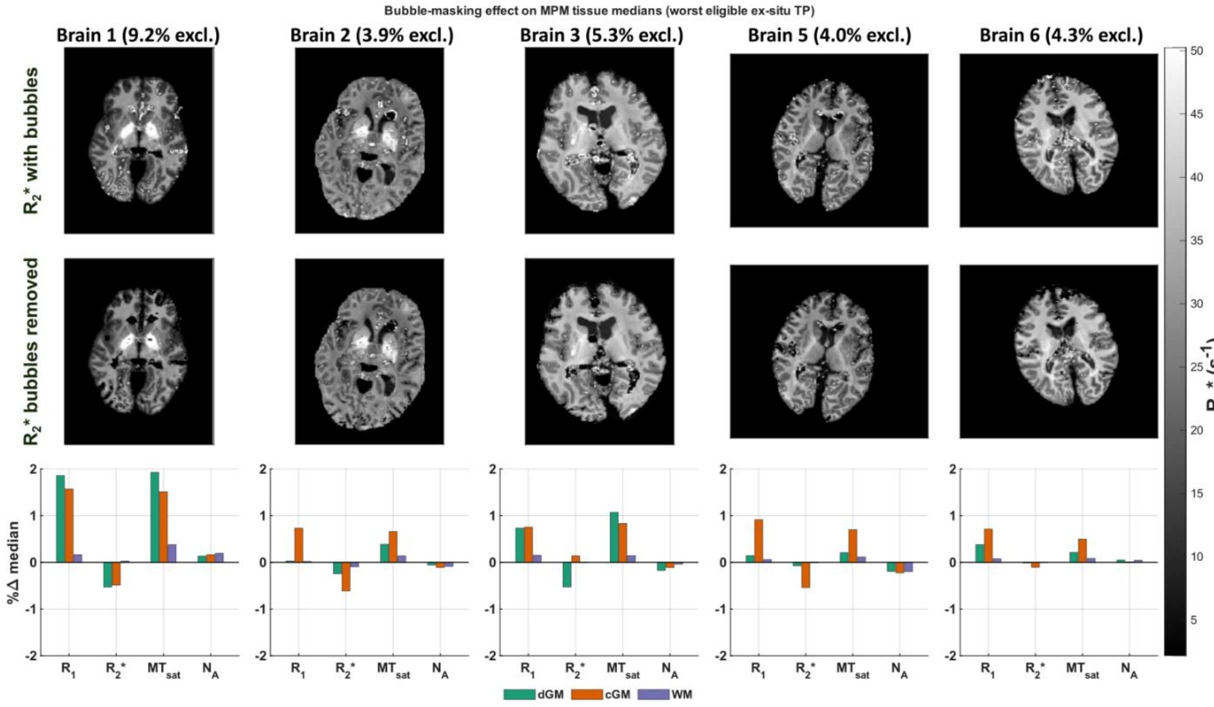


*Figure S3: Bubble-masking effect for the highest eligible ex situ time point per specimen (Brain1: 9.2%; Brain2: 3.9%; Brain3: 5.3%; Brain5: 4.0%; Brain6: 4.3%). Top: $R_2$* with bubble-affected voxels retained. Middle: same slice after bubble removal. Bottom: % change in tissue medians after masking, (%Δ) = (median_after / median_before − 1) × 100, for $R_1$, $R_2$*, $MT_{sat}$, and $N_a$ ($N_a$ = median(A_tissue)/median(A_CSF)) in dGM, cGM, and WM. Absolute median shifts remained within about ±2% (maximum |%Δ| = 1.93%). Color bar: $R_2$* ($s^{-1}$).*

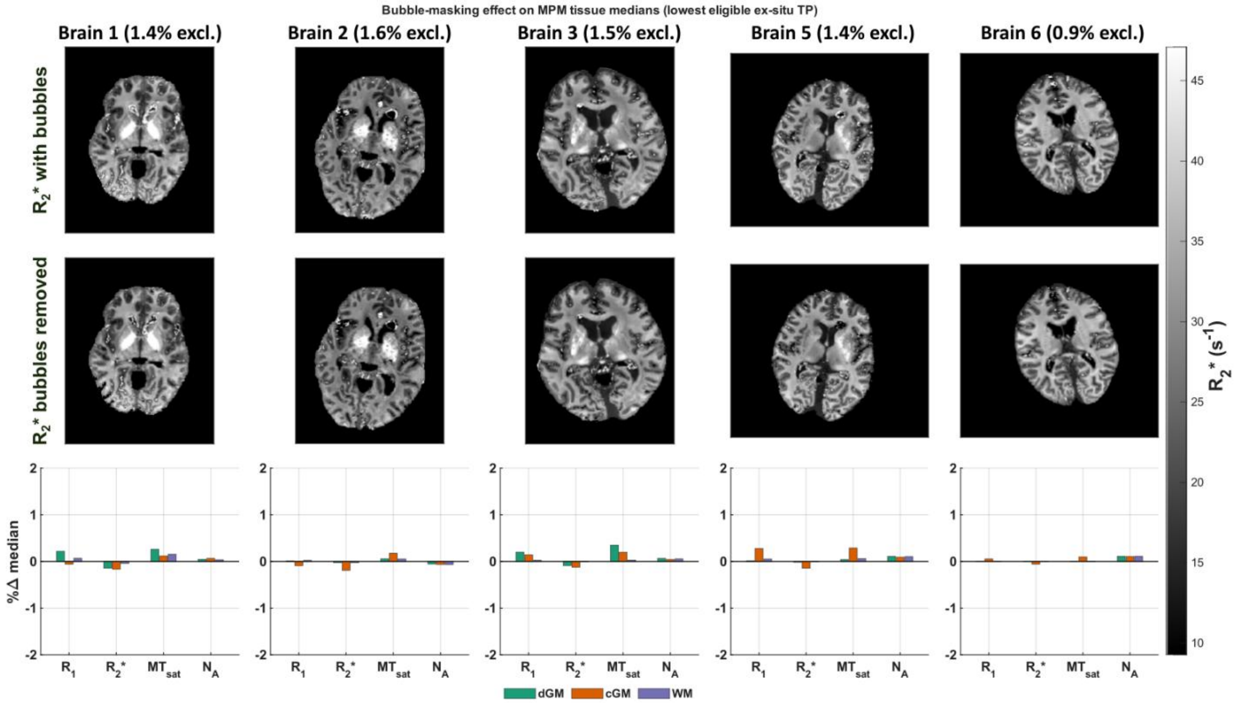


*Figure S4: Bubble-masking effect for the lowest eligible ex situ time point per specimen (Brain1: 1.4%; Brain2: 1.6%; Brain3: 1.5%; Brain5: 1.4%; Brain6: 0.9%). The layout is the same as presented in Figure S3. Absolute median shifts remained within about ±0.4% (maximum |%Δ| = 0.35%). Color bar: $R_2$* ($s^{-1}$).*

### S2. Comparison between the last formaldehyde *ex situ* scan and PBS *ex situ* scan per specimen

To characterize the mean change between consecutive *postmortem* processes, we estimated the difference in the MPM parameters per *postmortem* process. We report two complementary comparisons: (i) a common-time-point comparison using day 90 (available across specimens), and (ii) a specimen-specific comparison between each specimen's final formaldehyde *ex situ* scan (ranged from ~92 to 603 days) and its PBS *ex situ* scan (ranged from 32 to approximately 200 days).

To this end, the relative mean changes between the last formaldehyde and PBS *ex situ* scans were first calculated per specimen and then averaged across specimens, i.e., $\Delta MPM_{hyd,last-fA} = \frac{1}{4}\sum_i^4\left(\frac{MPM_{i,hyd}}{MPM_{i,last-fA}} - 1\right)$. This approach was used because the last point of fixation was not only specimen-dependent but also time-dependent, from 92 to 603 days. This estimation allowed us to assess the difference of at least 7 days of PBS immersion after different fixation times.

Figure S5 shows the change between the last formaldehyde *ex situ* and PBS *ex situ* per specimen, where each MPM parameter followed a different trend: $R_1$ increased, $R_2$* decreased, and both $MT_{sat}$ and $N_A$ (not shown) remained stable.

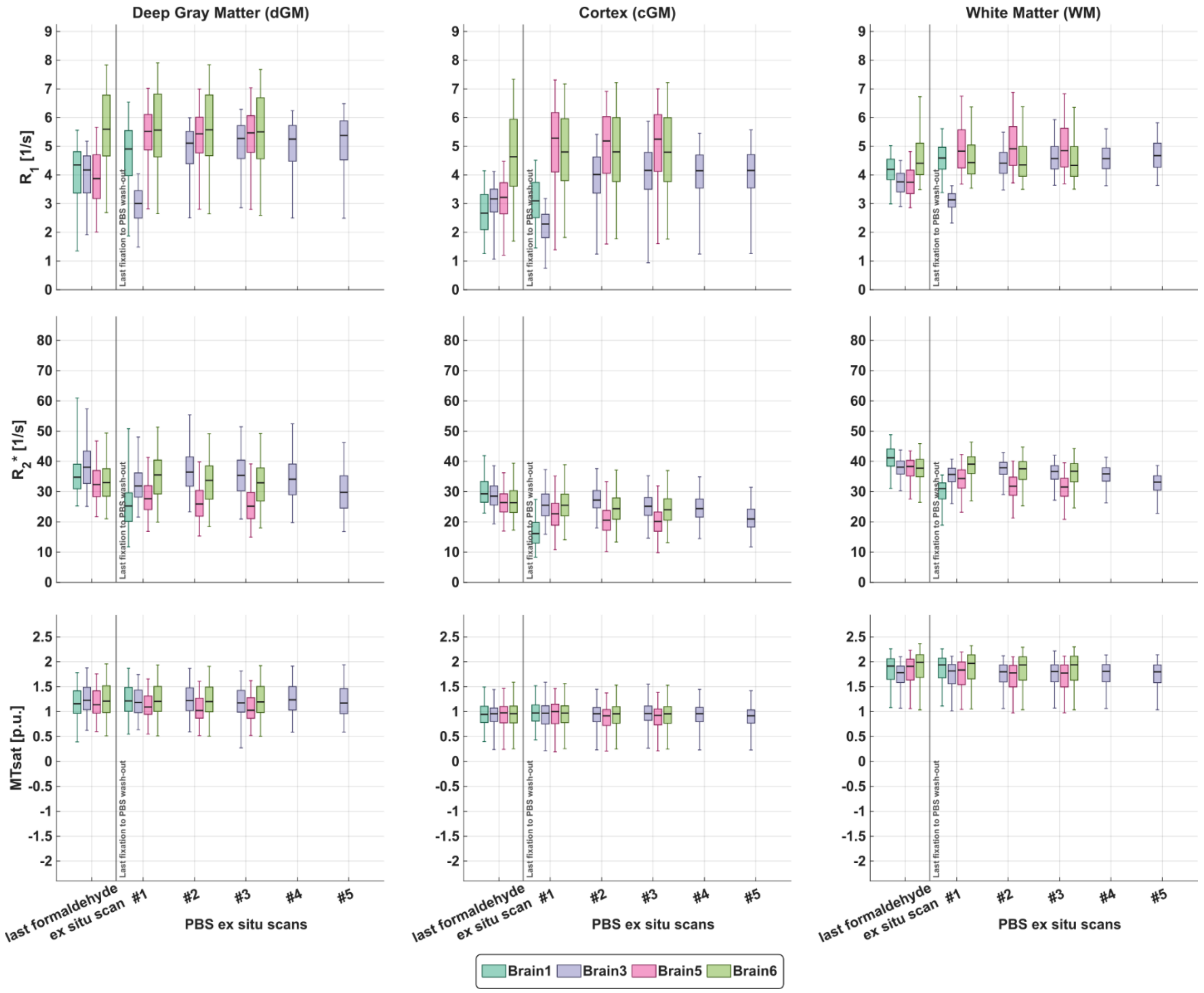


*Figure S5: Median and interquartile (25% and 75%) in the change of the MPM parameters ($R_1$, $R_2$* and $MT_{sat}$) between last formaldehyde ex situ and PBS ex situ scans per specimen (color). The PBS ex situ scan sessions started, at least, one week after changing the formaldehyde solution to PBS. A dashed x-line was added to separate the last formaldehyde ex situ scan time point and the PBS ex situ scan sessions. Note that the mean and standard deviation shown in Figure 3 (Results §3.1) represents the latest PBS ex situ scan session across all specimens (PBS ex situ scan #1 for Brain1, #3 for Brain5 and Brain6, and #5 for Brain3).*

To complement further the change between the last formaldehyde and PBS *ex situ* scan sessions across all MPM parameters, we present in Figure S6 the change of the fluid mask in both measurements. As observed, the trend that follows the fluid mask at the last formaldehyde and PBS is identical to the tissue classes in Figure S5 for all the contrasts except $N_A$: barely changed for $MT_{sat}$, variable change for $R_1$, variable decrease for A and high decrease for $R_2$*.

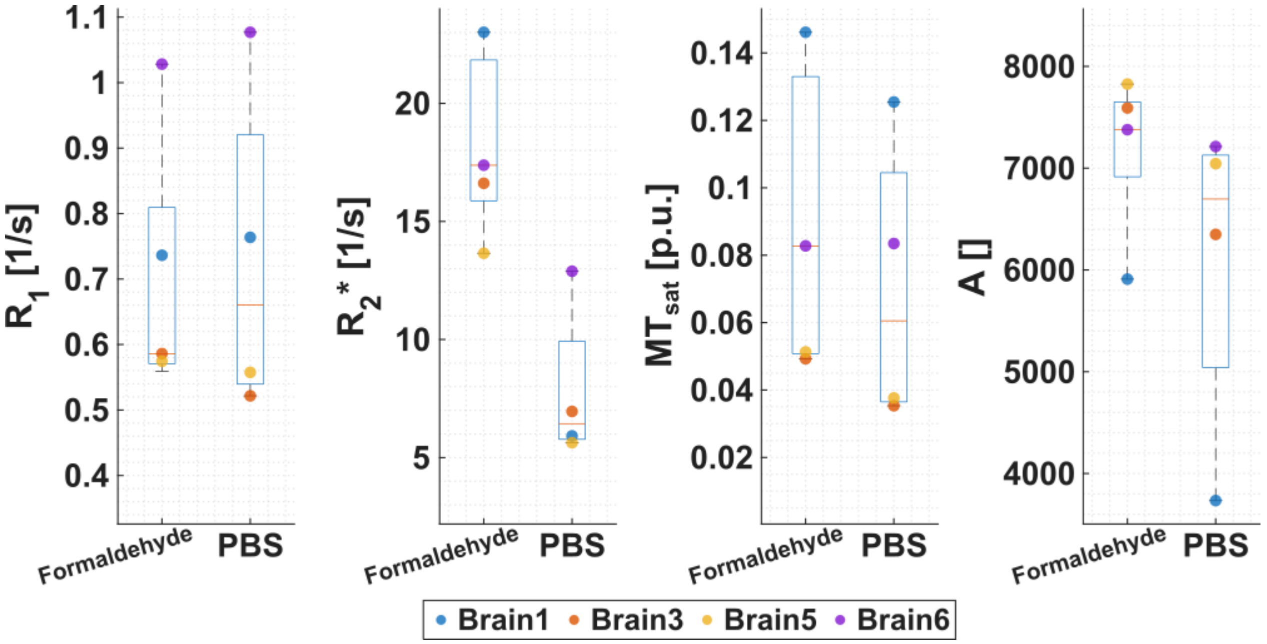


*Figure S6: Group median and quartiles (box plots) of the fluid mask in the last formaldehyde and PBS ex situ scan sessions per specimen (colored markers) for each MPM parameter: $R_1$, $R_2$*, $MT_{sat}$, and A. The last formaldehyde ex situ scan session varied between 93 to 603 days, while the last PBS ex situ scan session varied between 32 and 200 days. Brain2 is excluded from this comparison given its shorter fixation (8 days) and not acquired PBS ex situ scan.*

### S3. Model selection and predictive power of the preferred model

This section describes how we compared candidate temporal models for (i) fixation-related changes in the MPM-parameters (§S3.1) and (ii) relative tissue-volume shrinkage during formaldehyde *ex situ* immersion (§S3.2). We evaluated these two cases in two sequential steps: (1) assessing “model preference” by fitting and ranking the candidate models to the data using the Akaike Information Criterion corrected (AICc, (Burnham et al., 2011; Ghosh et al., 2007)) and (2) by estimating out of sample performance, or predictive power, of the preferred model using leave-one-out cross-validation (LOOCV, e.g. in (Molinaro et al., 2005)). These evaluations were performed for each tissue class and MPM parameter/relative volume.

For selecting the preferred model, we fitted the candidate models to the data and then evaluated them using the AICc, defined by:

$$AICc_m = n \cdot \log\left(\frac{SSE\left(R_f^m, R_n\right)}{n}\right) + 2 \cdot k + \frac{2\left(k \cdot (k+1)\right)}{n-k-1} \quad \text{(S2)}$$

where n is the number of time samples, k is the number of parameters per model m and $SSE\left(R_f^m, R_n\right)$ is the sum of squares difference between the fitted model and time samples $R_n$. The preferred model is the one with the lowest AICc, i.e., $AICc_{min}$.

Relative support for each competing model was then summarized by the evidence ratio (ER; (Burnham et al., 2011; Burnham and Anderson, 2004, 2002)), defined as:

$$ER(AICc_{min}, AICc_m) = \exp\left(0.5(AICc_m - AICc_{min})\right) \quad \text{(S3)}$$

where the subscripts in $AICc_m$ and $AICc_{min}$, i.e. m and min, indicate each model used; and the minimum AICc across the models, respectively.

By construction, the preferred model has always ER = 1. Larger ER indicates progressively weaker support relative to the preferred model. Table S1 summarizes the interpretative bands used in this work (adapted from (Burnham and Anderson, 2004)):

| $AICc_m - AICc_{min}$ | $ER(AICc_{min}, AICc_m)$ | Interpretation |
|---|---|---|
| $\leq 2$ | $\leq 2.71$ | The model "m" has substantial support for being as preferable as the preferred model |
| $2 \leq x \leq 10$ | $2.71 \leq x \leq 148.41$ | The model "m" has considerably less support in being preferable as the preferred model |
| $> 10$ | $> 148.41$ | The model "m" has essentially no support for being preferable as the preferred model |

*Table S1: Summary of the interpretation of the evidence ratio (ER) based on the difference between the corrected Akaike Information Criterion per model and the minimum across models.*

Then, after selecting the preferred model, we assessed how well that model generalized across specimens using leave-one-out cross-validation. This method consists of fitting the preferred model N times, where N is the number of observations (or datasets in our case), in which each time one observation is left out from the training set. The left-out observation was then used as the predicted dataset by the corresponding model. From this cross-validation, two parameters can be estimated: the Kendall's τ coefficient and the root-mean-squared error (RMSE).

Kendall's τ quantifies rank agreement between observed and predicted values, ranging from −1 indicating complete disagreement to +1 indicating perfect agreement. RMSE quantifies absolute prediction error. For that, the RMSE is computed from the LOOCV predictions as $RMSE = \sqrt{\frac{1}{N}\sum_{i=1}^{N}(y_i - \hat{y})^2}$, where $y_i$ denotes the measured value and $\hat{y}$ the corresponding LOOCV prediction. With only five specimens, LOOCV provides a transparent specimen-wise check of generality rather than definitive unbiased population estimates.

Subsections §S3.1 and §S3.2 apply this procedure to fixation modeling of the MPM parameters and to shrinkage modeling of relative volume, respectively.

#### S3.1 Fixative modeling

For the fixative modeling evaluation, we used four models: reciprocal logistic Yong-Hing (Equation 1), mono-exponential saturation (Equation 2), bi-exponential saturation (Equation 3) and the null (Equation 4) models. Table S2 summarizes the AICc values of each model fitted to each MPM parameter, i.e. $R_1$, $R_2$*, $MT_{sat}$ and $N_A$ and region of interest (ROI), during the fixation process across specimens (Results §3.2, Figure 4). From these values, ER is estimated (Equation S3) and illustrated in Figure S7.

Across almost all the MPM parameters and tissue classes, either the mono- or bi-exponential models were the clearly preferable model (indicated by a black arrow) in comparison to the YH model and the null model. For $N_A$, the null model was the uniquely preferable model for almost all tissue classes, except WM. This was expected given the lack of temporal dependency of this MPM parameter as shown in Figure 4 (Results §3.2). For WM-NA, on the other hand, this was affected by the specimen's inter-variability.

| MPM-Contrast | ROI | Inverse Yong-Hing | Mono-exponential | Bi-exponential | Null |
|---|---|---|---|---|---|

| | | | | | |
|---|---|---|---|---|---|
| $R_1$ | dGM | -504.67 | -531.86 | -529.53 | -38.365 |
| | cGM | -733.94 | -782.36 | -784.47 | -151.09 |
| | WM | -591.63 | -622.57 | -620.85 | -72.323 |
| $R_2$* | dGM | 254.63 | 254.64 | 257.53 | 331.46 |
| | cGM | 72.671 | 71.91 | 63.438 | 172.15 |
| | WM | 113.03 | 108.73 | 59.336 | 319.5 |
| $MT_{sat}$ | dGM | -1034.2 | -1034.2 | -1029.8 | -1013.6 |
| | cGM | -1388.2 | -1388.2 | -1383.8 | -1337.4 |
| | WM | -810.89 | -810.91 | -855.27 | -805.06 |
| $N_A$ | dGM | -1551.9 | -1552 | -1547.6 | -1556.2 |
| | cGM | -1457.2 | -1457.3 | -1453.1 | -1457 |
| | WM | -1459.5 | -1459.5 | -1493.7 | -1463.7 |

*Table S2: Summary of the estimated corrected Akaike Information Criterion (AICc, Equation S2) of each fitted model to the temporal change across all specimens of each MPM parameter during fixation per tissue class (ROI).*

In several cases (e.g., dGM-R2*, cGM- and dGM-$MT_{sat}$; and cGM-$N_A$), two (or three) models yielded very similar AICc values (Table S2), indicating weak preference by AICc alone. In these instances, we resolved model choice by considering (i) parameter uncertainty and (ii) parsimony (selecting the simpler functional form when fits were comparably supported) (Aho et al., 2017; Tredennick et al., 2021; Yates et al., 2023).

*Figure S7: Comparison of evidence ratios (ER) for the null, mono- and bi-exponential saturation, and the inverse Yong–Hing models; for characterizing fixation-related changes in MPM parameters up to 150 days. ER was computed from the corrected Akaike Information Criterion (AICc). The preferred model (ER = 1) is marked with a black arrow. Models with ER < 2.71 (lower dashed line) are considered similarly plausible as the preferred model, whereas models with ER > 148.41 (upper dashed line) have essentially no support. Results are shown for each tissue class (dGM, cGM, WM) and each MPM parameter. Triangular shape of the box plots indicates cases where ER diverged (ER → ∞).”*

Table S3 lists the fitted parameters of the preferred model for each MPM parameter and tissue class (Results §3.2, Figure 4). $R_{ev,0}$ is fitted intercept at fixation day 0, $\Delta R_{s1}$ (or $\Delta R_s$ in the case of the mono-exponential model, Equation 2) and $\Delta R_{s2}$ are the short and long saturation ratio, and $t_{s1}$ (or $t_s$ in the case of the mono-exponential model, Equation 2) and $t_{s2}$ are the short and long saturation times in days. Units depend on the MPM parameter and are specified in the table. For dGM $R_2$*, cGM and dGM $MT_{sat}$, and cGM $N_A$, we also report parameters from alternative models with similar AICc support.

| MPM-Contrast | ROI | Model | $R_{ev,0}$ (sd) | $\Delta R_{s1}$ (sd) | $t_{s1}$ (sd) | $\Delta R_{s2}$ (sd) | $t_{s2}$ (sd) |
|---|---|---|---|---|---|---|---|

| | | | | | | | |
|---|---|---|---|---|---|---|---|
| $R_1$ | dGM | Mono-exponential | 1.00133 (0.04478) | 2.9534 (0.063566923) | 39.2321 (2.48453474) | - | - |
| | cGM | Bi-exponential | 0.725941 (0.085162) | 0.188237 (0.083457509) | 1.402167 (1.16127952) | 2.4291 (0.060191942) | 62.7913 (4.27015349) |
| | WM | Mono-exponential | 1.09701 (0.033705) | 2.7463 (0.053686737) | 42.9235 (2.41012014) | - | - |
| $R_2^*$ | dGM | Inverse YH | 0.034373 (0.0006943) | 0.0054463 (0.00067616) | 12.518 (2.7819) | - | - |
| | | →Mono-exponential | 29.0029 (0.61682) | 5.5821 (0.61053661) | 13.6629 (3.09896706) | - | - |
| | cGM | Bi-exponential | 20.1246 (1.0368) | 3.6923 (1.0155762) | 1.2837 (0.726953544) | 2.90645 (0.42832741) | 25.87375 (9.58415813) |
| | WM | Bi-exponential | 26.9675 (0.60303) | 6.4759 (0.63854766) | 4.38075 (0.888601171) | 12.8841 (18.517021) | 293.6695 (538.75132) |
| $MT_{sat}$ | dGM | Inverse YH | 0.95679 (0.028118) | 0.10131 (0.027932) | 2.3865 (0.89505) | - | - |
| | | →Mono-exponential | 1.04297 (0.031801) | 0.12597 (0.031595779) | 2.47886 (0.927273068) | - | - |
| | cGM | Inverse YH | 1.1061 0.010364 | 0.061469 0.010211 | 4.8152 1.2185 | - | - |
| | | →Mono-exponential | 0.903644 (0.0086142) | 0.053645 (0.0084915177) | 4.92903 (1.24521095) | | |
| | WM | Bi-exponential | 1.70061 (0.045797) | 3.6657 (271.76057) | 6.9577 (33.1559232) | -3.55228 (271.77649) | 7.896567 (37.8444543) |

| | | | | | | | |
|---|---|---|---|---|---|---|---|
| $N_A$ | dGM | Null | 0.892604 (0.00073087) | - | - | - | - |
| | cGM | →Null | 0.918835 (0.00098352) | - | - | - | - |
| | | Inverse YH | 1.1075 (0.012422) | 0.019766 (0.012371) | 1.4356 (1.4913) | | |
| | | Mono-exponential | 0.90357 (0.0096589) | 0.015793 (0.0096189) | 1.6086 (1.6295) | | |
| | WM | Bi-exponential | 0.77721 (0.0031481) | 1.2305 (9961.7) | 98.164 (30277) | -1.3095 (9960.3) | 106.4 (34605) |

*Table S3: Parameters of the preferred models (see Figures 4 and S7, and Equations 1 to 4) in deep gray matter (dGM), cortical gray matter (cGM) and white matter (WM). Those models that have almost-to-equal AICc values (Table S2) per tissue class-MPM parameter are also reported (green boxes). The preferred model is indicated by an arrow (→). Standard deviation of each fitted parameter is given in parenthesis.*

A detailed view, i.e., individual y-scaling, of the best saturation model fitting the data per tissue class and MPM parameter is observed in Figure S8.

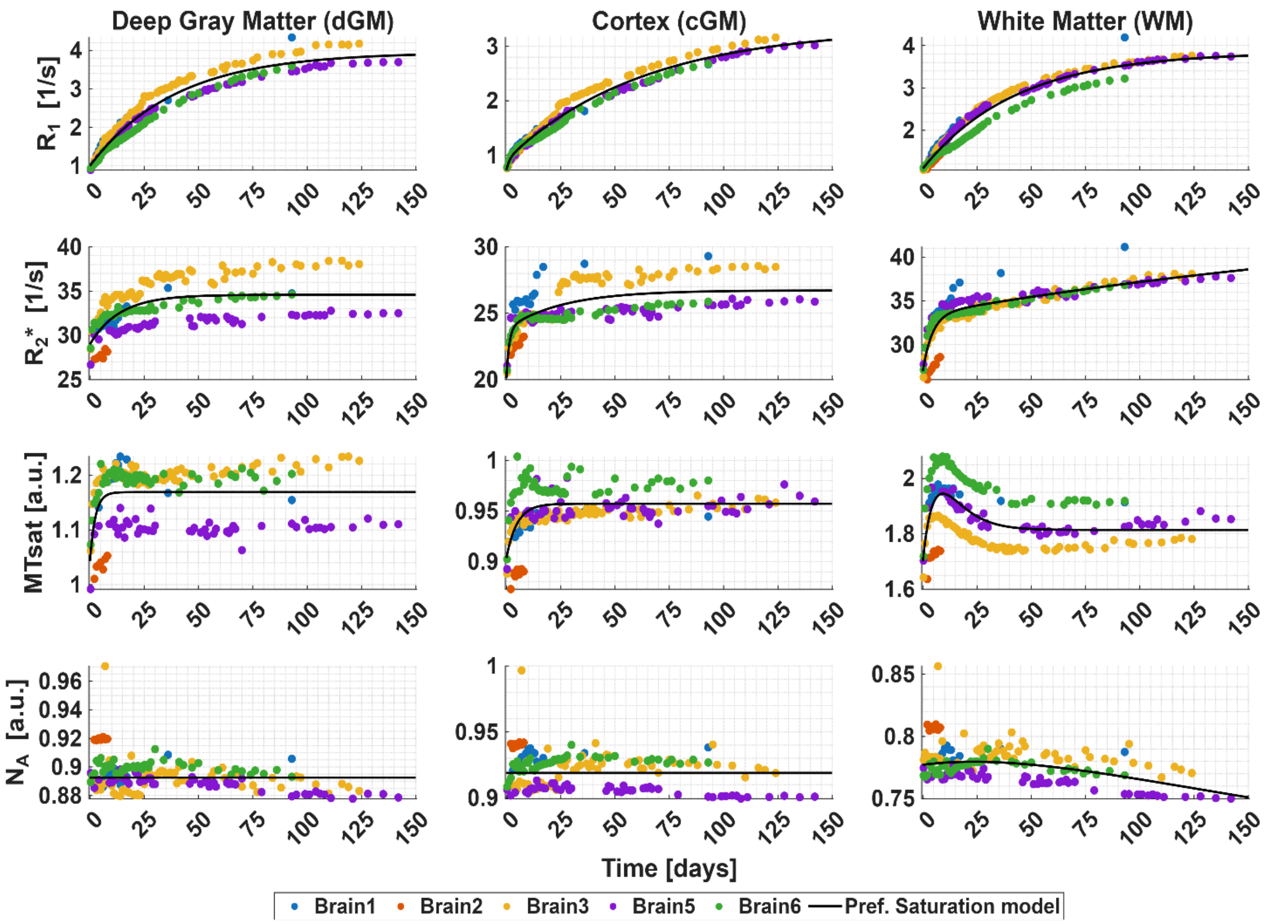


*Figure S8: Temporal evolution of the MPM parameters during fixation. For each tissue class (columns, left to right: dGM, cGM and WM), the median value of the MPM parameters is plotted (rows, top to bottom: $R_1$, $R_2$*, $MT_{sat}$ and $N_A$) for each brain specimen (colored markers) and per time point. The curve of the preferred saturation model ($\langle R_{ev}\rangle(t)$) is plotted for each tissue class and MPM parameter (black solid line) and its name shown on top of each subplot.*

Table S4 summarizes the harmonized fit parameters of the mono- and bi-exponential saturation models between tissue classes and MPM parameters. The definitions of each harmonized parameter are given in the following table:

| Harmonized parameters | Mono-exponential model | Bi-exponential model |
|---|---|---|
| Total saturation amplitude A | $\Delta R_s$ | $\Delta R_{s1} + \Delta R_{s2}$ |
| Total saturation S | $R_{ev,0} + \Delta R_s$ | $R_{ev,0} + A$ |
| Effective saturation rate $c_{eff}$ | $1/t_s$ | $\frac{\left(\frac{\Delta R_{s1}}{t_{s1}} + \frac{\Delta R_{s2}}{t_{s2}}\right)}{A}$ |
| Effective saturation time $t_{eff}$ | $t_s$ | $1/c_{eff}$ |

*Table S4: Definition of the harmonized parameters as a function of the fitted parameters from the mono- and bi-exponential models. Standard deviation of each harmonized fitted parameter is estimated via error propagation.*

Standard deviation of these harmonized parameters was also estimated via first-order error propagation, by assuming that the fitted parameters are uncorrelated and exhibit Gaussian-like behavior.

| MPM contrast | ROI | A (sd) | S (sd) | $C_{eff}$ (sd) [1/days] | $T_{eff}$ (sd) [days] |
|---|---|---|---|---|---|
| $R_1$ [1/s] | dGM | 2.9534 (0.0636) | 3.9547 (0.0778) | 0.0255 (0.0633) | 39.2321 (2.4845) |
| | cGM | 2.6173 (0.1029) | 3.3432 (0.1336) | 0.0661 (0.0434) | 15.1349 (0.6564) |
| | WM | 2.7463 (0.0537) | 3.8433 (0.0634) | 0.0233 (0.0561) | 42.9235 (2.4101) |
| $R_2$* [1/s] | dGM | 5.5821 (0.6105) | 34.5850 (0.8679) | 0.0732 (0.2268) | 13.6629 (3.0990) |
| | cGM | 6.5988 (1.1022) | 26.7234 (1.5132) | 0.4529 (0.2476) | 2.2080 (0.5467) |
| | WM | 19.3599 (18.5280) | 46.3275 (18.5378) | 0.0786 (0.0160) | 12.7190 (0.2040) |
| $MT_{sat}$ [p.u.] | dGM | 0.1260 (0.0316) | 1.1689 (0.0448) | 0.4034 (0.3741) | 2.4789 (0.9273) |
| | cGM | 0.0536 (0.0085) | 0.9573 (0.0121) | 0.2029 (0.2526) | 4.9290 (1.2452) |
| | WM | 0.1134 (384.339) | 1.8141 (384.339) | 0.6788 (404.54) | 1.4731 (595.94) |

*Table S5: Harmonized parameters of the mono- and bi-exponential preferred models (see Table S3) in deep gray matter (dGM), cortical gray matter (cGM) and white matter (WM) for $R_1$, $R_2$* and $MT_{sat}$. Standard deviation of each harmonized fitted parameter is given in parenthesis. A: total saturation amplitude, S: total saturation, Ceff: effective saturation rate and teff: effective saturation time. A and S have the units of the respective MPM parameters.*

Left column in Figure S9 shows the Kendall's $\tau$, while the right column shows the RMSE. While the Kendall's $\tau$ was already reported in the main manuscript, we will report the RMSE results here. The median RMSE (interquartile range: (Q1 Q3), with Q1 the 25% and Q3 the 75% percentile) was between 0.1122 (0.0741 0.1572) $s^{-1}$ in cGM and 0.2921 (0.1996 0.3194) $s^{-1}$ in dGM for $R_1$, between 0.9935 (0.8090 2.9571) $s^{-1}$ in WM and 3.4525 (0.8225 3.7725) $s^{-1}$ in dGM for $R_2$*, between 0.0132 (0.0118 0.0412) p.u. in cGM and 0.1203 (0.0456 0.1715) p.u. in WM for $MT_{sat}$, and between 0.0085 (0.0048 0.0106) in dGM and 0.0140 (0.0087 0.0156) in cGM for $N_A$.

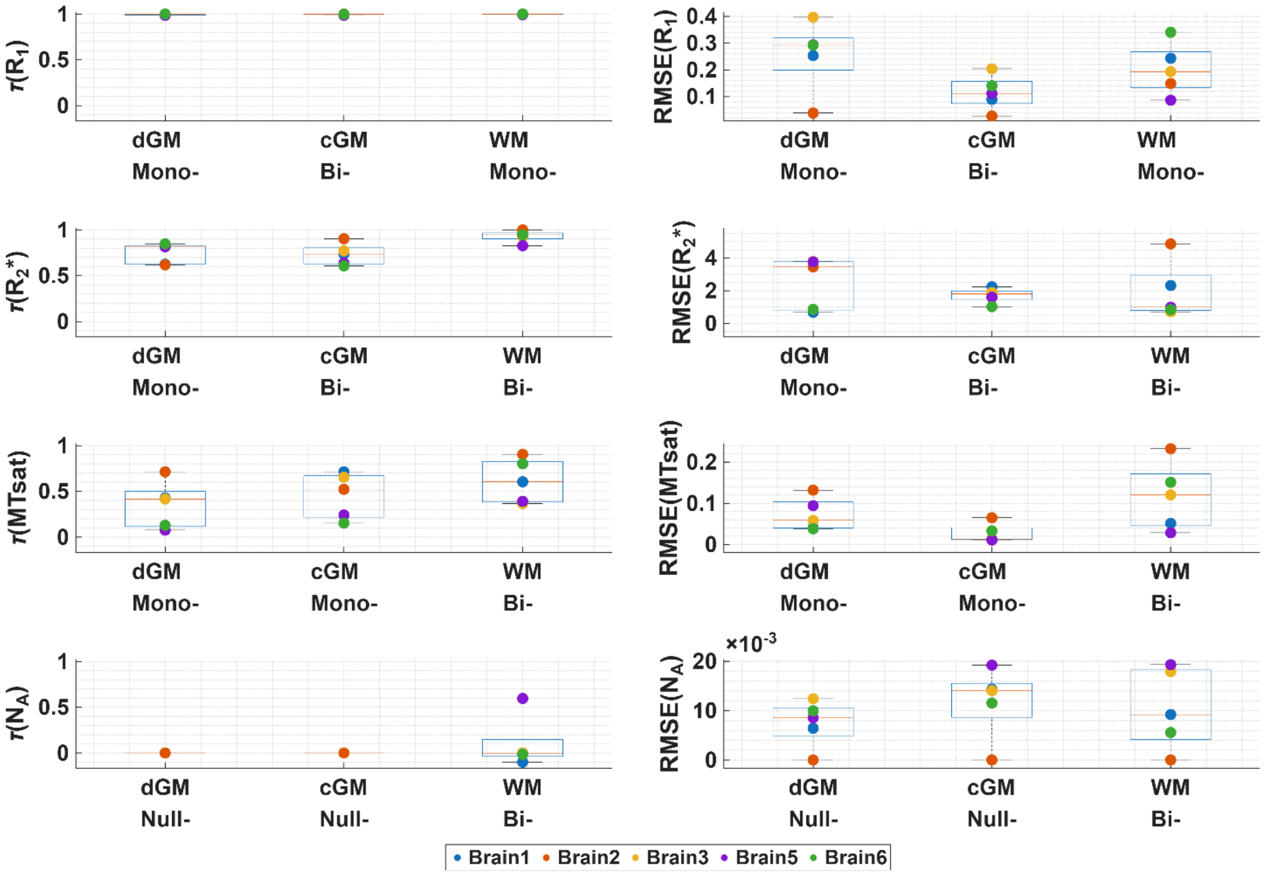


*Figure S9: Quantitative assessment of how well the preferred model characterizes the temporal fixation using "leave-one-out" cross validation. Depicted are the Kendall's correlation coefficient ($\tau$,first column) and the root-mean-square error (RMSE, second column) for each MPM parameter (rows, top to bottom: $R_1$, $R_2$*, $MT_{sat}$ and $N_A$). Each plot contains the estimated metric for each brain specimen (colored markers) per tissue class (deep gray matter, dGM; cortical gray matter, cGM; and white matter, WM) and preferred model (Mono-exponential model, mono-; bi-exponential model, bi- and null model, null-). The median and quartiles are shown across brain specimens (box plots).*

#### S3.2 Shrinkage modeling

For the tissue shrinkage modeling evaluation, we used two models: the null model (Equation 4) and the heuristic model (Equation 6).

Table S6 compiles the AICc values of each model fitted to each region of interest (ROI), during the tissue shrinkage process across specimens (Results §3.3, Figure 6). From these values, ER is estimated (Equation S3) and illustrated in Figure S10.

| ROI | Null | Heuristic |
|---|---|---|
| dGM | -1338.6 | -1398.02 |
| cGM | -1501.8 | -1687.13 |
| WM | -1411.4 | -1497.19 |

*Table S6: Summary of the estimated corrected Akaike Information Criterion (AICc, Equation S2) of each fitted model to the temporal tissue shrinkage across all specimens per tissue class (ROI).*

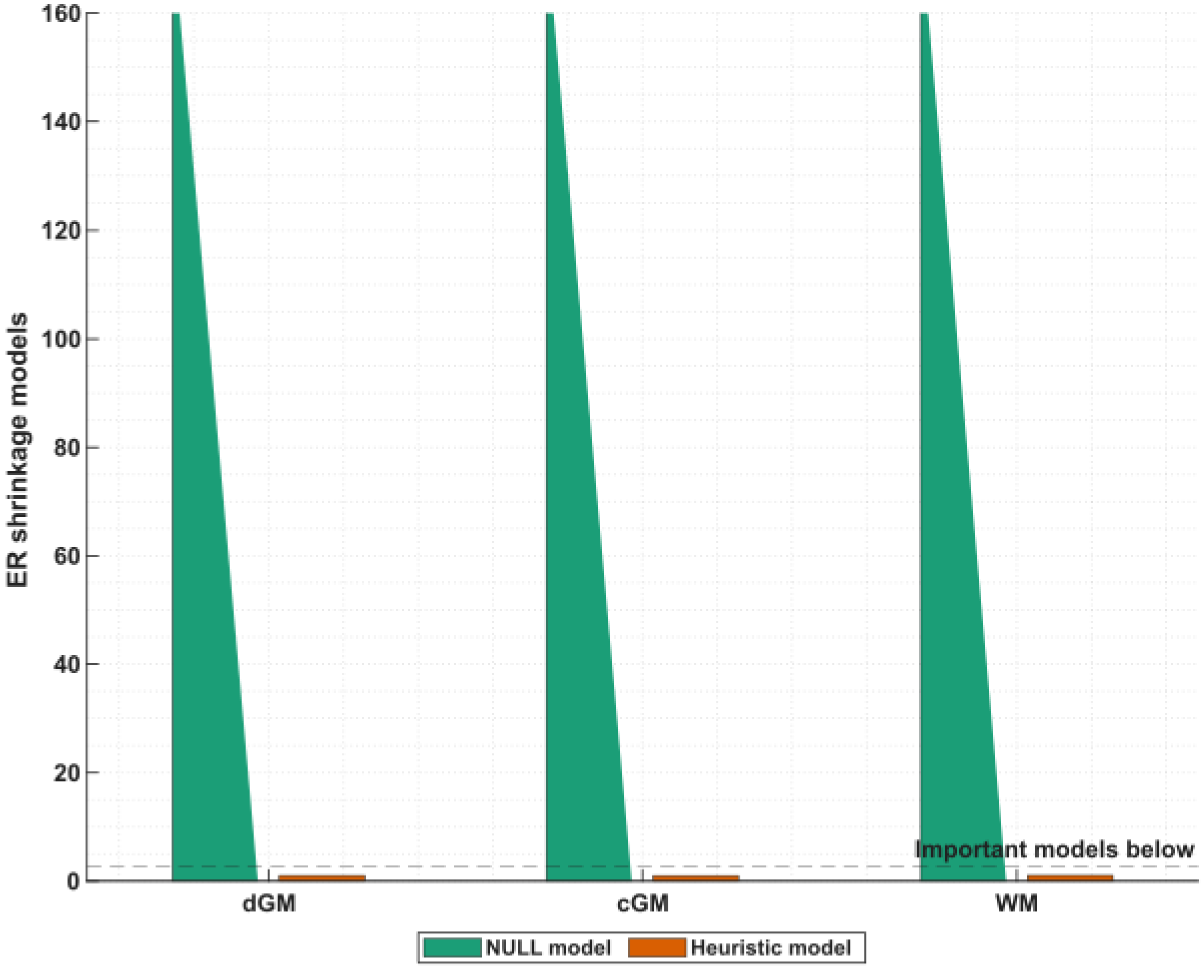


*Figure S10: ER comparison for the null model and heuristic shrinkage model up to 150 days. ER was computed from AICc. The heuristic model is preferred across tissue classes (ER < 2.71), whereas the null model shows essentially no support (ER > 148.41). Triangular markers indicate ER → ∞.*

Table S7 compiles the parameters of the preferred model (from Figure S10) that best described the temporal tissue shrinkage during the fixation process across specimens (Results §3.3, Figure 6) per tissue class. Each parameter is defined as: δ is the decrement shrinkage parameter (no units), and $t_s$ is the saturation time (days).

| ROI | Model | δ ± sd. | $t_s$ ± sd. |
|---|---|---|---|
| dGM | Heuristic | 0.05149 ± 0.00398 | 41.118 ± 6.7688 |
| cGM | Heuristic | 0.05178 ± 0.007996 | 85.4 ± 20.647 |
| WM | Heuristic | 0.04031 ± 0.00193 | 27.976 ± 3.3745 |

*Table S7: Parameters of the preferred models (see Figures 5 and S11, and Equations 6 and 7) in deep gray matter (dGM), cortical gray matter (cGM) and white matter (WM). Standard deviation of each fitted parameter is given after the ±.*

Left column in Figure S11 shows the Kendall's $\tau$, while the right column shows the RMSE. While the Kendall's $\tau$ was also reported in the main manuscript, we will report the RMSE results here. The median RMSE (interquartile range: (Q1 Q3), with Q1 the 25% and Q3 the 75% percentile) was 0.006478 (0.002696 0.01560) for cGM, 0.009017 (0.005988 0.01371) for WM and

0.014722 (0.008259 0.02035) for dGM while the magnitude of the interquartile ranges was 0.0129, 0.0077 and 0.0121, respectively.

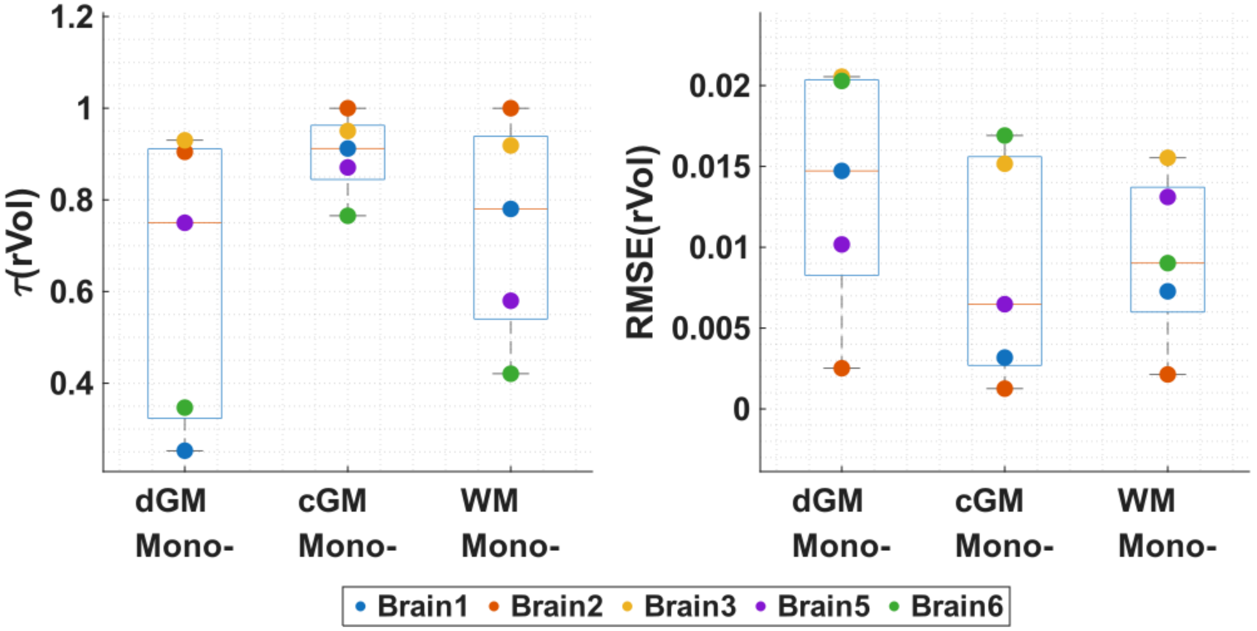


*Figure S11: Quantitative assessment of how well the preferred model characterizes the temporal tissue shrinkage using "leave-one-out" cross validation. Depicted are the Kendall's correlation coefficient (τ,first column) and the root-mean-square error (RMSE, second column). Each plot contains the estimated metric for each brain specimen (colored markers) per tissue class (deep gray matter, dGM; cortical gray matter, cGM; and white matter, WM) and preferred model (Mono-: heuristic model). The median and quartiles are shown across brain specimens (box plots).*

### S4. Assessment of potential technical confounds in MPM parameter estimation

Several technical factors could affect the estimated MPM parameters independently of biological and chemical tissue changes. These factors include use of an acquisition protocol optimized for *in vivo* imaging, scanner-platform differences between the Trio and Prisma systems, temperature variation during scanning, the age difference between the *in vivo* and *ex vivo* cohorts, and unequal sampling across tissue conditions. We therefore performed complementary sensitivity analyses to estimate the magnitude of these potential biases. First, we evaluated whether using the *in vivo*-optimized MPM protocol could bias parameter estimation under the range of fixation-related $R_1$, PD, and MTsat values observed in this study. Second, we performed a same-subject Siemens Trio versus Prisma comparison using matched MPM sequence parameters to assess scanner-platform effects. Third, we assessed intra-measurement temperature variation using ADC-derived estimates. Fourth, we bounded cross-state temperature effects and aging using published relaxometry relationships. Last, we present similar results from analyses 1 (Methods §2.3) and 2 (Methods §2.4), but by only considering the three completely acquired datasets, i.e. with *in situ* and PBS *ex situ* scans.

#### S4.1 Potential bias from the *in vivo*-optimized MPM protocol

Given the observed increase in $R_1$, PD and $MT_{sat}$, there is uncertainty regarding whether these quantitative and qualitative behaviors could be driven by the choice of MPM protocol. This uncertainty is because the MPM protocol sequence parameters such as repetition time (TR) and flip angle (FA) must be optimized with respect to the Ernst angle of the object to be measured, in this case our brain specimens[2]. To evaluate any potential issues, we used our fitted preferred models, Table S3, to generate "real" MPM parameter values and assessed both the normalized bias, defined as the ratio between the estimated MPM and “real” MPM parameters minus 1; and the variation, defined by the coefficient of variation, which we defined as the ratio between the standard deviation and the mean estimated MPM parameter per time point after adding simulated random noise. This noise was estimated by dividing the signal of the first echo of the simulated data by the signal-to-noise ratio calculated from the *in situ* time point of Brain1. For PDw signal, the SNR was 32.22 and for $T_1$w was 27.40.

The comparison between noisy and noise-free datasets revealed significant insights into the bias and variance introduced by the MPM protocol, as shown in Figure S12. Overall, $MT_{sat}$ showed the highest bias, followed by $R_1$ and PD; with a normalized bias less than 1.6% across all the studied MPM parameters and time points. However, the coefficient of variation was also higher for MTsat, up to 23%, stabilizing around 8.5% for $R_1$ and decreasing down to 3% for PD.

This analysis suggests that using the same *in vivo optimized* MPM protocol introduced only a small bias under the simulated conditions; therefore, no additional bias correction was applied. However, the increased variance suggests that higher SNR would be beneficial for improving parameter precision.

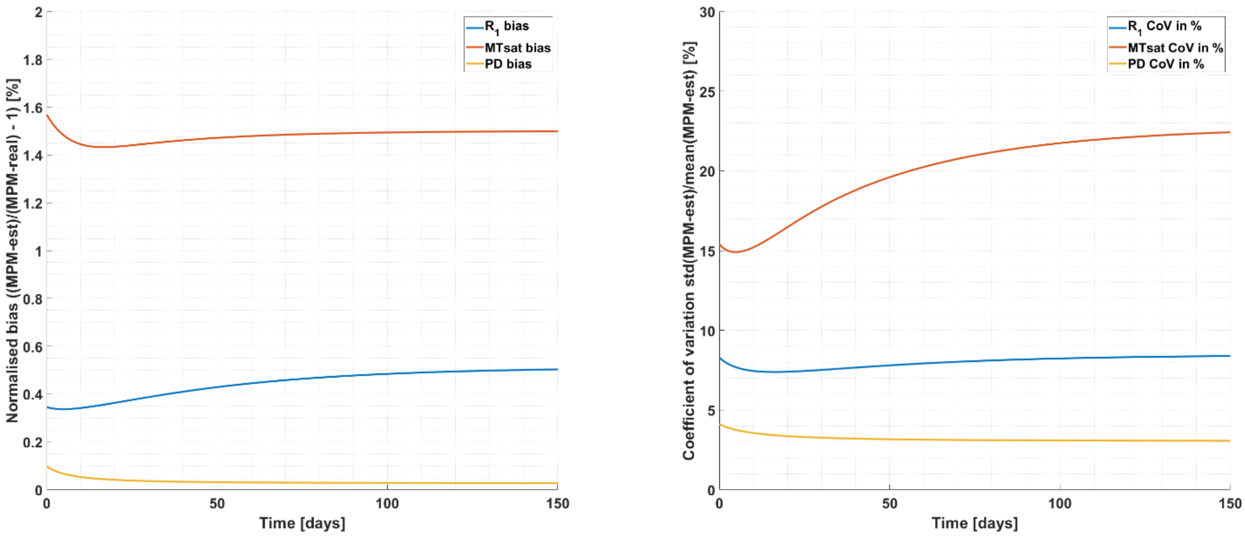


*Figure S12: Temporal bias and variation introduced by the sub-optimized MPM protocol given the temporal change of the MPM parameters, i.e. $R_1$, PD and $MT_{sat}$, due to fixation in white matter. For each MPM parameter (blue for $R_1$, red for $MT_{sat}$ and yellow for PD), a normalized bias (left plot) and the coefficient of variation (right plot) were calculated between the estimated MPM parameter given the in vivo MPM protocol and the “real” MPM parameter value, defined by the preferred model for each MPM parameter in white matter (Results §3.2, Figure 4; Figure S7 and Table S3).*

#### S4.2 Scanner-platform comparison between Siemens Trio (*in vivo* cohort) and Prisma (*in situ* and *ex situ* experiments)

Because the qualitative *in vivo* reference dataset was acquired on a Siemens Trio system whereas the *in situ* and *ex situ* experiments were acquired on a Siemens Prisma system, we

[2] $R_2$* does not have this problem, since it depends strongly on the echo time (TE) range rather than TR and FA parameters.

assessed whether scanner-platform differences could represent a relevant source of bias in the qualitative comparison. For this purpose, we performed a same-subject MPM comparison between a Trio acquisition obtained on 2016-01-19 and a Prisma acquisition obtained on 2018-02-26 using matched MPM sequence parameters. Both datasets were processed with the same hMRI pipeline, brain-masked, registered to a common pairwise mean-template space, and summarized within cortical gray matter, deep gray matter, and white matter masks derived from the mean registered MTsat image. Scanner-related differences were summarized descriptively using the symmetric median percent difference, defined as:

$$\Delta med(\%) = \frac{median(Prisma)-median(Trio)}{mean(medianPrisma, medianTrio)} \cdot 100\%, \qquad \text{(S4)}$$

for each MPM contrast and tissue class.

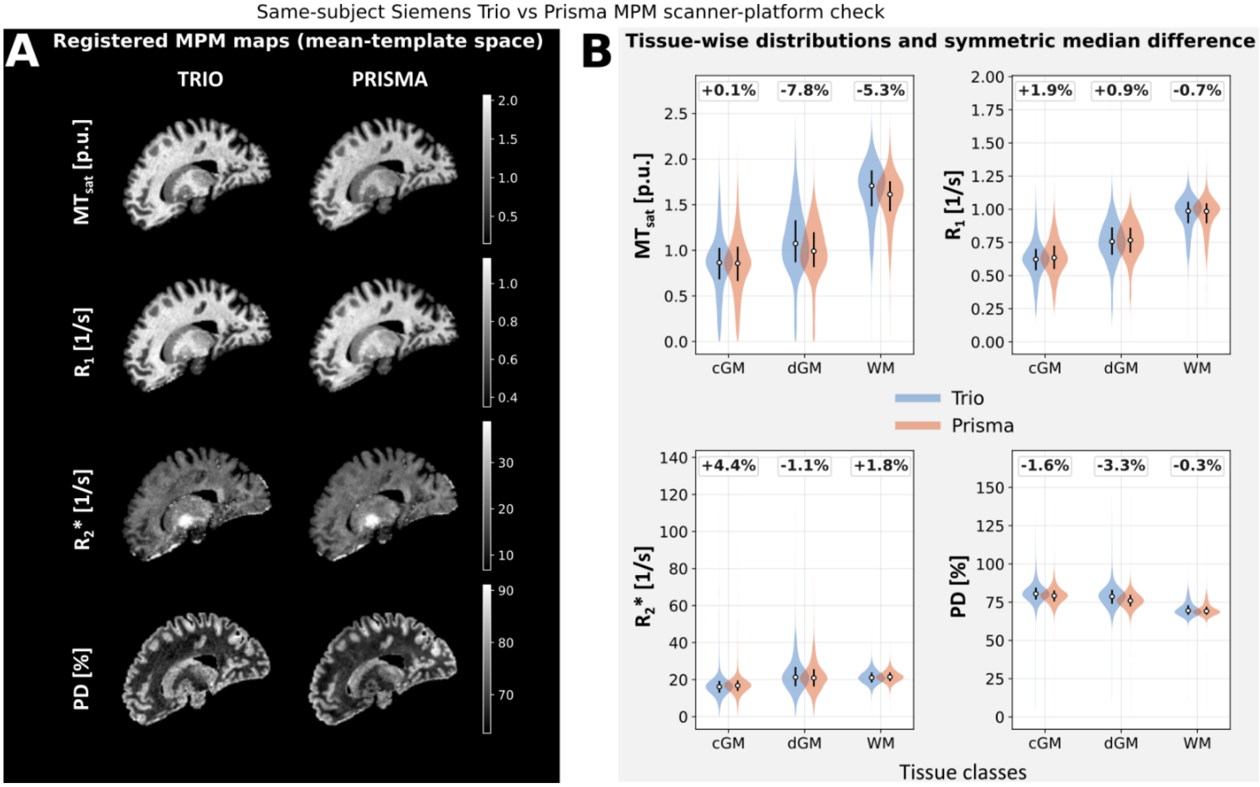


*Figure S13: Assessment of scanner-platform differences between Siemens Trio and Prisma MPM acquisitions. (A) Representative registered MPM maps from the same participant acquired on the Siemens Trio and Prisma systems, shown in common mean-template space for $MT_{sat}$, $R_1$, $R_2$*, and PD. (B) Tissue-wise distributions of MPM values in cortical gray matter (cGM), deep gray matter (dGM), and white matter (WM). Bold labels indicate the symmetric median percent difference, calculated as 100 × (medianPrisma − medianTrio) / mean(medianPrisma, medianTrio) (Equation S4).*

The resulting maps and tissue-wise distributions are shown in Figure S13. Spatial patterns were qualitatively similar across scanners, and tissue-wise median differences were generally small for $R_1$, PD, and $R_2$*, whereas the largest deviations were observed for $MT_{sat}$ in deep gray matter and white matter. This overall magnitude is in line with previous MPM reproducibility work showing low inter-site bias and modest inter-site variation for $R_1$, PD*, and $MT_{sat}$, with somewhat higher variability for $R_2$* in multi-site settings (Thomas et al., 2024; Weiskopf et al., 2013). Taken together, these results suggest that scanner-platform differences are unlikely to account for the substantially larger tissue- and fixation-related MPM changes described in the main manuscript and support the use of the *in vivo* cohort as a qualitative contextual reference rather than a quantitative comparator.

#### S4.3 Intra-measurement temperature variation assessed from apparent diffusion coefficient measurements

Because sample temperature was not monitored directly during MPM sessions, we used diffusion-derived mean diffusivity (MD/ADC) measurements that were interleaved with our routine *ex vivo* acquisition protocol. ADC was acquired in most longitudinal scan sessions across specimens; here we report one representative session in detail (Brain6, formaldehyde *ex situ*, day 1).

ADC maps were analyzed at repeated acquisition points distributed across the session (blue lines in Figure S14 top). Values were extracted within a labeled mask covering the complete container, including all voxels with labels greater than zero, and voxels with non-finite or non-positive MD values were excluded to avoid undefined logarithmic ratios. We evaluated two reference frames. First, we used the first analyzed ADC measurement as the session-start reference, representing the closest available proxy to the initial room-temperature state. Second, we estimated a stepwise ADC proxy between consecutive ADC measurements from the first MPM acquisition to the one after the last MPM measurement (indicated by labels in Figure S14 top). For each ADC measurement, the voxelwise ADC proxy change was computed as:

$$\Delta ADC_{proxy}(k) = \left(\frac{\ln\left(\frac{MD_v(k)}{MD_v(0)}\right)}{\ln\left(MD_v(0)\right)}\right) \cdot 100\%, \tag{S5}$$

where $MD_v(k)$ and $MD_v(0)$ are the MD at measurement k and first measurement for voxel v, respectively. Equation S5 is a dimensionless logarithmic relative change expressed in percent and not a calibrated temperature difference ($\Delta T$). For the stepwise ADC proxy difference estimation, we replaced $MD_v(0)$ with $MD_v(k-1)$.

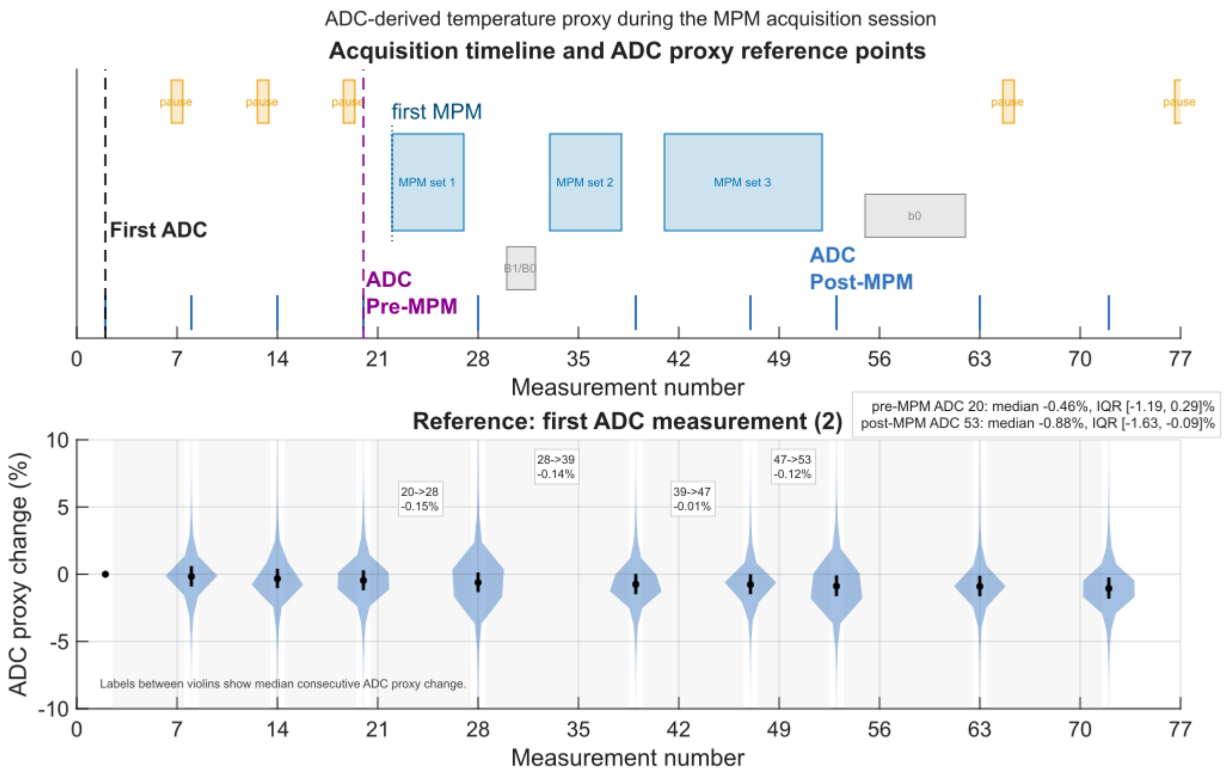


*Figure S14: ADC-derived temperature proxy during the MPM acquisition session. Repeated ADC measurements were used as a qualitative temperature-sensitive proxy during the scanning session. The upper panel shows the acquisition timeline, including ADC acquisitions (blue lines and color labels like First ADC and ADC pre- and post MPMs), pause periods, and MPM blocks. The lower panel shows voxelwise ADC proxy change relative to the first analysed ADC measurement, summarized as violin plots across all labelled voxels. Values between violins indicate median stepwise proxy change between consecutive ADC measurements to the first ADC measurement after the last MPM block.*

The ADC-derived proxy showed no substantial monotonic drift across the session (Figure S14 bottom). At the ADC measurement immediately before the first MPM acquisition, the median cumulative proxy change was -0.46% with an interquartile range of [-1.19, 0.29]%. At the first ADC measurement after the last MPM block, the median cumulative proxy change was -0.88% with an interquartile range of [-1.63, -0.09]%. We also computed stepwise changes between consecutive ADC measurements up to the post-MPM ADC measurement; the median stepwise changes were small across intervals (range: -0.17% to -0.01%). These results are consistent with the absence of large monotonic proxy drift during the MPM block in the representative session analyzed. ADC was also acquired in most other longitudinal sessions, but we did not repeat this analysis across all sessions because the check was exploratory and intended only to rule out large intra-session drift during MPM blocks. This analysis does not evaluate whether proxy changes would be the same in every future session, does not provide absolute temperature, and does not address temperature differences between *in vivo*, *in situ*, and *ex situ* conditions (main manuscript, Limitations). Because ADC in fixed tissue is not equivalent to free-water self-diffusion, this analysis should be interpreted only as qualitative temperature-related quality control and not as direct temperature monitoring or temperature correction.

#### S4.4 Theoretical bounds given the MPM parameter changes during the different tissue conditions: temperature and aging.

In this subsection, we illustrate predictions on how $R_1$ and $R_2$* MPM parameters will change from *in vivo* to *in situ* due to cohort's age difference and temperature. During fixation, these literature-based bounds address only temperature as a potential confounder; they do not imply that fixation-related MPM changes are caused solely by temperature. For age difference, we use the models defined in (Birkl et al., 2016, 2014; Callaghan et al., 2014). Note that we did not use any of the following calculations to correct any measured value in this work.

Callaghan et al., (2014) reported a linear relationship between age and $R_1$ and $R_2$*; expressed as $R_{old} = R_{young} + m_{slope}\Delta_{age}$, where $R_{old}$ and $R_{young}$ are either $R_1$ or $R_2$* for "old" and "young" groups, $m_{slope}$ is the linear slope that relates age and MPM parameter, and $\Delta_{age}$ is the age difference of both groups. Therefore, the % difference of the MPM parameters due to age difference will be estimated as $\Delta R_{age}[\%] = \frac{m_{slope}\Delta_{age}}{R_{young}} \cdot 100\%$. While for % difference for $R_1$ is only calculated for WM, % $R_2$* difference is calculated for cGM and dGM.

On the other hand, both studies of Birkl et al. reported a linear relationship between temperature and $T_1$ and $T_2$* of the form $T_{hot} = T_{cold} + m_{slope}\Delta_{temp}$, where $T_{hot}$ and $T_{cold}$ are either $T_1$ or $T_2$* for "hot" and "cold" temperatures, $m_{slope}$ is the linear slope that relates temperature and MPM-parameter, and $\Delta_{temp}$ is the temperature difference of both groups. In comparison to the % difference of MPM parameters due to age difference, the % difference due to temperature needs to be calculated first through $T_1$ and $T_2$* and then obtain the corresponding $R_1$ and $R_2$* and estimate the % difference.

1. From *in vivo* to *in situ*

The *in vivo* and *in situ* conditions differed in two relevant respects. First, tissue temperature decreased from approximately 37 °C *in vivo* to approximately 22 °C *in situ* (ΔT = −15 °C). Second, the mean age was 25.4 years in the *in vivo* cohort and 59.8 years in the *postmortem* cohort.

For $R_1$, we investigated the age-effect on only WM while for temperature we explored all tissue classes. Callaghan et al. reported significant age-related decreases mainly in WM, with slopes of 0.0007–0.0016 $s^{-1}$/year. For temperature, we used the corresponding unfixed-tissue $T_1$-temperature equations: 13.1 [ms/°C] T + 738.5 ms for dGM (based on basal ganglia), 17.4 [ms/°C] T + 962 ms for cGM and 3.4 [ms/°C] T + 695.8 ms for WM.

| Tissue | % $R_1$ in Fig 3 | % $R_1$ due to temp. | % $R_1$ due to age |
|---|---|---|---|
| dGM | 7.79% | 19.139% | No reliably reported |
| cGM | -3.57% | 19.408% | No reliably reported |
| WM | 3.65% | 6.618% | -2.4 to -5.5% |

*Table S8: Predicted %Δ$R_1$ (in vivo→in situ) from age (Callaghan) and cooling ~37→22–23 °C (Birkl); illustrative bounds only, not applied as corrections.*

For $R_2$*, we investigated cGM and dGM. For cGM, we used the lower part of Callaghan's reported significant $R_2$* slope range, approximately 0.03–0.06 $s^{-1}$/year; and for dGM, we used a basal-ganglia-relevant range of approximately 0.08–0.22 $s^{-1}$/year, reflecting the stronger effects reported in putamen, pallidum, and caudate. For temperature, we used the corresponding unfixed tissue $T_2$*-temperature equations: 0.2 [ms/°C] T + 31.9 ms for dGM (based on basal ganglia), 0.2 [ms/°C] T + 39.3 ms for cGM and 0.1 [ms/°C] T + 36.5 ms for WM.

| Tissue | % $R_2$* in Fig 3 | % $R_2$* due to temp. | % $R_2$* due to age |
|---|---|---|---|
| dGM | 16.28% | 8.264% | 13-35% |

| | | | |
|---|---|---|---|
| cGM | -2.22% | 6.865% | 6-12% |
| WM | 13.97% | 3.876% | No reliably reported |

*Table S9: Same for $R_2$* with Callaghan slope ranges and Birkl $T_2$*→$R_2$* conversion; bounds vs observed descriptive %.*

For $R_1$, cooling predicts an increase, while aging predicts a decrease at least in WM. This opposition makes the observed small WM R1 increase plausible, but cGM remains inconsistent because temperature would predict an increase whereas our data shows a slight decrease. For $R_2$*, cooling and aging both predict increases in dGM, so they could plausibly contribute to the observed +16.28% dGM increase. However, in cGM, both temperature and aging would predict an increase, opposite to what it is shown in our results. In WM, Callaghan does not support a simple broad age-related $R_2$* increase, so the observed +13.97% WM increase likely reflects other *postmortem* factors as well.

2. During the fixation process from day 0 to day 90 in formaldehyde immersion

In this transition, we observed $R_1$ increase exceeding up to 319.22%. To estimate the largest plausible temperature contribution, we considered an extreme cooling scenario from room temperature to approximately 4 °C ($\Delta T = -18$ °C). For that, we used the fixed-tissue equations from Birkl et al. (2016): for $T_1$, 3.1 [ms/°C] T + 285.6 ms for dGM (based on basal ganglia), 4.1 [ms/°C] T + 232.7 ms for cGM, and 1.1 [ms/°C] T + 230.6 ms for WM. For $T_2$*, 0.3 [ms/°C] T + 17.9 ms for dGM (based on basal ganglia), 0.2 [ms/°C] T + 26 ms for cGM and 0.1 [ms/°C] T + 24.9 ms for WM.

| Tissue | % $R_1$ in Fig 3 | % $R_1$ due to temp[(i)] | % $R_2$* in Fig 3 | % $R_2$* due to temp[(i)] |
|---|---|---|---|---|
| dGM | 319.22% | 18.725% | 22.54% | 28.272% |
| cGM | 245.59% | 29.6267% | 26.18% | 13.433% |
| WM | 255.55% | 8.426% | 45.89% | 7.115% |

*Table S10: Extreme fixation cooling (22°C to 4°C) using Birkl 2016 fixed-tissue equations vs observed $R_1$ and $R_2$* values from Figure 3. (i) Temperature difference of 18°C was used to estimate these changes.*

Predicted cooling effects during fixation were much smaller than the changes observed in the longitudinal $R_1$ trajectory, but not for $R_2$*. To explain the reported changes in $R_1$ during fixation (% $R_1$ in Figure 3, Table S10), the corresponding tissue classes would need to decrease their temperature to -33.97°C in cGM, -64.91°C in dGM and -144.49°C in WM. For $R_2$*, the possible temperature contribution varies by tissue and does not explain the overall pattern across them.

#### S4.5 Comparison across *postmortem* phases and fixation modeling by using only Brain1, Brain3 and Brain6.

To assess the influence of incomplete sampling, we repeated Analyses 1 and 2 after excluding Brain2, which had a short formaldehyde sampling period and no PBS scan, and Brain5, which had no *in situ* scan. The sensitivity analysis therefore included only Brain1, Brain3, and Brain6. We compared the resulting patterns with those in Figures 3 and 4. The final objective is to check how much our main results would differ when excluding these specimens.

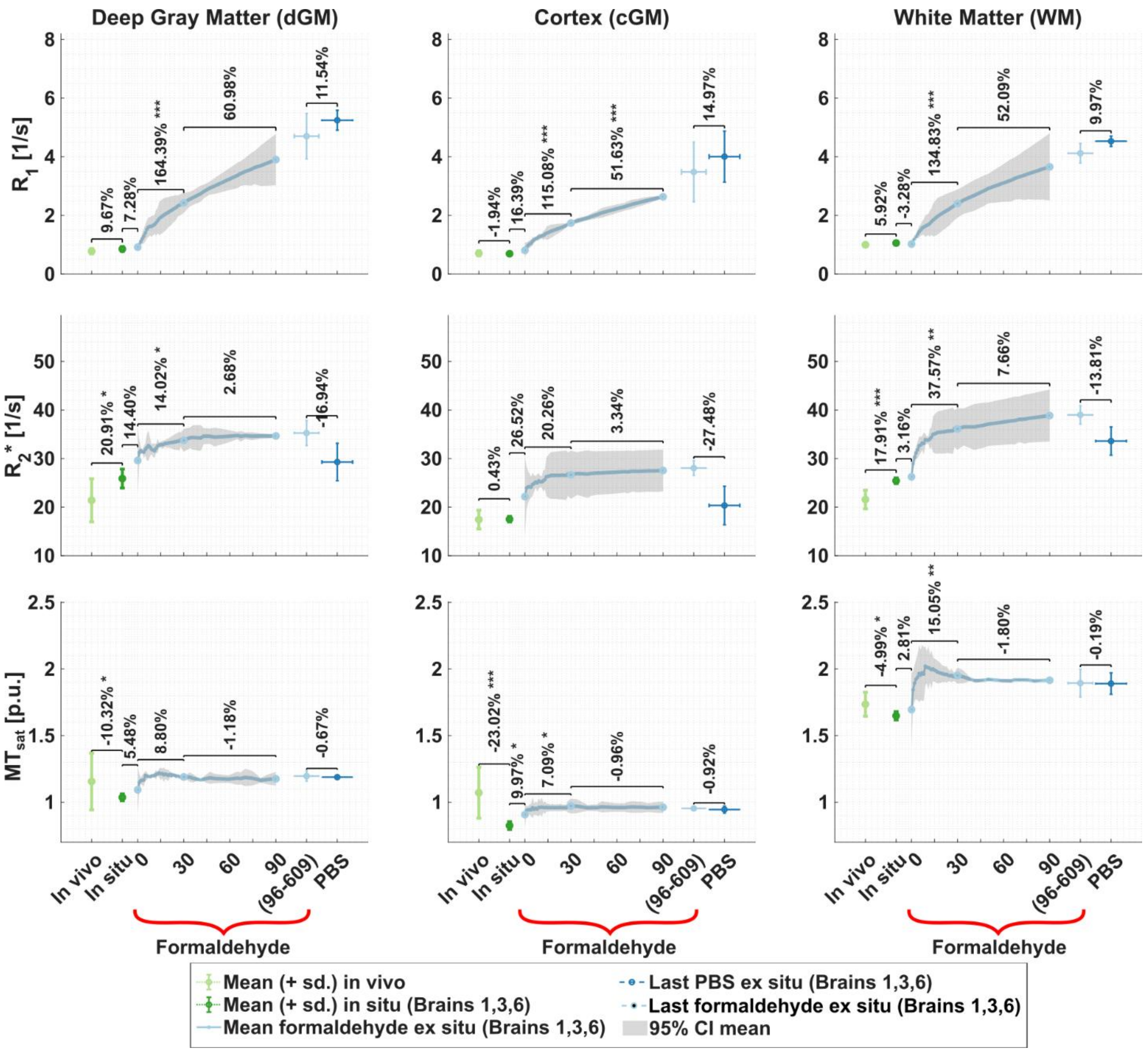


*Figure S15: Evolution of the MPM parameters ($R_1$, $R_2$* and $MT_{sat}$) across postmortem and fixation processes for only complete datasets: Brain1, Brain3 and Brain6. Summary markers show mean and standard deviation across specimen medians at different times and postmortem and fixation processes per tissue class (columns from left to right: deep gray matter, cortical gray matter, and white matter). Pairwise percentages compare those mean medians between consecutive postmortem processes in percentage: in vivo to in situ, in situ to formaldehyde day 0, formaldehyde day 0 to day 30, and formaldehyde day 30 to day 90. The final comparison is between each specimen's last formaldehyde and last PBS ex situ scan. Star symbols denote the magnitude of the descriptive p-value (**not statistical significance**): * reflects p-value < 0.05; ** reflects p-value < 0.01 and *** reflects p-value < 0.001. The last PBS ex situ scan was acquired when the specimen was, at least, day 7 in PBS + 0.1% sodium azide. The shaded curve during the fixation time represents the 95% confidence interval (CI) across specimens and the last formaldehyde scan session across specimens and time (between 93 to 603 days) is displayed by the x-y error bar and used to compare with the last PBS ex situ scan session.*

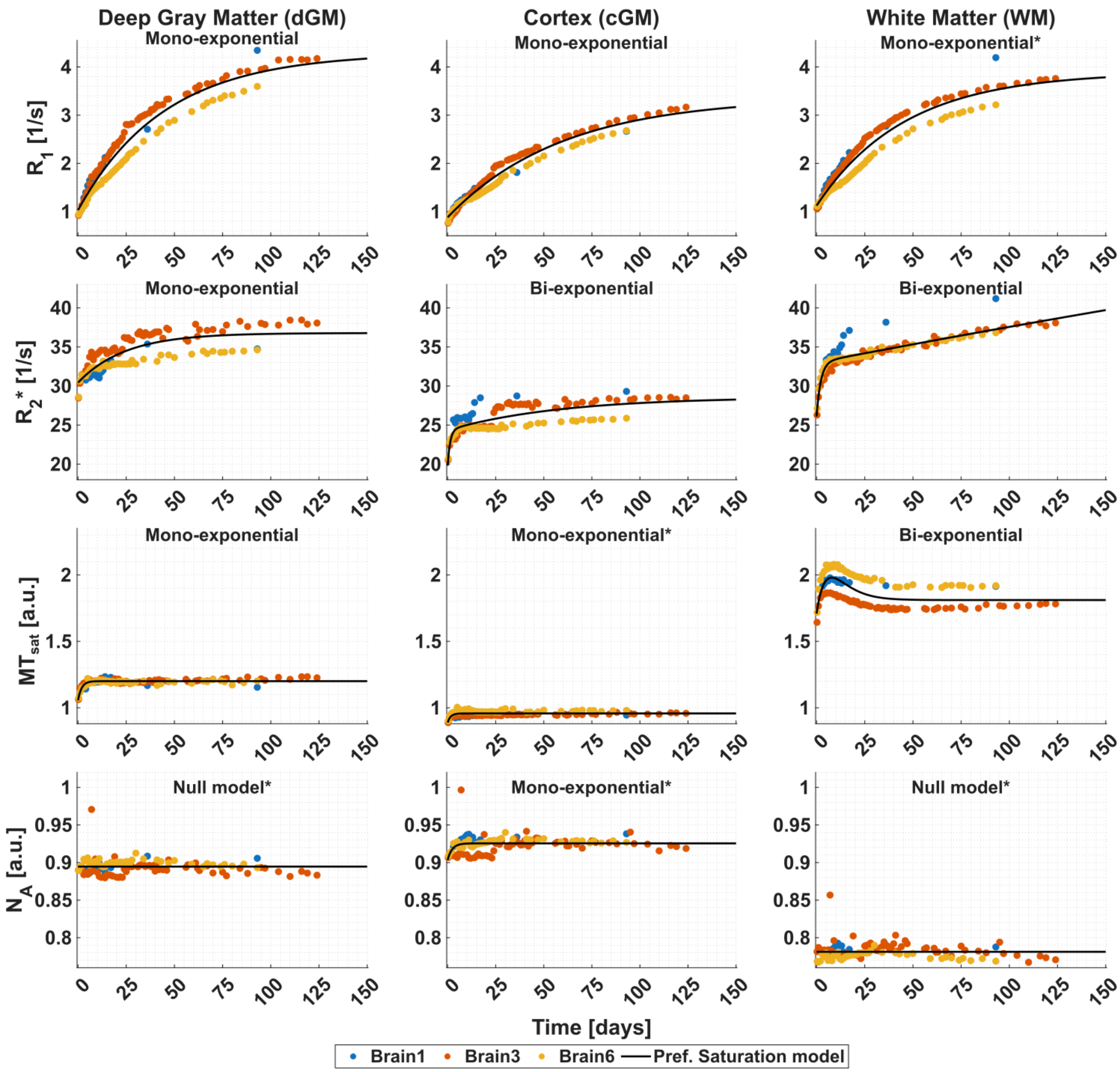


*Figure S16: Temporal evolution of the MPM parameters during fixation. For each tissue class (columns, left to right: dGM, cGM and WM), the median value of the MPM parameters is plotted (rows, top to bottom: $R_1$, $R_2$*, $MT_{sat}$ and $N_A$) for each brain specimen (colored markers) and per time point. The curve of the preferred saturation model ($\langle R_{ev}\rangle(t)$) is plotted for each tissue class and MPM parameter (black solid line) and its name shown on top of each subplot.*

In the complete-case subset (Brain1, Brain3, and Brain6), the qualitative pattern of MPM changes across *postmortem* conditions remained consistent with the full-cohort analysis in Figure 3 (main manuscript). R1 and R2* continued to show the largest fixation-related increases, $MT_{sat}$ remained comparatively stable after the early fixation period, and the last-formaldehyde to last-PBS contrast retained the same directional pattern (R1 increase, R2* decrease, small $MT_{sat}$ change; Figure S15). Because only three specimens enter this comparison, we treat the star symbols and exact p-values in Figure S15 as descriptive indices only, in line with Methods §2.3.

The preferred saturation models fitted to the complete-case trajectories (Figure S16) were the same as in the full cohort (Results §3.2 and §S3.1). Three exceptions were cGM-$R_1$ (from bi- to mono-exponential), cGM-$N_A$ (from null to mono-exponential) and WM-$N_A$ (from bi-exponential to null). The change in cGM-$R_1$ reflects the influence of earlier time-points from Brain2, while the changes in $N_A$ are consistent with weaker temporal structure and with choosing the simpler model form when model support is comparable.

Altogether, these checks suggest that the main descriptive conclusions of Analyses 1 and 2 are not driven solely by inclusion of Brain2 (short formaldehyde series; no PBS) or Brain5 (no *in situ* scan). They do not increase the effective sample size for inference; the complete-case analysis remains a sensitivity check under a reduced cohort.

#### S4.6 Relationship between *postmortem* interval and MPM parameters

We investigated the relationship between *postmortem* interval (PMI) against the median MPM parameters per tissue class, for the first formaldehyde *ex situ* scan session in Figure S17. The PMI for the formaldehyde *ex situ* scan is reported in Table 1.

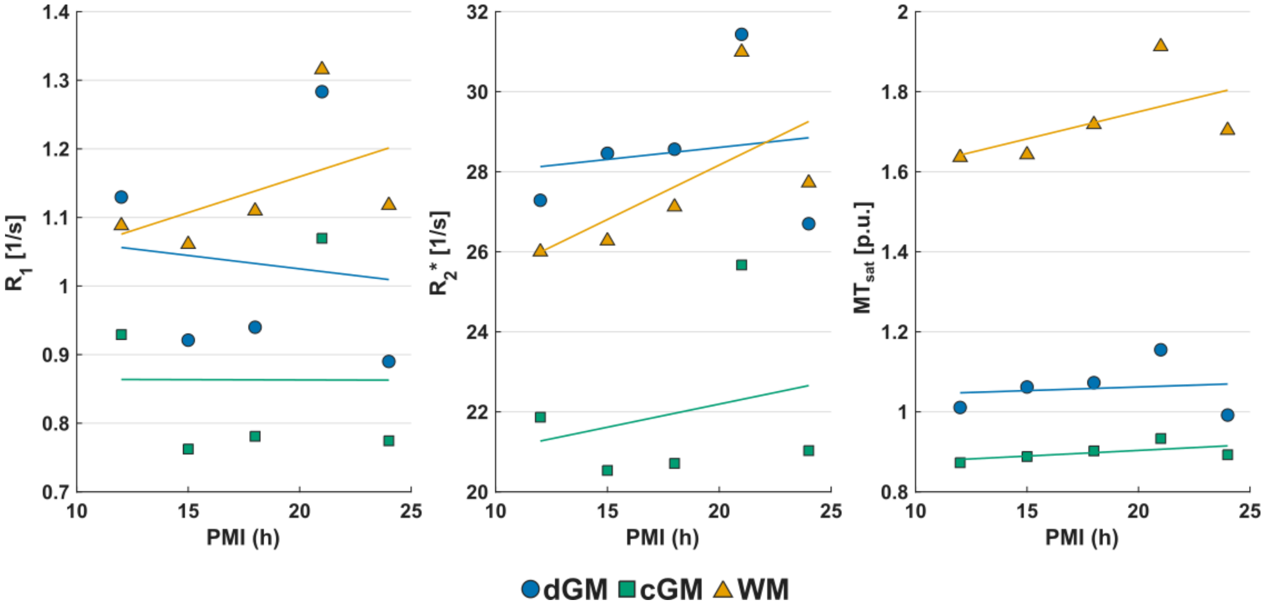


| MPM parameter | Tissue class | OLS model | $R^2$ |
|---|---|---|---|
| R1 | dGM | $y = -0.0164x + 1.25$ | 0.60 |
| | cGM | $y = -0.00991x + 0.983$ | 0.42 |
| | WM | $y = 0.00353x + 1.03$ | 0.51 |
| R2* | dGM | $y = -0.0716x + 29$ | 0.16 |
| | cGM | $y = -0.0445x + 21.8$ | 0.15 |
| | WM | $y = 0.151x + 24.2$ | 0.95 |
| MTsat | dGM | $y = -0.0025x + 1.08$ | 0.11 |
| | cGM | $y = 0.00154x + 0.862$ | 0.41 |
| | WM | $y = 0.00639x + 1.56$ | 0.61 |

*Figure S17: Tissue-class median $R_1$, $R_2$*, and $MT_{sat}$ at the first formaldehyde ex situ scan versus PMI (time from death to start of fixation) for four specimens (Brain1 excluded because its first formaldehyde session already reflected several days in fixative). Markers are colored by tissue class (dGM, cGM, WM). Ordinary least-squares lines and inserted in Figure, while slope, intercept, and $R^2$ values are given in table. Note that these values and the drawn linear fits are reported for visual guidance only and are not confirmatory.*

We found there is a slight increase in WM-$R_2$* and WM-$MT_{sat}$ with respect to PMI in formaldehyde *ex situ* condition, while the remaining tissue classes and MPM parameters do not show a clear trend, e.g., dGM- and cGM-$R_1$ and $R_2$*. Even more, WM-$R_2$* reported the highest $R^2$

(marked in red in Table), showing that a possible dependency could exist between PMI and $R_2^*$. However, to re-iterate, It is important to mention that, even with this finding, it is not appropriate to interpret any statistical significance of these results, given the low number of specimens (n = 4).

### S5. More about shrinkage

This section reports additional analyses that were performed on the relative volume change. Here, we are looking at two different analyses: another exploratory correlation between MPM temporal change and relative volume change (§S5.1); and relative volume change during formaldehyde and PBS *ex situ* scan sessions (§S5.2).

#### S5.1. Comparison between the inverse of rV(t) and MPM temporal change

We also investigated the comparison between MPM parameters and the relative volume change using a mechanistic-inspired relationship inspired a model proposed by (Fatouros et al., 1991) and evaluated by, e.g., (Gelman et al., 2001). Fatouros' model says that $R_1$ is related to water content, W, both bounded and free, by the following relationship:

$$R_1 = \kappa\left(R_{1b} - R_{1f}\right)\left(\frac{1}{W} - 1\right) + R_{1f} \quad \text{(S6)}$$

where $R_{1b}$ and $R_{1f}$ are the corresponding $R_1$ values for the bound and free water pools respectively; and κ is the hydration fraction, defined as the ratio of bound water to solid tissue component. Equation S6 can be rewritten as $R_1 = A + B/W$, where the slope B is defined by $B = \kappa(R_{1b} - R_{1f})$ and the intercept A is defined as $R_{1f} - B$.

In the same study, the authors assumed that $R_{1f} << R_{1b}$, which makes the slope and intercepts be approximately equal to $B = \kappa R_{1b}$ and $A = -B$; or:

$$R_1 = \kappa R_{1b}\left(\frac{1}{W} - 1\right) \quad \text{(S7)}$$

In our study, we hypothesized that fixation-related volume changes are mainly caused by water loss, and thus used 1/rV as a proxy for 1/W. Here, we tested whether this model correlates with $R_1$ (as originally defined) and extend it for exploratory purposes to $R_2^*$ and $MT_{sat}$ for which a volume change was observed in our study (see Figure 5). The results are depicted in Figure S18.

Note that equation S7 is the Fatouros water-content form and should not be confused with the iron–myelin construction in §S7 (Equations S8–S11).

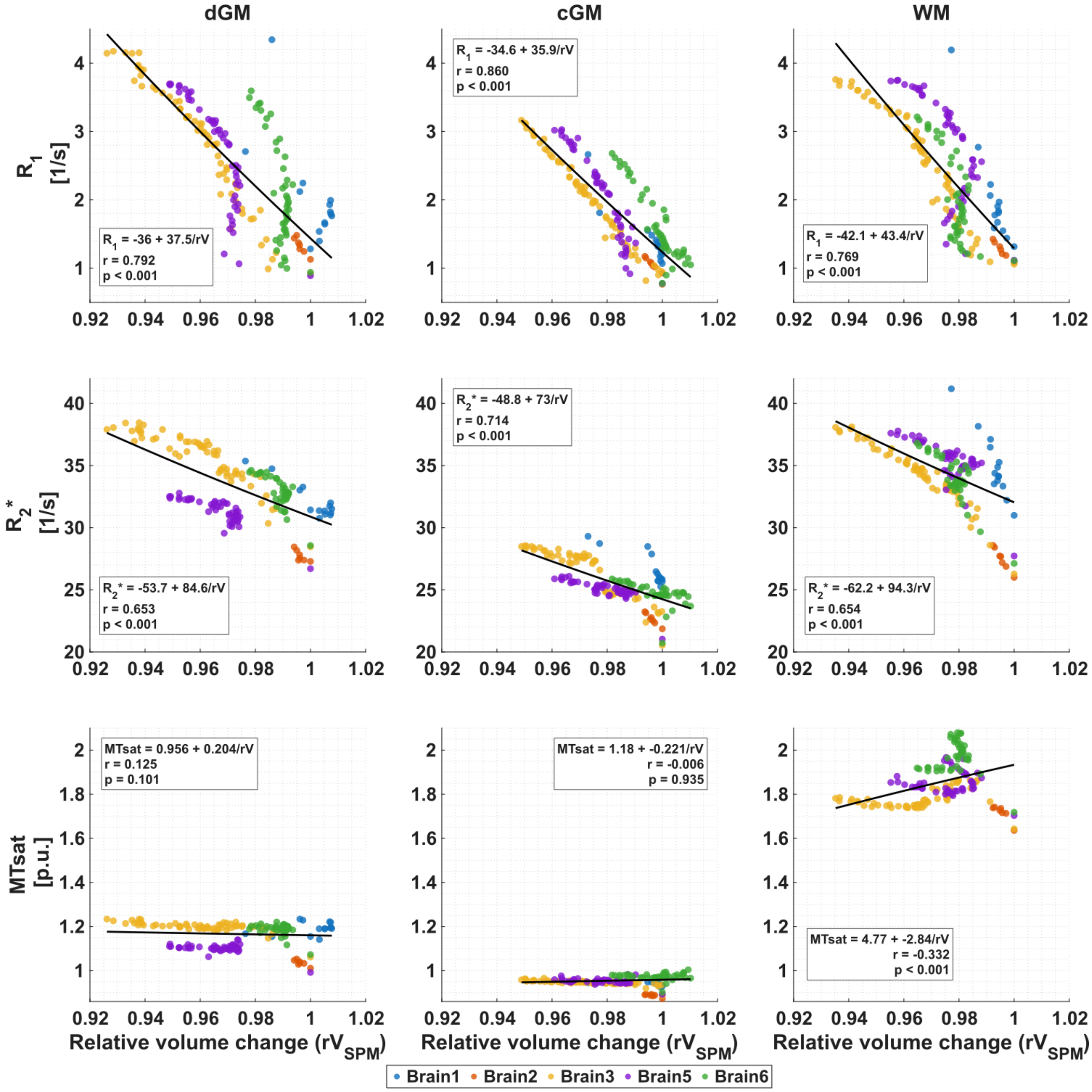


*Figure S18: Linear fits between the MPM parameters $R_1$, $R_2$*, and $MT_{sat}$ and the inverse of the relative volume change model ($1/rV(t)$) for each tissue class and across all specimens (color-coded dots). The line overlaid in each subplot (black line) is the fitted linear regression, whose equations are as follows:* $R_1 = A_{R_1} + \frac{B_{R_1}}{rV}$ *(top row),* $(R_2)^* = A_{R_2^*} + \frac{B_{R_2^*}}{rV}$ *(middle row) and* $MT_{sat} = A_{MT_{sat}} + \frac{B_{MT_{sat}}}{rV}$ *(bottom row).*

We observed that the fitted parameters between $R_1$ and 1/rV resembled Equation S7, i.e., the slope and intercept are similar, for all tissue classes. Particularly for WM, our fitted value for slope B was approximately 43 $s^{-1}$ while intercept A was approximately -42 $s^{-1}$, whereas in Fatouros et al. the corresponding values are about one order of magnitude smaller. This discrepancy could have three explanations: (1) that Fatouros's and Gelman's parameters were obtained from *in vivo* cat and human brain, (2) 1/rV is not equivalent to 1/W, and (3) Fatouros coefficient depends on the hydration fraction and relaxation properties of motionally restricted water ($R_{1b}$ and $R_{1f}$, and κ). Both are likely altered during formaldehyde fixation through protein cross-linking, changes in hydration layers and altered water–macromolecule exchange. The substantially larger $R_1$ response relative to the small macroscopic volume change therefore suggests that fixation-associated water loss or redistribution alone cannot explain the observed $R_1$ increase but likely acts together with fixation-induced changes in tissue relaxivity.

| Parameter | Fitted results in WM (Figure S18) | Fatouros et al. (i) parameters | Gelman et al. (ii) parameters |
|---|---|---|---|
| Slope (B, $s^{-1}$) | 43.357 ± 2.903 | 2.7 | 1.99 ± 0.06 |
| Intercept (A, $s^{-1}$) | -42.067 ± 2.977 | -2.7 | −1.75 ± 0.07 |
| $R_1$-weighted hydration fraction ($\kappa R_{1b}$) | 43.357 ± 2.903 | 2.7 | 1.99 ± 0.06 |

*Table S11: Parameters of the fitted model in Equations S6 and S7 for WM, compared with experimental values from previous studies ((i) Fatouros et al. (1991) and (ii) Gelman et al. (2001)), where slope (B) is defined as the R1-weighted hydrated fraction ($\kappa R_{1b}$), and intercept (A) defined as minus $\kappa R_{1b}$. The $\kappa R_{1b}$ was estimated based on the value of B rather than A. Whereas Gelman's parameters were obtained by fitting not only WM but also cGM and some dGM structures, Fatouros's parameters were obtained by extrapolating $\kappa R_{1b}$ from Figure 1 in their study. Standard error of each fitted parameter is given after the ± in our study and in Gelman's parameters.*

The fitted parameters for the other MPM didn't resemble the relationship observed for $R_1$, which indicated that this model is not sufficient to explain the relationship between water volume and MPM.

#### S5.2. Relative volume change during formaldehyde and PBS *ex situ* scan sessions

Figure S19 illustrates only the transition of mean relative volume change, rV(t), across formaldehyde and the latest PBS *ex situ* scan sessions (x-axis) per tissue class (columns). These rV(t) were obtained by averaging the rV(t) means across specimens per tissue class (Figure 5 in Results §3.3).

We observed that relative volume decreased by an additional 1.62% in dGM, 1.32% in cGM, and 1.90% in WM, between the last formaldehyde and latest PBS *ex situ* scans. This exploratory PBS-stage and last formaldehyde *ex situ* scan comparison describe changes under our experimental protocol rather than a fully equilibrated rehydration state. Therefore, further studies with a standardized PBS wash-out protocol must be performed, i.e., standardized PBS immersion duration, renewal of the solution.

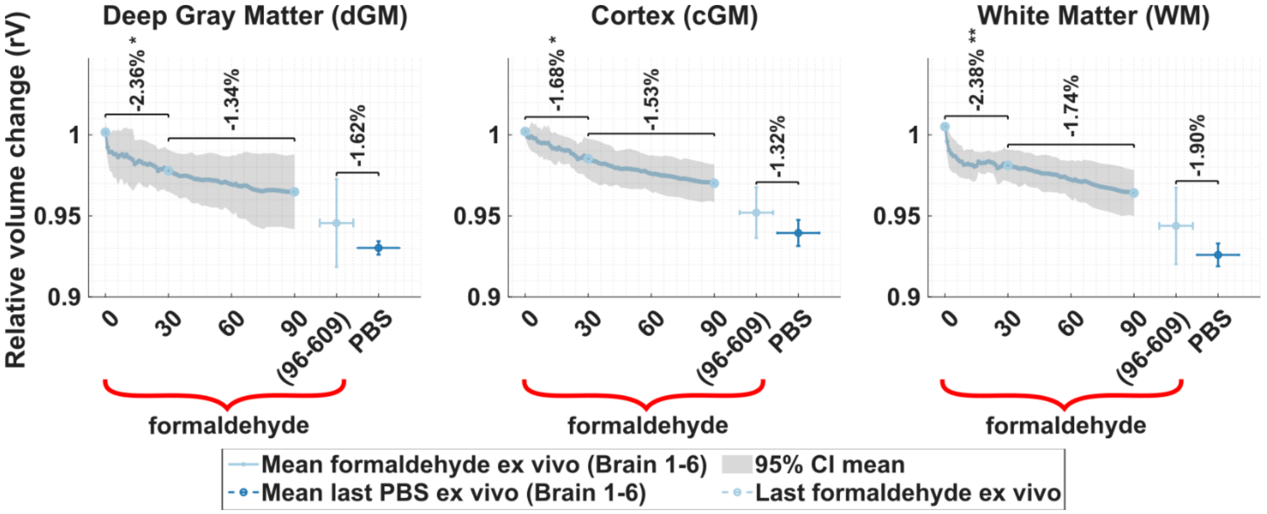


*Figure S19: Evolution of the relative volume change, rV(t), across formaldehyde and PBS ex situ scan sessions for all specimens. Depicted are the following metrics: the mean and the standard deviation at different times and the formaldehyde ex situ scan sessions, denoted as formaldehyde; and the last PBS ex situ scan, denoted by PBS; per tissue class (columns from left to right: deep gray matter, cortical gray matter, and white matter), and the difference between two consecutive postmortem processes in percentage: formaldehyde ex situ scan sessions at day 30 – day 0, at day 90 – day 30, and between last formaldehyde and last PBS ex situ scan sessions. The last PBS ex situ scan was acquired when the specimen was, at least, day 7 in PBS + 0.1% sodium azide. The shaded curve during the fixation time represents the 95% confidence interval (CI) across specimens and the last formaldehyde scan session across specimens and time (between 93 to 603 days) is displayed by the x-y error bar and used to compare with the last PBS ex situ scan session.*

### S6. Information about the videos

We created three videos for each of the five *postmortem* brain specimens: (1) longitudinally deformed MPM maps across all time points (complementary to Results §3.1 and §3.2), (2) composite Jacobians across all formaldehyde and PBS *ex situ* scan sessions (complementary to Results §3.3), and (3) slice-scrolling MPM maps for the latest formaldehyde *ex situ* scan session across MPM parameters (complementary to Results §3.1 and §3.2). These videos are intended to support visual inspection and verification of spatial alignment/deformation, and MPM contrast appearance, given that the underlying image data are not available for the reader. The videos included in this publication were created using create_temporal_video_three_views.m, create_temporal_video_mtsat_jacobian.m and create_spatial_slice_video_lastPFA.m, respectively. For the first video, we used the longitudinally registered MPM images, i.e. $R_1$, $R_2$* and $MT_{sat}$, with ANTs, after bubble masking and brain masking based on the estimated tissue probability maps for the first timepoint per specimen, i.e. cGM, WM, dGM and cerebellum. The second video shows the composite Jacobians (from §S1, "Jacobian Compositions") for all formaldehyde and PBS *ex situ* scan sessions relative to the first formaldehyde *ex situ* scan session. For the third video, we used the last longitudinally deformed formaldehyde *ex situ* scan session and scrolled through slices for all MPM images; these maps were likewise bubble- and brain-masked. Slice positions and image rotations used for visualization are summarized in Table S12.

| Specimen | First video (Long. Deformation registration check) | Second video (Jacobian) | Third video (spatial HWM check) |
|---|---|---|---|
| Brain1 | [140,**160**,112] | [87,89,80] | Through coronal |
| Brain2 | [140,**170**,112] | [74,114,111] | Through coronal |

| | | | |
|---|---|---|---|
| Brain3 | [140,**180**,112] | [94,89,71] | Through coronal |
| Brain5 | [160,140,**87 (90°)**] | [104,95,74] | Through transversal |
| Brain6 | [160,140,**127 (90°)**] | [105,97,81] | Through transversal |

*Table S12: Slice indices and display orientations used in the three supplementary videos. Voxel indices for the first and second videos are given as **[sagittal, coronal, transversal]** (1-based) in the corresponding image space of each video. For Brain5 and Brain6 in the first video, the indicated 90° rotation applies only to the locked transversal plane; the companion sagittal and coronal views are shown without rotation. The third video scrolls through consecutive non-zero (brain-masked) slices along a single anatomical axis (coronal or transversal). These settings were used solely for visualization to support qualitative inspection of alignment, deformation, and MPM contrast.*

### S7. Link between saturation-relaxation and microstructural relaxivities models

In the work of (Callaghan et al., 2015; Kirilina et al., 2020; Rooney et al., 2007; Stüber et al., 2014), both $R_1$ and $R_2$* showed an empirical linear dependence on myelin content, iron content and the “medium” in which the specimen is embedded. This relationship can be expressed as:

$$R = M_{Fe} r_{Fe} + M_{Mye} r_{Mye} + r_{env}, \qquad \text{(S8)}$$

where $M_{Fe}$ is the iron concentration, $M_{Mye}$ is the myelin volume fraction, $r_{Fe}$ and $r_{Mye}$ are the relaxivities of iron and myelin, respectively; $r_{env}$ is the relaxivity of the surrounding medium: While *in vivo* $r_{env}$ is the relaxivity of free water (Callaghan et al., 2015), here the surrounding medium will contain the formaldehyde reaction products from the fixative solution.

During fixation, chemical processes such as dehydration, iron oxidation and the cross-linking of the formaldehyde solution alter $r_{Fe}$ and $r_{Mye}$. In this scenario, Equation S8 can be re-written as a function of fixation time t:

$$R_{fix}(t) = M_{Fe} r_{Fe}(t) + M_{Mye} r_{Mye}(t) + r_{env}(\mathrm{t}) \qquad \text{(S9)}$$

To assess how each component changes during fixation, $R_{fix}(t)$ is referenced to t = 0 on the same formaldehyde-ex-situ clock as Methods §2.4. That intercept is the fitted $R_{ev,0}$ and it is not assumed to be chemically unaltered ($R_{ev,0} = R_{fix,0} = R_{fix}(0)$). Subtracting this baseline from Equation S9 gives:

$$R_{fix}(t) = R_{ev,0}(0) + s(t), \qquad \text{(S10)}$$

with:

$$s(t) = M_{Fe}\big(r_{Fe}(t) - r_{Fe}(0)\big) + M_{Mye}\left(r_{Mye}(t) - r_{Mye}(0)\right). \qquad \text{(S11)}$$

Here s(t) is the change in relaxivity-weighted contributions relative to day 0, under the assumptions $r_{env}(\mathrm{t}) = r_{env}(0)$ and constant $M_{Fe}$ and $M_{Mye}$. Equations 2 and 3 share the additive structure of Equation S10, $R_{ev}(t) = R_{ev,0}(0) + s(t)$; they do not identify the terms in Equation S11. A mono- or bi-exponential s(t) can be read as one or two effective temporal components, which might coincide with iron- or myelin-related relaxivity change, but that assignment is not constrained by the present data. If $r_{env}$ also varies (for example through formaldehyde uptake), or if $M_{Fe}$ or $M_{Mye}$ change with shrinkage, further terms enter s(t) and the mapping is no longer one-to-one. This construction applies to R1 and R2*; MTsat and NA are not covered by Equations S8–S11.

### S8. References


Aho, K., Derryberry, D., Peterson, T., 2017. A graphical framework for model selection criteria and significance tests: refutation, confirmation and ecology. Methods in Ecology and Evolution 8, 47–56. https://doi.org/10.1111/2041-210X.12648

Alexander, A.L., McCreery, T.T., Barrette, T.R., Gmitro, A.F., Unger, E.C., 1996. Microbubbles as novel pressure-sensitive MR contrast agents. Magnetic Resonance in Medicine 35, 801–806. https://doi.org/https://doi.org/10.1002/mrm.1910350603

Ashburner, J., Ridgway, G.R., 2013. Symmetric Diffeomorphic Modeling of Longitudinal Structural MRI. Frontiers in Neuroscience Volume 6-2012. https://doi.org/10.3389/fnins.2012.00197

Bazin, P.-L., Alkemade, A., van der Zwaag, W., Caan, M., Mulder, M., Forstmann, B.U., 2019. Denoising High-Field Multi-Dimensional MRI With Local Complex PCA. Frontiers in Neuroscience Volume 13-2019. https://doi.org/10.3389/fnins.2019.01066

Billot, B., Greve, D.N., Puonti, O., Thielscher, A., Van Leemput, K., Fischl, B., Dalca, A.V., Iglesias, J.E., 2023. SynthSeg: Segmentation of brain MRI scans of any contrast and resolution without retraining. Medical Image Analysis 86, 102789. https://doi.org/10.1016/j.media.2023.102789

Birkl, C., Langkammer, C., Golob-Schwarzl, N., Leoni, M., Haybaeck, J., Goessler, W., Fazekas, F., Ropele, S., 2016. Effects of formalin fixation and temperature on MR relaxation times in the human brain. NMR in Biomedicine 29, 458–465. https://doi.org/https://doi.org/10.1002/nbm.3477

Birkl, C., Langkammer, C., Haybaeck, J., Ernst, C., Stollberger, R., Fazekas, F., Ropele, S., 2014. Temperature-induced changes of magnetic resonance relaxation times in the human brain: A postmortem study. Magnetic Resonance in Medicine 71, 1575–1580. https://doi.org/10.1002/mrm.24799

Burnham, K.P., Anderson, D.R., 2004. Multimodel Inference: Understanding AIC and BIC in Model Selection. Sociological Methods & Research 33, 261–304. https://doi.org/10.1177/0049124104268644

Burnham, K.P., Anderson, D.R., 2002. Model selection and multimodel inference: a practical information-theoretic approach. Springer Verlag.

Burnham, K.P., Anderson, D.R., Huyvaert, K.P., 2011. AIC model selection and multimodel inference in behavioral ecology: some background, observations, and comparisons. Behavioral Ecology and Sociobiology 65, 23–35. https://doi.org/10.1007/s00265-010-1029-6

Callaghan, M.F., Freund, P., Draganski, B., Anderson, E., Cappelletti, M., Chowdhury, R., Diedrichsen, J., FitzGerald, T.H.B., Smittenaar, P., Helms, G., Lutti, A., Weiskopf, N., 2014. Widespread age-related differences in the human brain microstructure revealed by quantitative magnetic resonance imaging. Neurobiology of Aging 35, 1862–1872. https://doi.org/10.1016/j.neurobiolaging.2014.02.008

Callaghan, M.F., Helms, G., Lutti, A., Mohammadi, S., Weiskopf, N., 2015. A general linear relaxometry model of R1 using imaging data. Magnetic resonance imaging 73, 1309–14.

Edwards, L., Bazin, P.L., Tabelow, K., Ugurcan, B.E., Mohammadi, S., Weiskopf, N., 2024. Denoising improves contrast while retaining sharpness of high resolution multiparameter R1, R2* and proton density maps. Magnetic Resonance Materials in Physics, Biology and Medicine 37, 1–781. https://doi.org/10.1007/s10334-024-01191-6

Fatouros, P.P., Marmarou, A., Kraft, K.A., Inao, S., Schwarz, F.P., 1991. In Vivo Brain Water Determination by T1 Measurements: Effect of Total Water Content, Hydration Fraction, and Field Strength. Magnetic Resonance in Medicine 17, 402–413. https://doi.org/10.1002/mrm.1910170212

Fedorov, A., Beichel, R., Kalpathy-Cramer, J., Finet, J., Fillion-Robin, J.-C., Pujol, S., Bauer, C., Jennings, D., Fennessy, F., Sonka, M., Buatti, J., Aylward, S., Miller, J.V., Pieper, S., Kikinis, R., 2012. 3D Slicer as an image computing platform for the Quantitative Imaging Network. Magnetic Resonance Imaging 30, 1323–1341. https://doi.org/https://doi.org/10.1016/j.mri.2012.05.001

Gelman, N., Ewing, J.R., Gorell, J.M., Spickler, E.M., Solomon, E.G., 2001. Interregional variation of longitudinal relaxation rates in human brain at 3.0 T: Relation to estimated iron and water contents. Magnetic Resonance in Medicine 45, 71–79. https://doi.org/10.1002/1522-2594(200101)45:1%3C71::AID-MRM1011%3E3.0.CO;2-2

Ghosh, J.K., Delampady, M., Samanta, T., 2007. An Introduction to Bayesian Analysis: Theory and Methods, Springer Texts in Statistics. Springer New York.

Kirilina, E., Helbling, S., Morawski, M., Pine, K., Reimann, K., Jankuhn, S., Dinse, J., Deistung, A., Reichenbach, J.R., Trampel, R., Geyer, S., Müller, L., Jakubowski, N., Arendt, T., Bazin, P.-L., Weiskopf, N., 2020. Superficial white matter imaging: Contrast mechanisms and whole-brain in vivo mapping. Science Advances 6, eaaz9281. https://doi.org/10.1126/sciadv.aaz9281

Molinaro, A.M., Simon, R., Pfeiffer, R.M., 2005. Prediction error estimation: a comparison of resampling methods. Bioinformatics 21, 3301–3307. https://doi.org/10.1093/bioinformatics/bti499

Otsu, N., 1979. A Threshold Selection Method from Gray-Level Histograms. IEEE Transactions on Systems, Man, and Cybernetics 9, 62–66. https://doi.org/10.1109/TSMC.1979.4310076

Rooney, W.D., Johnson, G., Li, X., Cohen, E.R., Kim, S.-G., Ugurbil, K., Springer Jr., C.S., 2007. Magnetic field and tissue dependencies of human brain longitudinal 1H2O relaxation in vivo. Magnetic Resonance in Medicine 57, 308–318. https://doi.org/https://doi.org/10.1002/mrm.21122

Stüber, C., Morawski, M., Schäfer, A., Labadie, C., Wähnert, M., Leuze, C., Streicher, M., Barapatre, N., Reimann, K., Geyer, S., Spemann, D., Turner, R., 2014. Myelin and iron concentration in the human brain: A quantitative study of MRI contrast. NeuroImage 93, 95–106. https://doi.org/10.1016/j.neuroimage.2014.02.026

Tabelow, K., Balteau, E., Ashburner, J., Callaghan, M.F., Draganski, B., Helms, G., Kherif, F., Leutritz, T., Lutti, A., Phillips, C., Reimer, E., Ruthotto, L., Seif, M., Weiskopf, N., Ziegler, G., Mohammadi, S., 2019. hMRI – A toolbox for quantitative MRI in neuroscience and clinical research. NeuroImage 194, 191–210. https://doi.org/https://doi.org/10.1016/j.neuroimage.2019.01.029

Thomas, D.C., Deichmann, R., Nöth, U., Langkammer, C., Ferreira, M., Golbach, R., Hattingen, E., Wenger, K.J., 2024. A fast protocol for multicenter and multiparametric quantitative MRI studies in brain tumor patients using vendor sequences. Neuro Oncol Adv 6, vdae117. https://doi.org/10.1093/noajnl/vdae117

Tredennick, A.T., Hooker, G., Ellner, S.P., Adler, P.B., 2021. A practical guide to selecting models for exploration, inference, and prediction in ecology. Ecology 102, e03336. https://doi.org/10.1002/ecy.3336

Weiskopf, N., Suckling, J., Williams, G., Correia, M.M., Inkster, B., Tait, R., Ooi, C., Bullmore, E.T., Lutti, A., 2013. Quantitative multi-parameter mapping of R1, PD(*), MT, and R2(*) at 3T: a multi-center validation. Front Neurosci 7, 95.

Yates, L.A., Aandahl, Z., Richards, S.A., Brook, B.W., 2023. Cross validation for model selection: A review with examples from ecology. Ecological Monographs 93, e1557. https://doi.org/10.1002/ecm.1557
